# Ethnic differences in gut microbiota composition and functional capacity persist over six years in the HELIUS multi-ethnic longitudinal cohort

**DOI:** 10.64898/2026.10.12.757716

**Authors:** B.J.H. Verhaar, U. Boulund, H. Herrema, M. Nicolaou, A.H. Zwinderman, B.J.H. van den Born, N. Hamdiui, E. Elinav, T. Nielsen, M. Nieuwdorp

## Abstract

**Background:** Ethnic minority groups in Western Europe carry a disproportionate cardiometabolic disease burden not fully explained by conventional risk factors, and their gut microbiomes differ markedly from those of European-origin populations. Whether these differences persist over time has not been examined prospectively.

**Results:** We followed 1,903 participants from six ethnic groups in the HELIUS cohort over 6 years, using faecal 16S rRNA sequencing across all groups and shotgun metagenomics in a matched Dutch and South-Asian Surinamese subset (N = 238 per group). The extent of microbiota compositional change over time differed between ethnic groups: Moroccan participants showed the largest change, while the Dutch and South-Asian Surinamese participants were most stable. Diabetes mellitus and medication use at baseline were associated with greater compositional drift. However, ethnic differences in composition were largely preserved across all six groups (16S: ethnicity R² 0.044 to 0.037; all 15 pairwise contrasts significant at follow-up) and in the shotgun subset (R² 0.047 to 0.037). At the species level, 61 of 78 ethnicity-associated species remained differentially abundant at follow-up. *Alistipes putredinis*, the species converging most strongly, was absent at baseline in 43% of South-Asian Surinamese participants, and the convergence was consistent with acquisition of this species by this group. At strain level, within-person strain sharing declined with follow-up duration but was similar between ethnic groups. Within hosts, 90% of *A. putredinis* populations were retained as the same or a closely related lineage, and within the retained subset, microdiversity increased in both groups. Ethnic differences in functional profiles persisted to a similar extent as those in composition, with limited convergence in fibre-degrading enzyme families (GH5_37, GH26). The total antibiotic resistance gene (ARG) burden in the gut microbiome declined similarly in both groups over time. Dutch participants showed higher baseline ARG diversity that converged toward South-Asian Surinamese levels, while class-level differences were predominantly stable.

**Conclusions:** Six years of follow-up in a shared urban environment narrowed ethnic differences in the gut microbiome only modestly; assuming exponential decline, the observed change corresponds to a half-life of roughly two decades.

## Introduction

Ethnic minority groups in Western Europe carry a disproportionate burden of cardiometabolic disease that persists after adjustment for socioeconomic factors and is not fully explained by conventional risk factors. In a UK cohort study of South Asian and African Caribbean participants, a 3-to 4-fold excess of type 2 diabetes mellitus compared to Europeans was observed.^1^ A multi-site study of Ghanaian migrants found that cardiovascular disease (CVD) and diabetes risk was substantially higher in Amsterdam, Berlin, and London than in Ghana, suggesting that the Western urban environment itself contributes to this excess.^2,3^

The gut microbiome is a plausible contributor to ethnic cardiometabolic disparities. In the Healthy Living in an Urban Setting (HELIUS) cohort, gut microbiome composition was shown to differ substantially between ethnic groups despite shared urban residence.^4^ In the same cohort, we previously found that specific baseline taxa were associated with incident cardiometabolic disease.^5^ Beyond antibiotic use, diet is the strongest known modifiable determinant of microbiome composition,^6,7^ and dietary patterns differ substantially between ethnic groups in HELIUS and persist independent of socioeconomic position.^8^ Other lifestyle factors such as BMI, physical exercise and medication further contribute to these differences,^9,10^ in addition to host genetics, although the latter’s contribution is modest.^7,11,12^

How ethnic differences in microbiome composition change over time in a population sharing the same Western urban environment, and what drives that change, remains unknown. Prior work comparing ethnic groups in a shared Western environment has been cross-sectional or confined to the period immediately after migration. Shared urban residence could drive gradual convergence in microbiome composition as dietary patterns and lifestyle factors partially align across generations.^13^ Alternatively, many differences rooted in cultural dietary practices, living conditions and host genetic background may persist.^11,14,15^ Microbiome stability itself may have clinical relevance: compositional instability has been associated with adverse health outcomes such as inflammatory bowel disease, steatotic liver disease and diabetes.^16,17^

Here, we used 6 years of follow-up in HELIUS to ask whether ethnic differences in gut microbiome composition are stable, convergent, or divergent over time, and what drives individual-level change. Using 16S rRNA sequencing across all six ethnic groups and shotgun metagenomics in a Dutch and South-Asian Surinamese subset, we characterised longitudinal dynamics at compositional, species, strain, functional, and resistome level, and tested associations with cardiometabolic risk factors and disease.

## Methods

### Cohort and participants

We used baseline and follow-up data from the HELIUS cohort, a prospective multi-ethnic population-based cohort study (N = 24,780) conducted in Amsterdam, the Netherlands, with baseline visits between 2011 and 2015.^18^ Participants were randomly sampled from the municipal population registry, stratified by ethnicity to ensure representation of six ethnic groups: Dutch, African Surinamese, South-Asian Surinamese, Ghanaian, Turkish, and Moroccan origin. Ethnic origin was assigned based on country of birth of the participant and both parents: non-Dutch origin was defined as born outside the Netherlands with at least one parent born outside the Netherlands, or born in the Netherlands with parents born outside the Netherlands.^19^ Participants of Surinamese origin were further classified by self-reported ethnicity as African-Surinamese or South-Asian Surinamese. Between May 2019 and November 2022, all baseline participants still living in the Netherlands were invited for a first follow-up visit.^20^ Of 24,780 baseline participants, 371 had died and 640 had moved abroad, leaving 23,769 eligible for follow-up; 11,035 (46.4%) attended. Non-response analyses comparing participants who did and did not attend follow-up, in the total cohort and the microbiome substudy, are reported elsewhere.^5,20^ Ethical approval was obtained from the medical ethical review board (Amsterdam UMC, location AMC), the study followed the principles of the Declaration of Helsinki, and all participants provided written informed consent. In reporting our analyses, we followed the STROBE guidelines (checklist in **Supplementary Table 5**).^21^

Data collection at both timepoints included a general questionnaire, medical history, dietary assessment, physical examination, and fasting blood sampling. Study visits took place in the morning after an overnight fast. Height and weight were measured and BMI was calculated. Blood pressure was measured three times using a semi-automatic oscillometric device (Microlife WatchBP Home; Microlife AG, Switzerland), and the mean of the last two readings was used. Fasting venous blood samples were collected, processed, and aliquoted; plasma samples were stored temporarily at 4–7°C at the local research site before transfer to central laboratories for storage at −80°C. Total cholesterol, HDL-C, and triglycerides were measured using a colorimetric reagent on an ABX Pentra 400 chemistry analyser (Horiba ABX SAS, Oberursel, Germany); LDL-C was calculated using the Friedewald equation. Participants were asked to bring their current medication to the study visit. For our analyses, we used statins (ATC C10AA), metformin (ATC A10BA02), glucose lowering drugs (ATC A10), blood pressure lowering drugs (ATC C02, C03, C07, C08, C09), corticosteroids (ATC H02) and psychotropic medication (at least one type of psychiatric medication, including medication without ATC classification). Hypertension was defined as a systolic blood pressure ≥140 mmHg, diastolic blood pressure ≥90 mmHg, or use of antihypertensive medication. Metabolic syndrome was defined according to the IDF/AHA/NHLBI harmonised criteria.^22^ Diabetes mellitus was defined as fasting plasma glucose ≥7.0 mmol/L or use of glucose-lowering medication.

Dyslipidaemia was defined as the use of lipid-lowering medication or an LDL cholesterol concentration above 3.0 mmol/L. Current smoking was defined as self-reported current smoking, with former and never smokers combined as non-smokers. Alcohol use was assessed as any self-reported alcohol consumption, and consumption level was categorised using sex-specific weekly limits: low (men 0–4, women 0–2 glasses/week), moderate (men 5-14, women 3-7 glasses/week) and high (men >14, women >7 glasses/week). Sufficient exercise was defined as meeting the Dutch physical activity norm, i.e. at least 30 minutes of moderate-to-vigorous activity on at least 5 days per week, as assessed by the SQUASH questionnaire at baseline only.^23^

We used Food Frequency Questionnaires (FFQs) to record dietary intake, which have been validated in this multi-ethnic population.^24^ From the FFQs, the intake of total calories and the intake of nutrient groups (including carbohydrates, fibre, protein, animal protein, fatty acids, saturated fat, sodium) was calculated based on a nutrient database derived from the Dutch Composition Table 2011.

### Faecal sample collection and participant selection

All HELIUS participants were asked to bring fresh stool samples in a stool tube provided by the research team. These samples were directly transported to the central laboratory and stored at -80°C. All available paired samples (baseline and follow-up sample of the same participant) with sufficient counts (>13,000) of participants that did not use antibiotics in the three months prior to sample collection, in total 1903, were included in the analyses (flowchart in **Supplementary Figure 1**). Shotgun metagenomic sequencing was restricted to Dutch and South-Asian Surinamese participants, the two largest ethnic groups. Participants were eligible if they had not used antibiotics at baseline. Eligible participants were matched 1:1 between the two ethnic groups using the MatchIt R package using nearest-neighbour matching on the continuous variables age and BMI and the factors sex, metformin use, proton pump inhibitor use, statin use, and antihypertensive use. Matched pairs were ranked by the sum of scaled absolute differences in age and BMI, and the 238 best-matched pairs (476 participants) were selected for shotgun sequencing.

### 16S rRNA sequencing

The protocols for faecal sample collection were identical for baseline and follow-up, but sequencing was performed at different time points (in 2016-2017 and 2023-2025, respectively) and locations (Gothenburg, Sweden, and Amsterdam, the Netherlands, respectively). DNA was extracted from 150 mg faecal samples using a repeated bead-beating protocol and subsequently purified using the Maxwell RSC Whole Blood DNA kit. The V4 region of the 16S rRNA gene was amplified with 515F/806R primers, duplicate reactions were combined and purified with the NucleoSpin Gel and PCR Clean-up kit, and pooled amplicons were purified with Ampure XP beads. Libraries were sequenced on an Illumina MiSeq platform with 2×250 bp paired-end reads. The reads were processed with a publicly available Nextflow workflow vsearchpipeline (v1.0.0, run with Nextflow v25.10.3). This workflow uses VSEARCH (v.2.30.6) to infer amplicon sequence variants (ASVs).^25^ Paired-end reads were merged, allowing for a maximum of 30 differences, and then quality filtered with a maximum of 1 expected error. The resulting reads were dereplicated, whereafter ASVs were inferred through denoising with the UNOISE3 algorithm (minimum size 10, alpha 2). The ASVs were sorted and singleton ASVs were discarded, as these are most likely spurious. All merged reads were mapped against the ASVs to create a count table. The *assignTaxonomy* (minboot = 80) and *addSpecies* (allowmultiple =3, tryrevcompl = true) functions from the DADA2 R package (v.1.38.0) were used to assign taxonomy to the ASVs, with the SILVA databases (v.138.1) as reference.^26^ MAFFT (v.7.520) was used for sequence alignment, followed by the construction of a phylogenetic tree using FastTree (v.2.1.11) with a generalized time-reversible model (GTR). Phyloseq (v.1.54.0) was used to integrate the ASV table, taxonomy, tree and reference sequences.^27^ Contamination was identified and removed using the decontam R package.^28^ Both frequency-based and prevalence-based decontamination methods were applied. The count table was rarefied to 13,000 counts per sample, resulting in a table with 3806 samples and 9374 ASVs.

### Shotgun metagenomics

Metagenomic sequencing in 476 subjects with baseline and follow-up samples (952 samples) was performed by Novogene (Munich, Germany) on an Illumina HiSeq platform with 150 bp paired-end reads. After metagenomic processing and quality control, one sample with insufficient read depth was excluded, leaving 951 samples from 476 participants available for analysis, of which 950 paired baseline and follow-up samples of 475 individuals. Reads were processed to compositional MetaPhlAn profiles and HUMAnN pathways with a Nextflow workflow (v0.5.0, run with Nextflow v23.10.1).^29^ Read quality was assessed with FastQC v0.12.1 and reads were quality-filtered and adapter-trimmed using fastp v0.23.4.^30^ Human host reads were removed by alignment to the GRCh38 reference genome using Bowtie2 v2.5.2,^31^ with aligned reads filtered using Samtools v1.18. Reads were subsampled to a uniform depth using SeqKit v2.7.0.^32^ Taxonomic profiling was performed with MetaPhlAn v4.0.5 and ChocoPhlAn database vJun23_202307.^33^ We used the utility script *fix_relab_mpa4.py* of MetaPhlAn v4.1.1 to update the taxonomic labelling to the ChocoPhlAn vJun23_202403 database. Functional pathway abundances were estimated using HUMAnN v3.9.^34^

Strain-level profiling was performed with StrainPhlAn 4 in a Nextflow pipeline (v1.0; Nextflow v23.10.1), using MetaPhlAn v4.0.5 alignment files and merged taxonomic profiles as input. For each species-level genome bin (SGB), StrainPhlAn reconstructed sample-specific consensus marker sequences, built a multiple sequence alignment and phylogeny across samples, and provided pairwise phylogenetic distances.^33^ Species-level genome bins (SGBs) were retained for strain-level analysis when at least one sample met StrainPhlAn’s filtering criteria: a sample was included for an SGB if it contained at least 20 of that SGB’s markers, markers were kept if present in at least 50% of included samples, and samples were removed if they retained fewer than 10 markers after marker filtering. For each retained SGB, a phylogeny of the included samples was built (PhyloPhlAn, accurate mode), and pairwise phylogenetic distances were normalised by the total branch length of the tree (normalised genetic distance; nGD). Species-specific strain-sharing thresholds were determined following Valles-Colomer et al.,^35^ with baseline and follow-up samples from the same participant as related pairs and samples from different participants as unrelated pairs. For SGBs with more than 50 related pairs, the threshold was the lowest of either the nGD maximising Youden’s index, or the 5th percentile of the unrelated-pair distribution. For SGBs with 25 to 50 related pairs, the 3rd percentile of the unrelated-pair distribution was used. SGBs with fewer than 25 related pairs were excluded. Two samples were considered to share a strain when their nGD was below the SGB-specific threshold. Of the 1,224 SGBs profiled, 211 had a sharing threshold (135 using the Youden’s index, 18 the 5^th^ percentile and 58 the 3^rd^ percentile method).

Carbohydrate-active enzyme (CAZyme) profiling of paired-end shotgun metagenomic reads was performed using Cayman (v0.10.1) in a Nextflow pipeline. Cayman maps reads to a gut microbial gene catalogue, normalizes counts to RPKM, and annotates families according to a hierarchical substrate scheme covering dietary fibre (DF), mucin, glycosaminoglycan (GAG), and other glycan classes.^36^ Quality filtered reads were functionally annotated using Cayman against the human gut CAZyme database. We additionally calculated mucin-to-DF and GAG-to-DF ratios.

Metagenomic assembly and metagenome-assembled genome (MAG) reconstruction were performed using the nf-core/mag pipeline (v5.1.0) using Nextflow v25.10.0.^37,38^ Prior to assembly, reads were normalized to a target coverage depth of 40x (minimum 2x) using BBNorm (v39.18) to reduce the impact of uneven sequencing depth. Co-assembly was performed per subject (two samples) using MEGAHIT (v1.2.9).^39^ Contigs were binned using MetaBAT2 (v2.17),^40^ MaxBin2 (v2.2.7),^41^ and CONCOCT (v1.1.0),^42^ and bins were subsequently refined using DAS Tool (v1.1.7).^43^ Bin quality was assessed using CheckM2 (v1.1.0),^44^ and then filtered for >70% completeness and <10% contamination. This resulted in a set of 12,129 bins with a median completeness of 92.78% (interquartile range (IQR) 86.42-97.52) and median contamination of 3.50% (IQR 1.83-5.45). Taxonomic classification of the resulting MAGs was performed using GTDB-Tk (v2.5.2) against the GTDB release226 database.^45^

Functional annotation of 184 *Alistipes putredinis* MAGs was performed using a Nextflow annotation pipeline (v1.1 run with Nextflow 25.10.3, link in Code Availability), which uses Bakta (v1.10.4) for genome annotation,^46^ and Diamond (v2.1.12) to search predicted proteins against the Virulence Factor Database (VFDB) for identification of virulence factors.^47,48^ Hits were retained if they had at least 30% amino acid identity to the VFDB entry and a bitscore of at least 50, retaining 24,166 of 35,037 hits (69%; maximum E-value among retained hits 2×10⁻⁶). Each retained hit was assigned to a virulence factor category by matching its VFG accession to the category given in the VFDB entry headers; hits whose category could not be resolved were excluded. Hits were then counted per MAG and per category. For comparisons between Dutch and South-Asian Surinamese participants, counts were expressed as a proportion of that MAG’s total virulence factor hits.

We additionally constructed a phylogenetic tree of the *A. putredinis* MAGs using Panaroo (v1.6.0) and IQTree3 (v3.1.1) (link in Code Availability, run with Nextflow v25.10.3).^49,50^ For the 164 participants whose own co-assembled A. putredinis MAG was detected at both timepoints, strain retention was assessed directly with inStrain, since MAG-based clade assignment cannot itself detect within-person strain turnover (each participant contributes a single co-assembled MAG). Comparisons focused on the 126 participants with ≥5× depth at both timepoints. For each, reads from each timepoint were mapped to that participant’s own MAG with Bowtie2 (v2.5.5), sorted and indexed with samtools (v1.24), and profiled separately with inStrain profile (v1.10.0; min_read_ani 0.95). The two timepoint profiles were then compared with inStrain compare (min_cov 5, breadth 0.5) to obtain the population average nucleotide identity (popANI) between baseline and follow-up. Comparisons with <50% of the genome compared between timepoints were excluded, leaving 120 valid comparisons. A pair of timepoints was classified as the same strain at popANI ≥99.999%. To determine whether within-person retention was distinguishable from the background similarity of unrelated members of the same phylogenetic clade, a between-person comparison was performed: for each clade, one high-quality anchor MAG was compared by inStrain against baseline reads from multiple other, unrelated participants in the same clade. No between-person comparison reached the same-strain threshold.

Antibiotic resistance genes were detected using a Nextflow workflow that was based on the funcscan nf-core pipeline, run with Nextflow v25.10.3.^38,51^ Sequence quality was assessed with SeqKit (v2.9.0),^32^ and contigs were taxonomically classified using MMseqs2 (v2.17).^52^ Open reading frames in the MAGs were predicted with Pyrodigal (v3.6.3),^53^ and the resulting protein sequences were screened for ARGs using AMRFinderPlus (4.0.23).^54^ ARG annotations were normalized to the Antibiotic Resistance Ontology (ARO) using argNorm (v0.5.0).^55^ For ARG abundance quantification, detected ARG nucleotide sequences were extracted and deduplicated at 95% nucleotide identity per sample group using CD-HIT-EST (v4.8.1).^56^ Metagenomic reads were mapped to the resulting participant-specific ARG catalogue using BWA-MEM2 (v2.2.1),^57^ and abundance metrics were calculated, so baseline and follow-up samples of one participant were quantified against the same reference. The number of predicted genes across each participant’s MAGs was recorded as a proxy for assembly yield.

All Nextflow workflows were run on a high-performance cluster (Snellius, the Netherlands) using Singularity software containers. All repositories with workflows are referenced in the Code Availability section.

### Statistics

For tables with participant characteristics, continuous variables were reported as mean ± standard deviation, except triglycerides, which is a variable with non-normal distribution, which was reported as median [interquartile range]. Categorical variables were reported as counts and percentages. No statistical comparisons between timepoints were performed in the main tables. For cross-sectional comparisons between ethnic groups within each timepoint, continuous variables were compared with t-tests or Mann-Whitney U tests (for non-normal variables), and categorical variables were compared using chi-squared tests. All tables were generated using the tableone package (v0.13.2).

To visualize baseline dietary patterns across ethnic groups, principal component analysis (PCA) was applied to FFQ-derived nutrient variables (carbohydrates, fibre, total protein, animal protein, fatty acids, saturated fat, sodium). Energy adjustment was performed using a residual method (Willett’s method),^58^ in which nutrient intake was regressed on total energy intake at baseline using a linear regression model, followed by zero-mean-unit variance scaling. The first two principal components were visualised and used in downstream analyses described below. Nutrient variables were visualized using an energy-adjustment dividing by 1000 kcal and compared across ethnic groups using pairwise Mann Whitney U tests.

In the 16S sequencing data analyses, alpha diversity was assessed by calculating the Shannon index for each sample and beta diversity was quantified using Bray-Curtis distances. Longitudinal community-level change was tested by permutation of multivariate analysis of variance (PERMANOVA) using the adonis2 function from the vegan R package with 999 permutations. Because participants contributed samples at two timepoints, permutations were restricted to within-individual blocks using the how function from the permute package, exchanging only timepoint labels within each individual and preserving the paired structure of the data. Bray-Curtis distances (baseline to follow-up) were regressed on age, sex, BMI, metformin, proton pump inhibitors, and follow-up time in a linear model; confounder-adjusted dissimilarities were obtained as model residuals shifted to the grand mean. A sensitivity analysis additionally adjusted for dietary principal components (PC1 and PC2 derived from dietary analyses described above). Next, we tested homogeneity of dispersion across the six ethnic groups at each timepoint using PERMDISP (betadisper with type “centroid”), with permutest using the same blocked permutation as for the PERMANOVA test. Within each ethnic group, change in distance to centroid between baseline and follow-up was tested with a paired Wilcoxon signed-rank test. Whether this change differed by ethnicity was tested with a linear mixed model of distance to centroid with an ethnicity × timepoint interaction and a random intercept for participant. Pearson correlations were used to assess associations between follow-up time and Bray-Curtis dissimilarity, and between baseline Shannon index and Bray-Curtis dissimilarity. Ethnic group comparisons for Bray-Curtis dissimilarity and alpha diversity were performed with Mann-Whitney U tests.

Associations between a range of covariates (cardiometabolic risk factors, disease, medication, gut microbiota diversity, diet) at baseline and microbiota instability over time using linear regression (outcome: Bray-Curtis dissimilarity). Continuous predictors (e.g. age, BMI) were z-scored to allow comparison of effect sizes. Analyses were run both across the full cohort and stratified by ethnic group. We additionally performed a sensitivity analysis in which we excluded all participants that had the exposure of interest at the follow-up timepoint (‘starters’) per variable, since these could have influenced the associations. Associations between baseline predictors and longitudinal alpha diversity change were assessed by linear mixed models (LMM) with the formula *Shannon ∼ predictor × timepoint + follow-up duration + (1 | participant ID)*. Per covariate of interest, only complete cases were included in the models. P-values were adjusted for multiple testing using the Benjamini-Hochberg false discovery rate (BH-FDR).

The analyses of the shotgun metagenomics data were performed in a matched subgroup of Dutch and South-Asian Surinamese participants. Since these participants were matched for age, sex and BMI, we did not additionally adjust the downstream analyses for these covariates. Species-level analyses were performed using MetaPhlAn abundance profiles. Species were included if they were detected in more than 20% of samples in either the Dutch or South-Asian Surinamese group >0.1% relative abundance, resulting in a list of 181 species. Abundances were log10-transformed with a pseudocount of 0.01 prior to modelling. Longitudinal ethnic differences were tested using linear mixed models (LMMs) with the formula: *species ∼ ethnicity × timepoint + follow-up time + (1 | participant ID)*. Cross-sectional ethnic differences at baseline and follow-up were additionally tested in separate linear models, without adjustment at baseline and adjusted for follow-up duration at follow-up. These analyses were not adjusted for age, sex, or BMI since the two ethnic groups were matched on these variables. P-values were adjusted for multiple testing (BH-FDR).

*Alistipes putredinis* was selected for strain-level analysis as the species with the lowest FDR-adjusted p-value for the ethnicity × timepoint interaction in the species-level linear mixed models. The maximum-likelihood tree of the 184 *A. putredinis* MAGs was visualised using ggtree (v4.0.4) and treeio (v1.34.0). Clades were defined as the largest monophyletic subtrees whose maximum pairwise patristic distance did not exceed a threshold h = 0.03, applied top-down from the root. h = 0.03 was chosen as the centre of the widest stable plateau in cluster count (9 clusters, unchanged for *h* = 0.029– 0.040). This yielded four clades with ≥5 MAGs (Clades I–IV), plus five additional MAGs in singleton or near-singleton groups. Clade assignment was unchanged when clustering was repeated on the tree pruned to the 51 high-quality MAGs (≥90% completeness, <5% contamination): all four clades were recovered in this subset. Clade-level comparisons of virulence factor categories included all MAGs in clades with ≥5 tips (n = 177 across four clades) and were made using Kruskal-Wallis tests and multiple testing correction (BH-FDR).

For each within-person inStrain comparison, we computed population-level SNP density (SNPs per Mb compared), alongside the equivalent distribution from between-person comparisons between unrelated individuals carrying the same *A. putredinis* clade. Comparisons meeting the popANI same-strain threshold were classified as stable (popANI ≥99.999%). Comparisons below this threshold were further classified as divergent but related when their SNP density fell below the between-person floor, or consistent with replacement when it fell within the between-person range. popANI itself was compared between ethnic groups with a Mann Whitney U test. Within-strain microdiversity (nucleotide diversity, π) was compared between baseline and follow-up with paired Wilcoxon signed-rank tests, restricted to participants whose strain was retained over time. Differential change in microdiversity between ethnic groups over time was assessed using a linear mixed model (ethnicity*time interaction). Cross-sectional differences in nucleotide diversity between ethnic groups were tested at each timepoint separately with two-sided Mann-Whitney U tests. Because sequencing depth could have inflated nucleotide diversity estimates, the ethnic difference was additionally tested with linear regression of log-transformed nucleotide diversity on ethnicity and sequencing coverage, run separately at each timepoint.

Strain-level stability was quantified as the percentage of detected strains shared between a participant’s baseline and follow-up sample using the StrainPhlAn analyses results (see ‘Shotgun metagenomics’ section). Associations of strain sharing percentage with a panel of covariates (cardiometabolic risk factors, disease, medication, gut microbiota diversity, diet) were assessed by linear regression, adjusting for follow-up time. Analyses were run both across the full cohort and stratified by ethnic group (Dutch and South-Asian Surinamese). Strain stability was correlated to age, follow-up time, Shannon diversity and Bray-Curtis dissimilarity in the shotgun metagenomics compositional profiles using Spearman’s correlations. Strain sharing percentage per participant was compared between ethnic groups and tested with a Mann-Whitney U test. Differential strain stability between ethnic groups was tested with a logistic regression model, only including the 153 SGBs that were present in more than 50 sample pairs.

Carbohydrate-active enzyme (CAZyme) family abundances (RPKM) were log10-transformed and filtered to families present in more than 5% of samples, resulting in 466 families. HUMAnN pathways were filtered for relative abundances (CPM) higher than 0.0025 in more than 25% of samples, which resulted in 140 pathways. The differential change of CAZyme families and pathway abundances over time was modelled with linear mixed models (formula: *log10(abundance + half the minimal non-zero value) ∼ ethnicity × timepoint + follow-up duration + (1 | participant ID)*). For CAZymes, an additional model was used to assess the impact of baseline diet in 232 participants with available dietary data, adjusting for the dietary principal components. Cross-sectional ethnic differences for HUMAnN pathways and CAZyme families were tested at each timepoint with linear regression of log10-transformed abundances on ethnicity, and p-values were corrected for multiple testing (BH-FDR). Canberra distances (more sensitive to rare families) of CAZyme compositional profiles were calculated using the vegan package and differences were tested with PERMANOVA (999 permutations). Mucin/DF and GAG/DF ratios and richness (the number of CAZyme families per sample) were log10-transformed and compared between ethnic groups at each timepoint using Mann-Whitney U tests, and differential change over time was tested with a linear mixed model. The richness model was adjusted for the number of reads that mapped to CAZyme genes. To identify which substrate class drove ratio differences, the substrate classes were analysed separately as log10-transformed relative abundance of total CAZymes, using the same tests.

Total ARG burden was the sum of ARG-mapped reads per million total reads (RPM), log10-transformed. ARG Shannon diversity was computed from RPKM proportions of detected genes, so that longer genes would not contribute disproportionately. For gene-level analyses, ARGs detected in fewer than 5% of samples were excluded. Gene-level abundance was expressed as counts per million reads and log10(CPM + 1) transformed. Cross-sectional differences in gene and class prevalence between Dutch and South-Asian Surinamese participants were tested at each timepoint with chi-square tests, and gene-level abundance differences with linear models with ethnicity as predictor. Longitudinal analyses used linear mixed models with a random intercept per participant and follow-up time as a covariate. Total burden, richness, Shannon diversity and each gene were modelled for the ethnicity*timepoint interaction. BH-FDR correction was applied to gene-level p-values. Because ARG presence depends on sequencing depth and on how many genes were assembled and binned, we performed sensitivity analyses, in which we repeated the models with sequencing depth and the number of predicted genes per participant’s MAGs as covariates. We also repeated Shannon analyses requiring at least 5 or 10 mapped reads for detection, and repeated the timepoint contrast in participant pairs with depth differing by less than 0.1 log10 units.

All statistical analyses were conducted in R 4.5.3 in Positron 2026.04.0. Linear mixed models were fitted using lme4 (v1.1-38) and lmerTest (v3.2-1); 95% confidence intervals were estimated by the Wald method. P-values after multiple testing correction are reported throughout as q-values. Figures were generated using ggplot2 (v4.0.2) and ggpubr (v0.6.3). R package versions were controlled using pixi (v0.62.2), and the pixi environment can be reconstructed using the pixi.toml and pixi.lock files that can be found in the code repository.

## Results

### Baseline microbiome diversity is associated with longitudinal stability

We included 1,903 participants across six ethnic groups from the HELIUS cohort with both baseline and follow-up faecal samples (flowchart presented in **Supplementary Figure 1**). The population had a baseline age of 52.5 ± 10.1 years, of whom 52.3% were women (**Supplementary Table 1-2**). Follow-up time was on average 6.3 ± 1.3 years and differed modestly between ethnic groups: the Moroccan and Turkish groups had the shortest average follow-up time (5.6 ± 0.8 and 5.7 ± 1.0 years, respectively), while the Ghanaian group had the longest follow-up (6.7 ± 0.9 years; **Figure 1A**). We measured gut microbiota composition using 16S sequencing, and found a significant shift in microbiome composition between the two timepoints (**Figure 1B**; PERMANOVA R^2^ = 0.023, p = 0.001). To characterize within-individual microbiome change over time, we computed Bray-Curtis dissimilarity between baseline and follow-up samples of the same individual in all ethnic groups using 16S rRNA sequencing data. Dissimilarity increased with longer follow-up duration (r = 0.11, p = 8.1×10^−6^; **Figure 1C**). The degree of change differed substantially between ethnic groups in ways that were not accounted for by differences in follow-up duration or covariates such as age, sex, body mass index (BMI), metformin and proton pump inhibitor (PPI) use (**Figure 1D**; unadjusted distance in **Supplementary Figure 2A**). The Moroccan group showed the greatest compositional change over time, followed by the Ghanaian and African Surinamese groups, while the South-Asian Surinamese and Dutch groups were most stable compared to their baseline samples. Differences between the Moroccan group and four other ethnic groups were significant (African-Surinamese, Turkish, Dutch and South-Asian Surinamese). Adjusting these differences additionally for dietary intake in a subgroup of participants (N = 799, 42%) led to an attenuation of the ethnic differences and loss of significance between groups (**Supplementary Figure 2B**).

**Figure 1:**
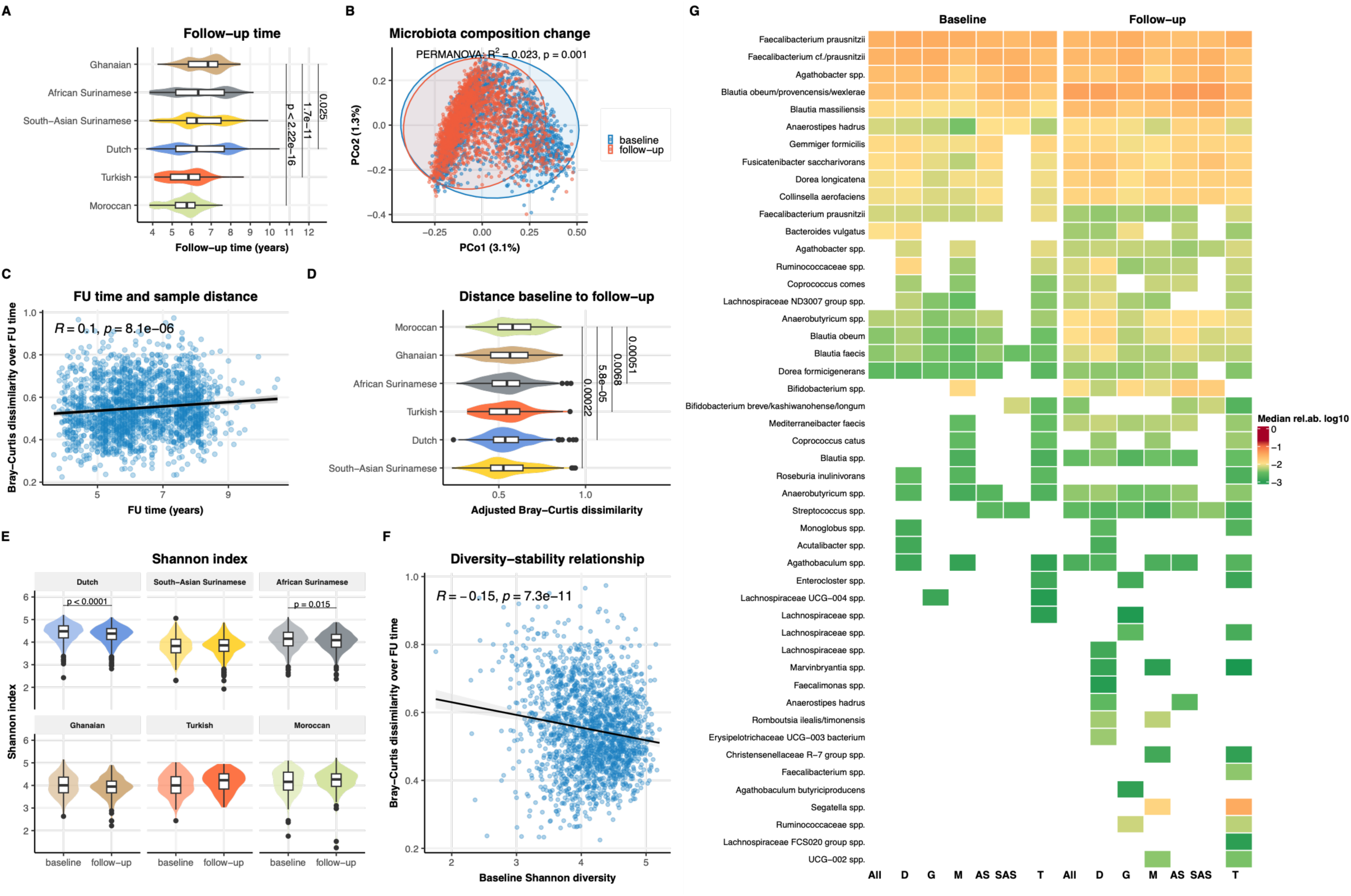
Longitudinal gut microbiota stability across ethnicities in the HELIUS cohort. (A) Distribution of follow-up duration (years) by ethnic group, shown as violin plots with box plots. Differences tested with a Mann Whitney U test. (B) Principal coordinate analysis (PCoA) of Bray-Curtis distances between all samples, with the colours indicating time points (baseline or follow-up). Differences between timepoints tested with PERMANOVA. (C) Scatter plot of Bray-Curtis dissimilarity between paired baseline and follow-up samples and follow-up duration (years). Pearson’s correlation is shown; each point represents one participant. (D) Bray-Curtis dissimilarity between paired samples stratified by ethnic group, adjusted for baseline age, sex, BMI, metformin, PPI, and follow-up time using linear models. (E) Shannon diversity at baseline and follow-up per ethnic group. (F) Scatter plot of baseline Shannon diversity against Bray-Curtis dissimilarity over follow-up. Pearson’s correlation is shown. (G) Heatmap showing median relative abundance (log10-transformed) of core ASVs, defined as ASVs with ≥95% prevalence in at least one ethnic group at baseline or follow-up, across six ethnic groups (Dutch [D], Ghanaian [G], Moroccan [M], African Surinamese [AS], South-Asian Surinamese [SAS], Turkish [T]) and the overall cohort (All), at baseline (left) and follow-up (right). Only fields where prevalence ≥95% are shown; fields below this threshold are left blank. Rows are clustered hierarchically. Colour indicates median relative abundance (log10-transformed), ranging from green (low) to dark red (high). FU = follow-up.

The R² for ethnicity in gut microbiota composition at baseline was 0.0438, comparable to full baseline sample in Deschasaux *et al.*,^4^ while it was 0.0368 at follow-up. All 15 ethnicity pairs remained significantly separated at both timepoints (R² 0.008– 0.049). The difference was largest between Dutch and South-Asian Surinamese participants and smallest between Turkish and Moroccan participants. In a combined model with permutations restricted within participants, the ethnicity effect differed between timepoints (ethnicity × timepoint R² = 0.003, p = 0.001), although this interaction was much smaller than the main effects of ethnicity and timepoint. The pairwise between-group distances in 15 ethnicity-pairs on average showed a small decrease over time (median ΔR² −0.0029 [IQR −0.0042 – −0.0015] over six years; **Supplementary Figure 2C-D**). However, within-group dispersion also differed across ethnicity-timepoint groups (PERMDISP p = 0.001), and the change in dispersion over time differed by ethnicity (linear mixed model, ethnicity × timepoint p = 0.0012). The distance to the group centroid decreased in Dutch, South-Asian Surinamese, African Surinamese and Turkish participants, but not in Ghanaian or Moroccan participants (**Supplementary Figure 2E**). The differential change in Bray-Curtis distance over time between ethnicities therefore reflects both shifts in group centroids and differential changes in within-group dispersion. We also associated each non-Dutch participant’s distance to the Dutch centroid to residence duration. Distance to the Dutch centroid decreased with longer residence (-0.007 per decade, 95% CI −0.0111 – −0.0029, p < 0.001; **Supplementary Figure 2F**), although the effect was very small compared to the median distance to the Dutch centroid (0.696).

At baseline, clear ethnic differences in alpha diversity were present. Shannon diversity was highest in the Dutch group and lowest in the South-Asian Surinamese group (**Figure 1E; Supplementary Figure 3**). Longitudinally, the Dutch and African Surinamese groups showed significant decreases in Shannon diversity, while other ethnic groups showed a slight increase, resulting in a partial convergence of alpha diversity between groups at follow-up. A higher baseline Shannon diversity was associated with a more stable gut microbiota composition over time, as reflected by lower Bray-Curtis dissimilarity compared to baseline (**Figure 1F**; Pearson’s r = −0.15, p = 7.3×10^−11^).

High-prevalence amplicon sequence variants (ASVs; ≥95% prevalence in at least one ethnic group) showed some differences in breadth across ethnic groups (**Figure 1G**). At baseline, the Dutch, Moroccan, and Turkish groups shared a relatively broad set including *Faecalibacterium prausnitzii*, *Agathobacter* spp., *Anaerostipes hadrus*, several *Blautia* spp., and a number of *Lachnospiraceae* members that were less consistently represented in the South-Asian Surinamese group. At follow-up, this difference was less apparent, with the South-Asian Surinamese group gaining high-prevalence status for several of these taxa, consistent with the partial alpha diversity convergence observed over time.

### Cardiometabolic disease is associated with greater compositional change

Over the follow-up time, there was a slight increase in BMI (**Supplementary Table 1**; +0.38 kg/m²) and a decrease in the proportion of participants who actively smoked or reported moderate to high alcohol consumption. At baseline (22.2%) and follow-up (31.9%), diabetes was considerably more prevalent in the South-Asian Surinamese participants than in any other ethnic group (4.2-14.2% baseline and 5.7-19.5% follow-up; **Figure 2A**). Dyslipidaemia and metabolic syndrome were most prevalent in South-Asian Surinamese participants at both timepoints (81.8% and 62.5% at follow-up), while hypertension was most prevalent in the Ghanaian group (64.5% at follow-up).

**Figure 2.**
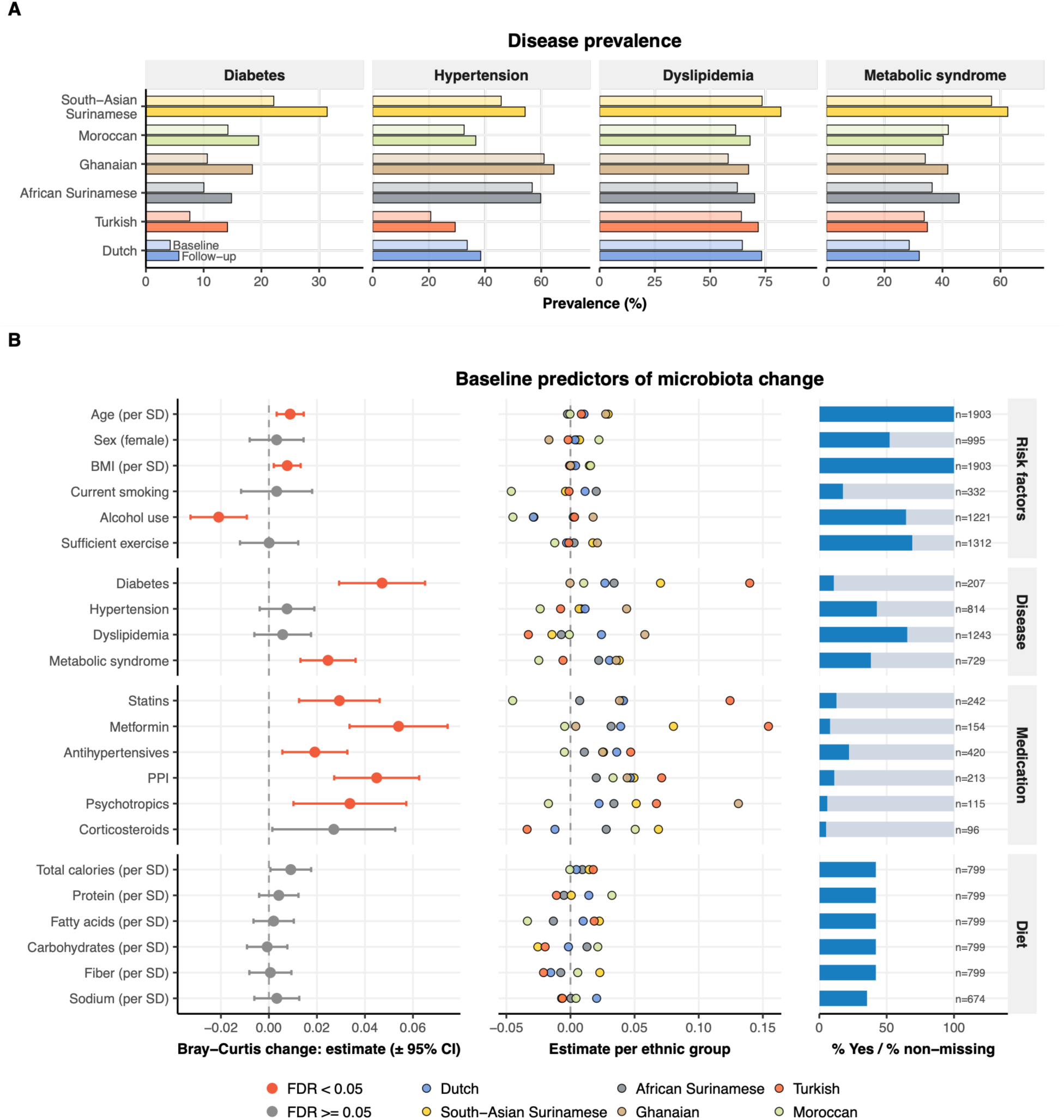
Cardiometabolic disease burden and gut microbiota instability in the HELIUS cohort. (A) Prevalence of diabetes, hypertension, dyslipidaemia and metabolic syndrome at baseline (light bars) and follow-up (dark bars) across six ethnic groups. (B) Left panel: Effect of baseline predictors on microbiota instability, expressed as linear regression estimates (± 95% CI) of Bray-Curtis dissimilarity over the follow-up period. Continuous variables were plotted per standard deviation (SD), all other variables are categorical (Yes/No) with No as reference. Middle panel: Estimates of the same model shown per ethnicity. Right panel: the number of cases that were labelled as Yes (categorical variables), with grey stacked bar showing the No group, or number of cases with available data (continuous variables). FDR: Benjamini-Hochberg false discovery rate (multiple testing correction). PPI: proton pump inhibitor. BMI: body mass index.

To test whether ethnic disparities in lifestyle, risk factors and disease burden at baseline were associated with longitudinal compositional change, we tested the effects of these variables on Bray-Curtis dissimilarity between paired baseline and follow-up samples. Risk factors such as higher age (β = 0.009 (95%CI 0.003 – 0.014), q = 0.006) and BMI (β = 0.008 (0.002 – 0.013), q = 0.017) at baseline were associated with more microbiota compositional change (**Figure 2B**). Alcohol use showed a reversed pattern: alcohol use at baseline was associated with a more stable composition (β = -0.021 (-0.033 – -0.009), q = 0.002). Participants with cardiometabolic diagnoses at baseline showed greater compositional change over time than those without, with the most pronounced effect observed for diabetes (β = 0.047 (0.029–0.065), q = 2.90 ×10^−6^). This association was present across ethnic groups but was not uniform: the relationship between diabetes and microbiome instability was strongest in the South-Asian Surinamese and Turkish groups. Medication use at baseline such as metformin (β = 0.054 (0.034–0.074), q = 2.39 ×10^−7^), proton pump inhibitors (PPI; β = 0.045 (0.027– 0.063), q = 7.15 ×10^−7^), psychotropic medication (β = 0.034 (0.010–0.057), q = 0.005), statins (β = 0.029 (0.013 –0.046), q = 0.0006) and antihypertensive drugs (β = 0.019 (0.006–0.033), q = 0.006) were also associated with more change. Of the separate dietary nutrient groups derived from food frequency questionnaires (FFQ), none of the groups were associated with compositional change, while we did observe ethnic differences in dietary patterns at baseline (**Supplementary Figure 4-5**). For all variables that were available at the follow-up timepoint, we performed a sensitivity analysis in which we excluded all participants that had the exposure of interest at the follow-up but not at the baseline timepoint (the ‘starters’ per variable). All associations persisted in this sensitivity analysis (**Supplementary Figure 6**).

We performed a similar analysis with the Shannon index as measure of alpha diversity using linear mixed models (time × covariate interactions), and found that BMI (β_BMI×time_ = 0.041 (0.019–0.062), q = 0.003) and energy-adjusted carbohydrate intake (β_carbohydrates×time_ = 0.046 (0.014–0.079), q = 0.029) had positive associations with Shannon index change over time, while female sex (β_sex×time_ = -0.060 (-0.104 – -0.017), q = 0.029), alcohol use (β_alcohol×time_ = -0.093 (-0.139 – -0.048), q = 0.001) and energy-adjusted fatty acid intake (β_fattyacids×time_ = -0.046 (-0.079– -0.013), q = 0.029) showed negative associations (**Supplementary Figure 7**).

### Ethnic differences in gut microbiota composition persist at species and strain level

To investigate gut microbiome dynamics at species level, we performed shotgun metagenomic sequencing in a matched subset of Dutch (n = 238) and South-Asian Surinamese (n = 238) participants, the two groups with the most divergent cardiometabolic disease prevalence (**Supplementary Tables 3-4**). Approximately half of the shotgun metagenomics subgroup had dietary data available at baseline (N = 232). The South-Asian Surinamese participants had significantly higher energy-adjusted protein, animal protein and carbohydrate intake, while the Dutch participants had higher fatty acid and fibre intake (**Supplementary Figure 8**). The follow-up time was similar between these two subgroups (p = 0.32).

We first replicated the alpha and beta diversity findings of the 16S data in the shotgun metagenomics data (**Supplementary Figure 9**). The ethnic differences in beta diversity were attenuated from baseline to follow-up (R^2^ = 0.047 to R^2^ = 0.037 at follow-up). Linear models identified 78 species that were differentially abundant between ethnicities at baseline, with a number of *Bifidobacterium* species higher abundant at South-Asian Surinamese and several *Alistipes* species higher abundant in Dutch participants (**Supplementary Figure 10**). At follow-up, there were 66 species with differential abundance between groups in a model that was additionally adjusted for follow-up time, of which 61 overlapped with the 78 differentially abundant baseline species (**Supplementary Figure 10**). Next, we used linear mixed models to test differential changes of the groups over time, which identified 9 species that changed differentially between Dutch and South-Asian Surinamese participants (ethnicity x timepoint interaction; **Figure 3A**). To characterise the nature of these longitudinal differences, we classified interaction patterns as convergence (cross-sectional differences present at baseline attenuating over time), divergence (existing differences amplifying over time), or emergence (differences absent at baseline becoming significant at follow-up). The predominant pattern was convergence because essentially all differentially changing species showed loss of significance or lower significance of the ethnic differences at follow-up. *Alistipes putredinis* showed the most significant change over time, with a positive interaction estimate for ethnicity x time, leading to convergence of the South-Asian Surinamese group to the higher baseline abundance of Dutch participants (**Supplementary Figure 11**).

**Figure 3.**
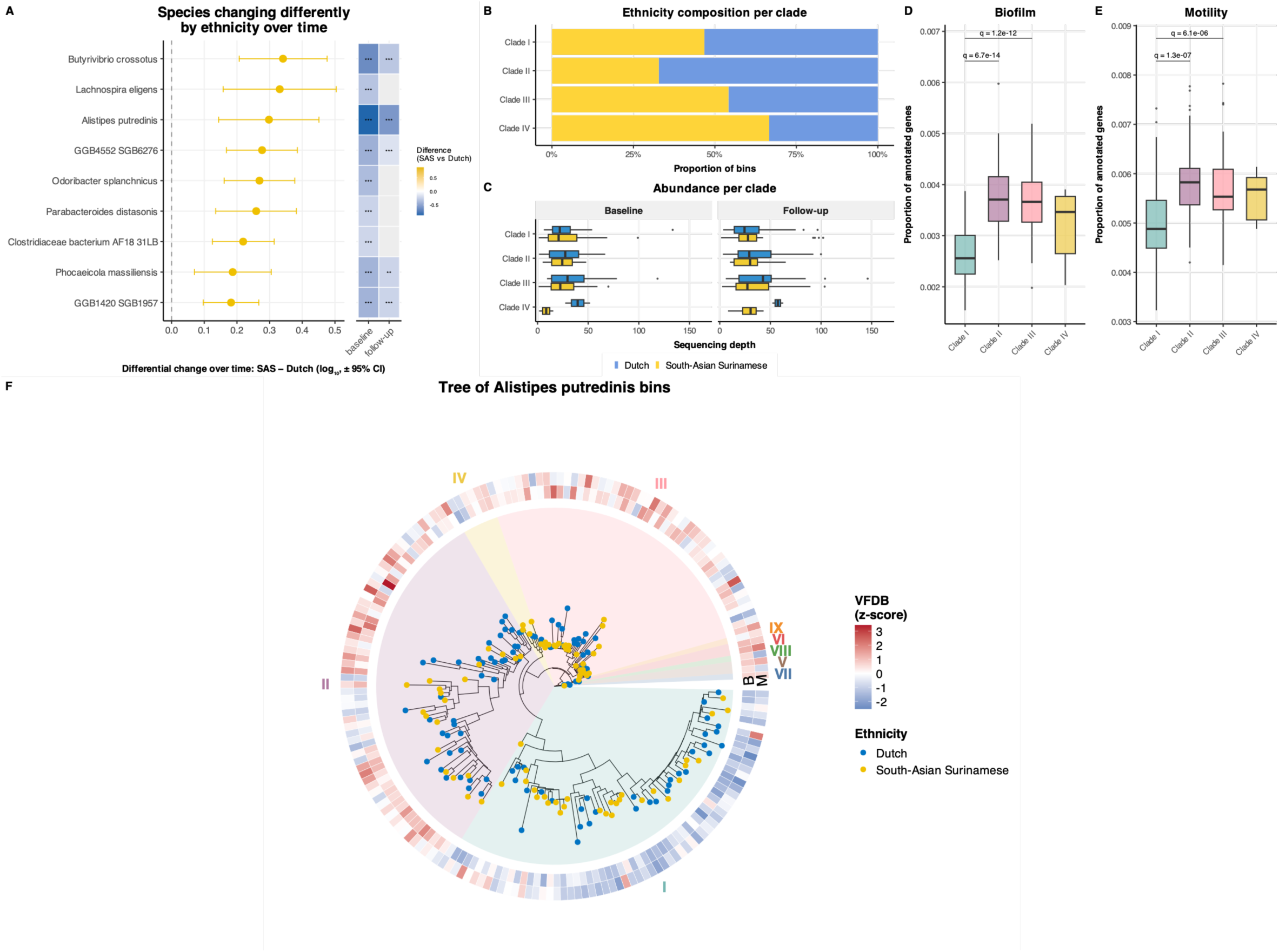
Species-level dynamics between Dutch and South-Asian Surinamese participants (shotgun metagenomics). (A) Forest plot of species showing a significant ethnicity × timepoint interaction in linear mixed models (LMM). Points indicate interaction effect sizes for South-Asian Surinamese compared to Dutch participants (β) with 95% confidence intervals. (B) Stacked bar chart showing the proportion of Dutch and SAS participants per phylogenetic clade. (C) Boxplots showing the baseline and follow-up MAG coverage per ethnic group per clade. Differences between ethnic groups were tested per clade using Mann-Whitney U tests; none of the tests were significant. (D-E) Boxplots of VFDB gene category proportions across clades. Pairwise comparisons with Mann-Whitney U test, corrected for multiple testing using Benjamini-Hochberg; only comparisons significant after Kruskal-Wallis (biofilm: p = 1.4×10^−16^ and motility: p = 2.5×10^−8^) are shown. (F) Maximum-likelihood phylogenetic tree of *Alistipes putredinis* metagenome-assembled genomes (MAGs). Coloured sectors indicate phylogenetic clades (I–IX). Tip points indicate host ethnicity (colour) of each MAG. The outer heatmap ring shows z-scored virulence factor database (VFDB) gene proportions per MAG, with the inner ring showing biofilm (B) category and the outer ring showing motility (M) category gene content proportions.

The baseline colonization prevalence of *A. putredinis* differed strongly by ethnicity (90.7% of Dutch vs. 56.5% of South-Asian Surinamese participants; Fisher’s exact p = 8.3×10⁻¹⁸). Focusing to participants without *A. putredinis* at baseline, the proportion who acquired detectable *A. putredinis* by follow-up did not differ significantly between groups (31.8% of Dutch, n=22, vs. 39.8% of South-Asian Surinamese, n=103; OR=1.41, p=0.63). Thus, although South-Asian Surinamese participants showed more acquisition events, the acquisition rate between ethnicities was not different.

To determine whether the convergence of *Alistipes putredinis* extended to strain composition and MAG gene annotation, we performed phylogenetic clustering on 184 *Alistipes putredinis* MAGs (completeness ≥70%, contamination <10%), which identified four major clades and five small clusters of 1 or 2 MAGs. Four clades had five or more tips (Clade I, n=62; Clade II, n=61; Clade III, n=48; Clade IV, n=6; **Figure 3B-C, F**), without any differences in completeness or contamination between clades. Clade assignment was robust to MAG quality: restricting to 51 high-quality MAGs (completeness ≥90%, contamination <5%) reproduced the same clade structure. Clade II had significantly more Dutch than South-Asian Surinamese participants (Fisher’s exact OR 0.47 for South-Asian Surinamese vs Dutch, p = 0.026), while the other clades showed no ethnic differences in clade membership. Of 14 virulence-factor (VF) categories screened with the virulence factor database (VFDB), biofilm-and motility-associated gene content differed most strongly across clades (Kruskal-Wallis, both p < 1×10^−7^; **Figure 3D-E**). Clades II and III had significantly higher proportions of Biofilm and Motility-associated gene content than Clade I (pairwise Wilcoxon, q < 1×10^−5^ for the four comparisons).

Using inStrain population-level ANI (popANI) comparisons between independently mapped baseline and follow-up reads, only 20.0% of 120 within-person comparisons met the threshold for an identical *A. putredinis* strain (popANI≥0.99999) over the six-year follow-up. Of the remainder, the majority (70.0%) showed only modest divergence, less than the lowest between-person divergence; a smaller fraction (10.0%) showed substantially greater divergence, more consistent with strain replacement (**Supplementary Figure 12**). There were no differences in popANI between ethnicities (p = 0.75). Among the subset of participants whose strain was retained (13 Dutch and 11 South-Asian Surinamese), within-host nucleotide diversity increased significantly from baseline to follow-up, in both groups (paired Wilcoxon, overall p = 1.2×10⁻⁷; Dutch p = 2.4×10⁻⁴; South-Asian Surinamese p = 9.8×10⁻⁴). The change over time was not different between these two ethnic groups (linear mixed model: group*time p = 0.779). However, cross-sectionally, Dutch participants carried more genetically heterogeneous *A. putredinis* populations than South-Asian Surinamese participants (baseline p = 0.005; follow-up p = 0.021; **Supplementary Figure 12**). Microdiversity in *A. putredinis* did not correlate with any of the clinical variables (age, sex, BMI, alcohol, medication use) at baseline. The ethnic difference persisted after adjusting for coverage in a linear regression model (baseline p = 0.011, follow-up p = 0.039).

For a broader strain-level analysis, we used StrainPhlAn for a strain sharing comparison between baseline and follow-up samples per participant. We included 211 SGBs and determined the threshold for strain sharing between the two timepoints as described by Valles-Colomer *et al*.^35^ The proportion of stable strains per participant decreased gradually with longer follow-up time (ρ −0.166, p = 0.0003) but was not associated with baseline age (ρ 0.014, p = 0.75). Other factors that showed associations with strain stability were alcohol use, which was associated with more stability (β = 8.12 (3.71–12.52), q = 0.002), and PPI use which was associated with less strain stability (β = -9.23 (-15.99– -2.48), q = 0.029); **Supplementary Figure 13**). Strain stability was additionally correlated with baseline alpha diversity (ρ = 0.11, p = 0.016) and Bray-Curtis dissimilarity (ρ = −0.47, p = 1.6 ×10^−27^; **Supplementary Figure 14**). The overall strain stability was comparable between ethnic groups (**Supplementary Figure 14**; p = 0.50). Only one SGB showed differential strain stability between ethnicities, *Oscillibacter sp. ER4* showed significantly more stability in South-Asian Surinamese than Dutch participants (OR = 3.57 South-Asian Surinamese vs. Dutch, 95%CI 2.06-6.35, q = 0.0005; **Supplementary Figure 15**).

### Microbial functional capacity shifts differently by ethnicity

To characterise whether longitudinal changes in microbiome composition translated to functional shifts, we profiled metabolic pathway abundances using HUMAnN3. Of 140 pathways present in sufficient prevalence for analysis, 104 differed significantly between ethnic groups at baseline and 97 at follow-up (q < 0.05), with 85 significant at both timepoints. Fourteen pathways showed a significant ethnicity × timepoint interaction (q < 0.05; **Figure 4A**). The predominant pattern was emergence of new differences, observed in eight pathways, followed by convergence in five pathways, with one pathway showing divergence. Four pathways showed new differences over the follow-up period because of a lower decrease in South-Asian Surinamese participants, including gondoate and cis-vaccenate biosynthesis (both long-chain unsaturated fatty acid pathways), L-arginine biosynthesis and 4-amino-2-methyl-5-diphosphomethylpyrimidine biosynthesis. Gluconeogenesis III showed the strongest South-Asian Surinamese-specific increase (β = 0.110, q = 0.005) and was significantly higher in Dutch participants at both timepoints, consistent with convergence (**Supplementary Figure 16-17**). Another converging pathway was L-lysine biosynthesis II, which showed the strongest Dutch-specific increase and was higher in South-Asian Surinamese at both timepoints.

**Figure 4.**
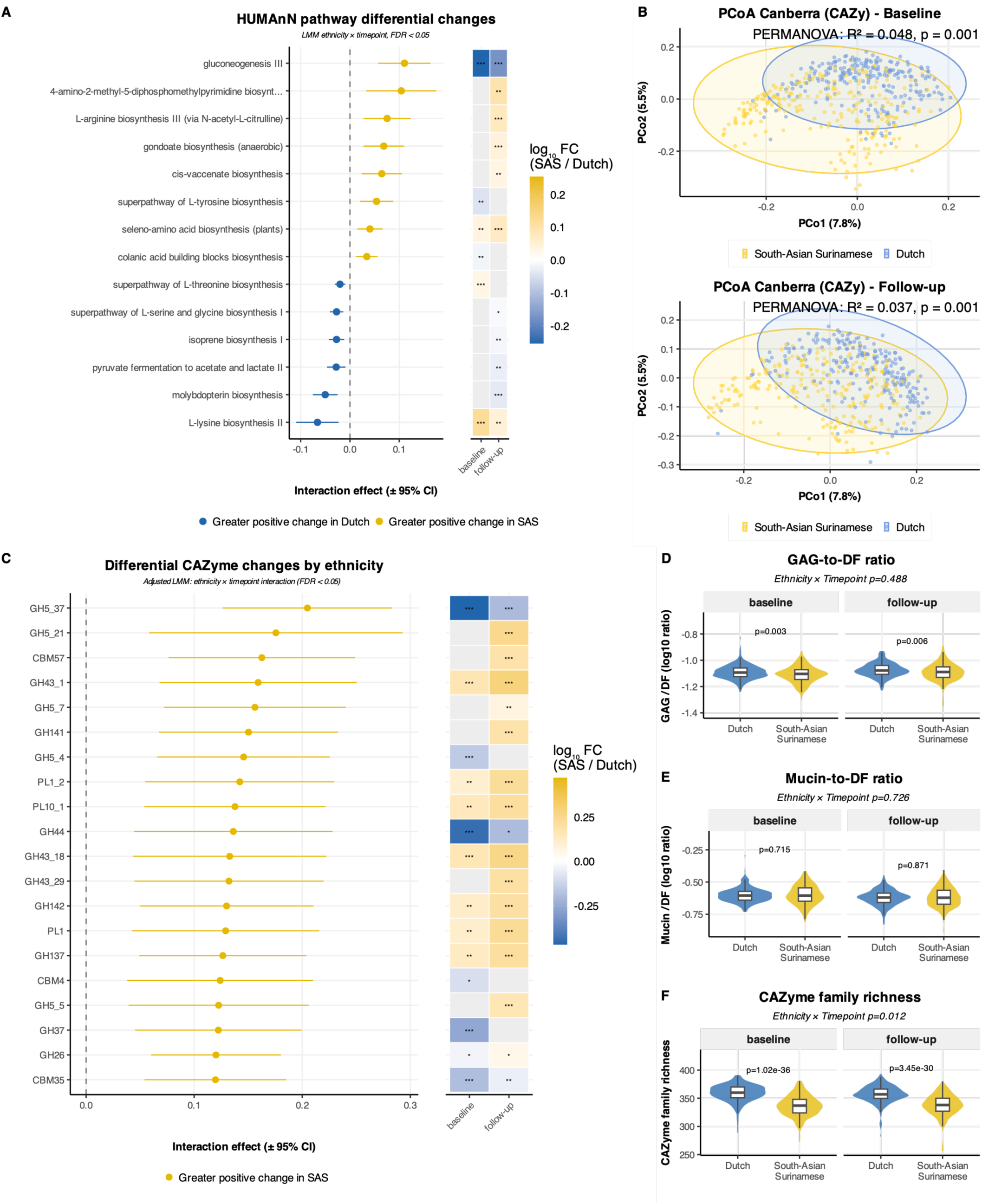
Longitudinal shifts in gut microbiota functional capacity by ethnicity (shotgun metagenomics). (A) Forest plot of HUMAnN pathways with a significant ethnicity × timepoint interaction (LMM, FDR < 0.05). The adjacent heatmap indicates cross-sectional significance at baseline and follow-up. (B) Principal coordinates analysis (PCoA) of CAZyme gene family composition based on Canberra dissimilarity at baseline (left) and follow-up (right), coloured by ethnicity; PERMANOVA p-value and R² are annotated per panel. (C) Forest plot of CAZyme gene families (top 20) with significant ethnicity × timepoint interaction (LMM, FDR < 0.05), with the adjacent cross-sectional significance heatmap. (D) Glycosaminoglycan-to-dietary fibre (GAG/DF) ratio, (E) Mucin-to-dietary-fibre (Mucin/DF) ratio (log10) and (F) CAZyme family richness at baseline and follow-up by ethnicity; differences tested with paired Wilcoxon tests.

To characterize ethnic differences in carbohydrate-degrading capacity, we profiled CAZyme families using Cayman.^36^ CAZyme composition differed between Dutch and South-Asian Surinamese participants at both timepoints, with an R² of 0.048 at baseline and 0.037 at follow-up (Canberra distances; **Figure 4B**). Of 466 CAZyme families, 290 showed significant ethnic differences at baseline (q < 0.05) and 292 at follow-up (q < 0.05). At baseline, Dutch-and South-Asian Surinamese-enriched families were balanced (149 and 141 families, respectively); at follow-up, the balance shifted toward South-Asian Surinamese enrichment (121 higher in Dutch, 171 higher in South-Asian Surinamese). CAZyme families more abundant in Dutch participants were broad and included glycosyltransferases and plant-cell-wall degraders such as GH5 and GH11, while the CAZyme families that were more abundant in South-Asian Surinamese participants included α-glucan- and sucrose-active enzymes (several GH13; GH70 and GH68). Linear mixed models identified 50 families with a significant ethnicity × timepoint interaction, of which convergence was the dominant pattern (20 of 50; **Figure 4C; Supplementary Figure 18**). The most pronounced differential change occurred in three dietary-fibre-degrading families targeting hemicellulose and mixed-linkage β-glucans: GH5_37 (active on β-glucan, xyloglucan, glucomannan and lichenin), GH26 (a β-mannanase targeting β-mannan, galactomannan and lichenin), and CBM57 (a malectin-like carbohydrate-binding module). GH5_37 and GH26 were more abundant in Dutch participants at baseline and converged over time as South-Asian Surinamese participants increased these families relatively more (GH5_37: q = 0.0002; GH26: q = 0.006), while CBM57 showed no baseline difference but was higher abundant in South-Asian Surinamese at follow-up (q = 0.011) (**Figure 4C; Supplementary Figure 18**). All three lost significance after adjustment for baseline dietary intake (p = 0.001–0.03, q > 0.05); five families (GH5_2, GH5, CE9, GH13_16, GT51) remained significant after dietary adjustment.

To assess the balance between host glycan-targeting and dietary fibre (DF)-degrading enzyme capacity, we calculated mucin/DF and glycosaminoglycan (GAG)/DF ratios, following the annotations of Ducarmon et al.^36^ The GAG/DF ratio was significantly higher in Dutch compared to South-Asian Surinamese participants at both timepoints (baseline: p = 0.003, follow-up: p = 0.006; **Figure 4D**), and increased significantly over time in both groups similarly (ethnicity × timepoint: p = 0.49). The mucin/DF ratio decreased over time in both groups, and did not differ by ethnicity at either timepoint (**Figure 4E**). To determine which component drove the difference, we compared DF, GAG and mucin abundance separately, expressed relative to total CAZyme abundance, since total CAZyme abundance was higher in South-Asian Surinamese participants (CAZyme reads 3.94% vs 3.56%, p = 3 ×10^−18^). DF and mucin did not differ by ethnicity, while GAG was lower in South-Asian Surinamese participants (p = 0.001). The lower GAG/DF ratio is therefore attributable to relatively reduced GAG-targeting capacity. CAZyme family richness was lower in South-Asian Surinamese participants at both timepoints (median 337 versus 360 families at baseline: p = 1.02 ×10^−36^, follow-up: p = 3.45 ×10^−30^; **Figure 4F**). Thus, South-Asian Surinamese participants carried a smaller CAZyme repertoire at higher relative abundance. Higher GAG/DF ratio correlated with more favourable cardiometabolic parameters, such as higher HDL and lower waist-hip ratio and diastolic blood pressure (ρ = -0.13 to 0.14, p = 0.002–0.008; **Supplementary Figure 19**). However, these correlations lost significance after multiple testing correction.

### Total gut resistome burden declines in both ethnic groups

In the shotgun metagenomic subset, we analysed ARGs using annotation of MAGs with AMRFinderPlus and quantified ARGs in the sequencing reads. The total ARG burden in the gut microbiome declined significantly from baseline to follow-up across the full cohort (p = 2.6×10^−15^; **Figure 5A**). This decline was unchanged after adjusting for sequencing depth and for the number of genes predicted in each participant’s assemblies (p = 3.5×10^−15^) and in participants with similar sequencing depth at both timepoints (n = 371, p = 4.0×10^−15^). A decline in beta-lactam and tetracycline class ARGs accounted for the largest proportion of the decline (74.5%). In line, sixteen genes that changed significantly over time in the total population included multiple beta-lactam and tetracycline genes, such as *blaR1* (beta-lactam; q = 1.2×10^−13^), *tet(Q)* (tetracycline; q = 1.0×10^−8^) and *cfxA6* (beta-lactam; q = 4.1×10^−7^).

**Figure 5.**
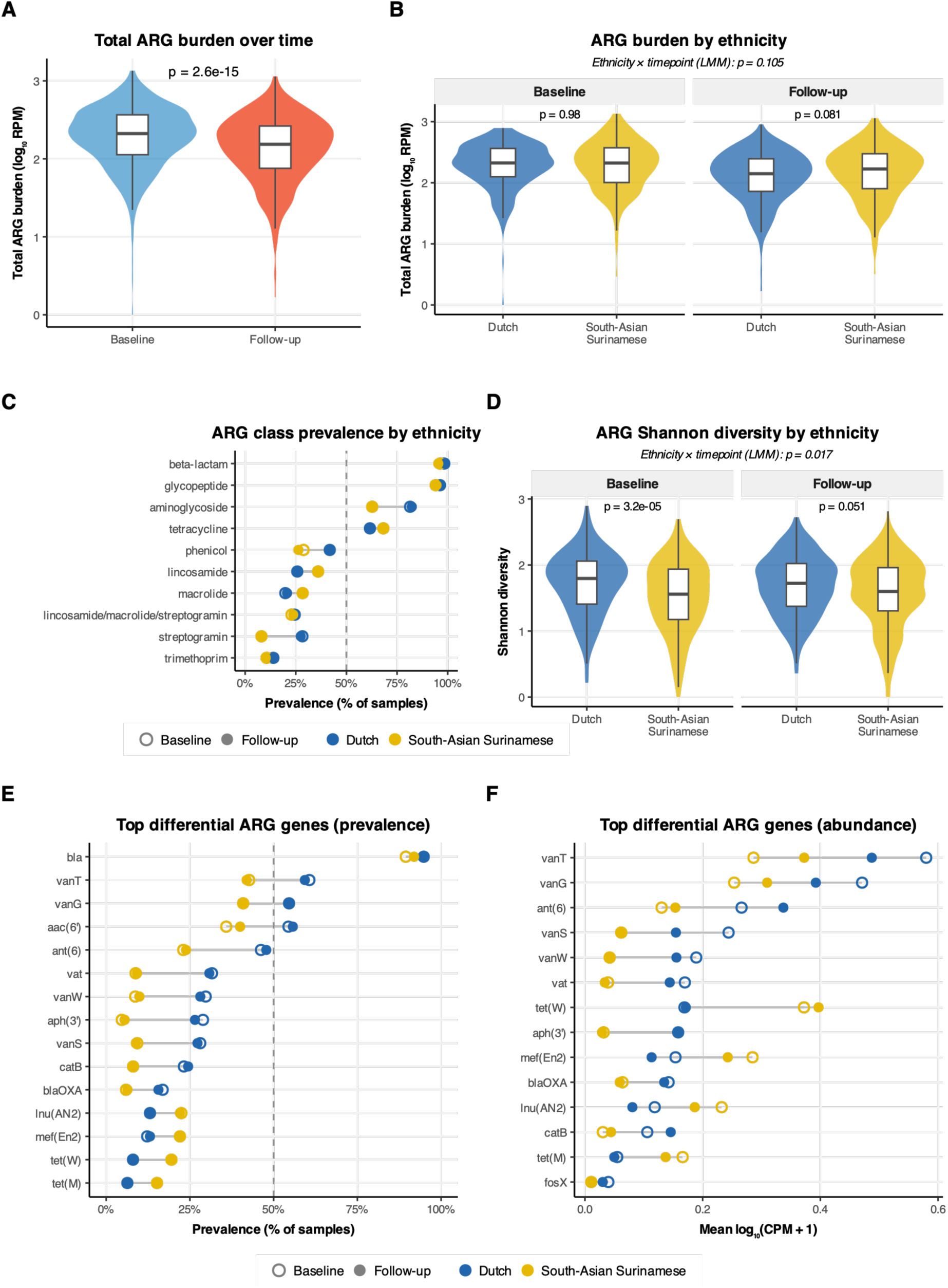
Antimicrobial resistance gene (ARG) dynamics in the HELIUS cohort. (A) Violin and boxplot of total ARG burden (RPM, log10-transformed) at baseline and follow-up across all participants. Overall change was tested with a linear mixed model (LMM). (B) Total ARG burden at baseline and follow-up stratified by ethnicity (Dutch vs. South-Asian Surinamese). Differences tested with Mann-Whitney U tests. (C) Dumbbell plot of ARG class prevalence (% of samples with at least one gene from the class) for the ten most prevalent resistance classes, by ethnicity and timepoint. (D) ARG Shannon diversity by ethnicity at baseline and follow-up; differences tested with Mann-Whitney U tests. (E) Gene-level dumbbell plot showing prevalence of 15 ARGs with significant ethnic differences at baseline (FDR < 0.05, chi-squared test). Prevalence is shown at baseline and follow-up for both ethnicities. (F) Dumbbell plot of mean log10(CPM + 1) abundance for the 14 ARGs with significant baseline ethnic differences (FDR < 0.05, linear regression). *blaOXA* and *fosX* lost significance after adjustment for sequencing depth and total predicted genes.

At baseline and follow-up, the total ARG burden did not differ between Dutch and South-Asian Surinamese participants, and the decline over time was comparable between groups (ethnicity x timepoint interaction p = 0.105; **Figure 5B**). At the class level, aminoglycoside (81.4% vs 62.7%, q < 0.001), streptogramin (28.3% vs 8.1%, q < 0.001), and phenicol (41.8% vs 28.8%, q = 0.019) ARGs were more prevalent in Dutch than South-Asian Surinamese participants. These class-level differences were largely stable across the follow-up period (**Figure 5C**). ARG Shannon diversity was higher in Dutch than South-Asian Surinamese participants at baseline (p = 3.2×10^−5^), but the groups converged at the follow-up timepoint (β_ethnicity×time_ = 0.093, 95% CI 0.017–0.170, p = 0.017; **Figure 5D**). This interaction remained significant after depth and total genes adjustment and after requiring at least 5 or 10 mapped reads per gene.

Several resistance genes showed ethnic differences at baseline. The aminoglycoside genes *aph(3’)* and *ant(C),* the streptogramin gene *vat*, and the vancomycin genes *vanW* and *vanS* were more prevalent and abundant in Dutch than South-Asian Surinamese participants (all q < 0.001; **Figure 5E-F**). Adjusting for sequencing depth and total predicted genes attenuated these differences; 12 of the 14 genes with differential abundance at baseline remained significant, while *blaOXA* and *fosX* lost significance. For prevalence, 11 of 15 remained significant, while *vanT*, *vanG*, *aac(6’)* and *bla* lost significance. Although most ethnic differences in gene prevalence persisted into follow-up, two vancomycin gene abundances (*vanT*, *vanG*) converged between groups over time (q = 0.002 and q = 0.010, respectively; **Supplementary Figure 20**), and both remained significant after adjustment. Thus, ethnic differences in ARG composition were present at baseline, and the decline over time led to partial convergence of vancomycin gene abundance and ARG diversity at follow-up.

## Discussion

Ethnic differences in gut microbiome composition persisted across compositional, functional, strain, and assembly-level layers despite a shared urban environment. In the 16S data, we observed minimal convergence between ethnicities in the alpha and beta diversity analyses over six years. The observed rate of R² decline for ethnicity, assuming exponential decline, corresponds to a half-life of approximately 20 years. This aligns with previous evidence from the HELIUS cohort showing that microbiome differences remain detectable across generations of migration, and evidence from Irish Travellers showing that a non-industrialized-like microbiome persists decades after enforced settlement and lifestyle change.^15,59^ The latter study also emphasized the importance of non-dietary factors in the persistence of these differences, including housing conditions, household crowding, and animal ownership.^15^ The rapid Westernization reported in recently arrived immigrants reflects an acute post-migration transition;^13^ our cohort, on average resident in Amsterdam for decades, instead shows what remains long after the acute transition. The forces sustaining these residual differences -including housing and animal-contact factors implicated in Irish Travellers -were not measured here and remain to be established.

Higher baseline alpha diversity was associated with less compositional change over time, consistent with the macroecological principle that more diverse communities resist perturbation.^60^ Several baseline cardiometabolic and medication-related factors predicted greater microbiome change over time, including diabetes and metabolic syndrome, and metformin and PPI use. Disentangling disease from treatment effects is difficult here: multiple medication classes showed associations with compositional shifts, and metformin in particular has well-documented effects on gut microbiota composition.^61^ Our findings complement our previously published analysis in this cohort, in which baseline microbiome composition predicted incident cardiometabolic outcomes.^5^ That analysis investigated whether the microbiome predicts disease; the current paper assessed whether disease changes how the microbiome evolves. While most prior work has examined the microbiome as a predictor of cardiometabolic risk,^62–65^ disease and its clinical management are equally likely to reshape microbiome composition over time.

The shotgun metagenomics analyses in a subgroup of Dutch and South-Asian Surinamese participants showed the same persistence of differences. A few species including *Alistipes putredinis* showed differential change over time, which often led to convergence. In *Alistipes putredinis,* convergence occurred, mostly through new acquisition by South-Asian Surinamese participants. Clade structure was similar between groups, with one clade more prevalent in Dutch participants. Strain turnover in this species was comparable between ethnic groups, and within-host microdiversity increased at a similar rate in both. Dutch participants did carry more heterogeneous populations at both timepoints, concordant with their higher community diversity, which is associated with higher intra-species polymorphism.^66^ Thus, the persistence of ethnic differences over six years was driven by species presence and abundance rather than by strain-level dynamics, which were comparable between groups.

Functional dissimilarity declined at the same rate as taxonomic dissimilarity (ethnicity R² 0.047 to 0.037 for species and 0.048 to 0.037 for CAZymes). Approximately two-thirds of the CAZyme families detected differed between ethnic groups at both baseline and follow-up. Three hemicellulose- and β-glucan-targeting families shifted toward a relatively higher increase or smaller decrease in South-Asian Surinamese compared to Dutch participants, likely driven by differences in dietary intake. Ducarmon et al. found higher GAG/DF and mucin/DF ratios in high-income-setting metagenomes, consistent with diets high in animal products and low in fibre.^36^ Within one high-income setting, South-Asian Surinamese participants had a lower GAG/DF ratio and lower CAZyme family richness than Dutch participants. The ratio difference was small and was driven by reduced GAG-targeting capacity, not by higher fibre-degrading capacity, therefore the lower fibre intake does not seem to explain this difference. Colonic GAGs are derived from host proteoglycans and from animal-derived dietary sources. Additionally, GAG-targeting loci are concentrated in *Bacteroides*, so the ratio could reflect compositional differences or GAG availability. The ratio and substrate class differences were of similar magnitude at baseline and follow-up, while taxonomic and functional dissimilarity both declined.

Total ARG burden declined significantly in both groups over the study period, driven primarily by beta-lactam and tetracycline resistance genes, consistent with the broader trend of falling antibiotic prescribing rates during this period in the Netherlands.^67^ Within HELIUS, antibiotic prescription rates did not differ significantly between Dutch and South-Asian Surinamese participants after adjustment for health status, with disease burden rather than ethnicity being the primary prescription driver,^68^ and GP and specialist contact rates were comparable between groups.^69^ However, the resistome composition showed limited convergence: ARG diversity converged between groups, while class-level differences were largely stable. Vancomycin resistance genes (*vanT*, *vanG*) showed the most consistent convergence. Antibiotic use and gut microbial composition are the two strongest population-level predictors of individual gut microbial ARG load;^70^ the similar antibiotic exposure between groups therefore points to the persistent compositional differences as the likely explanation for the retained resistome distinctions.

## Conclusions

Ethnic differences in gut microbiome composition were stable over six years in a shared Westernized urban environment, with limited convergence across compositional, functional, strain-level and MAG layers. Within individuals, change was driven mainly by type 2 diabetes and medication use rather than by ethnicity. Convergence, where it occurred, had a half-life of roughly two decades, so ethnic differences in the gut microbiome are likely to persist in multi-ethnic urban populations for the foreseeable future, and cohorts recruited in such settings should account for them.

## Data availability

The Amsterdam UMC, located at AMC in Amsterdam, The Netherlands, owns the HELIUS data. Researchers interested in accessing this data can submit proposals to the HELIUS Executive Board via http://www.heliusstudy.nl/en/researchers/collaboration. The HELIUS Executive Board will evaluate proposals for alignment with the study’s overall objectives, ethical approvals, and informed consent forms. The 16S microbiota sequencing data and shotgun metagenomics sequencing data have been deposited in a repository of the European Genome-Phenome Archive (https://ega-archive.org/studies/EGAS00001002969).

## Code availability

We used a number of Nextflow workflows for processing the 16S and shotgun data, that are all publicly available in Github repositories: <u>vsearchpipeline</u>, <u>metagenomicspipeline</u>, <u>strainflow</u>, <u>caymanflow</u>, <u>annotationpipeline</u>, <u>phylotree</u>, <u>amrfinderflow</u>. The code for the downstream statistics and visualizations was published in a <u>separate repository</u>. All launch scripts for the workflows processing of the sequencing data can also be found there.

## Acknowledgements

We would like to thank the Amsterdam UMC Biobank for their support with sample storage and the participants, research nurses and HELIUS staff for their help in data collection.

## Author contributions

B.J.H.vd.B., A.Z. and M.Nie. were responsible for the design and management of the HELIUS cohort. U.B., B.J.H.V., H.H., B.J.H.vd.B., M.Nic., N.H., E.E., T.N. and M.Nie. designed the current project. B.J.H.V. was responsible for the bioinformatic workflows and analyses. B.J.H.V. and M.Nie. wrote the first draft of the manuscript. All authors contributed to the manuscript. All authors had full access to all the data in the study and had final responsibility for the decision to submit for publication.

## References

1. Tillin, T. et al. The Relationship Between Metabolic Risk Factors and Incident Cardiovascular Disease in Europeans, South Asians, and African Caribbeans. J Am Coll Cardiol 61, 1777–1786 (2013).

2. Boateng, D. et al. Migration and Cardiovascular Disease Risk Among Ghanaian Populations in Europe: The RODAM Study (Research on Obesity and Diabetes Among African Migrants). Circ Cardiovasc Qual Outcomes 10, e004013 (2017).

3. Agyemang, C. et al. Obesity and type 2 diabetes in sub-Saharan Africans – Is the burden in today’s Africa similar to African migrants in Europe? The RODAM study. BMC Med 14, 166 (2016).

4. Deschasaux, M. et al. Depicting the composition of gut microbiota in a population with varied ethnic origins but shared geography. Nature medicine 24, 1526–1531 (2018).

5. Verhaar, B. J. H., Bouwmeester, T. A., Galenkamp, H., van den Born, B.-J. H. & Nieuwdorp, M. Specific gut microbes are associated with the incidence of cardiometabolic disease in the HELIUS cohort. npj Biofilms Microbiomes 12, 83 (2026).

6. Sonnenburg, E. D. et al. Diet-induced extinctions in the gut microbiota compound over generations. Nature 529, 212–215 (2016).

7. Rothschild, D. et al. Environment dominates over host genetics in shaping human gut microbiota. Nature 555, 210–215 (2018).

8. Mackenbach, J. D. et al. Socioeconomic and ethnic differences in the relation between dietary costs and dietary quality: the HELIUS study. Nutr J 18, 21 (2019).

9. Weersma, R. K., Zhernakova, A. & Fu, J. Interaction between drugs and the gut microbiome. Gut 10.1136/gutjnl-2019-320204 (2020) doi:10.1136/gutjnl-2019-320204.

10. Shah, S. et al. Physical activity-induced alterations of the gut microbiota are BMI dependent. The FASEB Journal 37, e22882 (2023).

11. Goodrich, J. K. et al. Human Genetics Shape the Gut Microbiome. Cell 159, 789–799 (2014).

12. Kurilshikov, A. et al. Large-scale association analyses identify host factors influencing human gut microbiome composition. Nat Genet 53, 156–165 (2021).

13. Vangay, P. et al. US Immigration Westernizes the Human Gut Microbiome. Cell 175, 962–972.e10 (2018).

14. Othman, S. I., Fertig, A., Trofholz, A. & Berge, J. M. How time in the US and race/ethnicity shape food parenting practices and child diet quality. Appetite 171, 105870 (2022).

15. Keohane, D. M. et al. Microbiome and health implications for ethnic minorities after enforced lifestyle changes. Nat Med 26, 1089–1095 (2020).

16. Halfvarson, J. et al. Dynamics of the human gut microbiome in inflammatory bowel disease. Nat Microbiol 2, 17004 (2017).

17. Frost, F. et al. Long-term instability of the intestinal microbiome is associated with metabolic liver disease, low microbiota diversity, diabetes mellitus and impaired exocrine pancreatic function. 10.1136/gutjnl-2020-322753 (2021) doi:10.1136/gutjnl-2020-322753.

18. Snijder, M. B., et al. Cohort profile: the healthy life in an urban setting (HELIUS) study in Amsterdam, the Netherlands. BMJ open 7, e017873 (2017).

19. Stronks, K., Kulu-Glasgow, I. & Agyemang, C. The utility of ‘country of birth’ for the classification of ethnic groups in health research: the Dutch experience. Ethnicity & Health 14, 255–269 (2009).

20. Galenkamp, H. et al. The Healthy Life in an Urban Setting (HELIUS) study in Amsterdam, The Netherlands: cohort update 2024 and key findings. 2024.07.16.24310494 Preprint at 10.1101/2024.07.16.24310494 (2024).

21. Elm, E. von et al. The Strengthening the Reporting of Observational Studies in Epidemiology (STROBE) statement: guidelines for reporting observational studies. The Lancet 370, 1453–1457 (2007).

22. Alberti, K. G. M. M. et al. Harmonizing the Metabolic Syndrome. Circulation 120, 1640–1645 (2009).

23. Nicolaou, M. et al. Validation of the SQUASH Physical Activity Questionnaire in a Multi-Ethnic Population: The HELIUS Study. PLOS ONE 11, e0161066 (2016).

24. Beukers, M. H. et al. Development of the HELIUS food frequency questionnaires: ethnic-specific questionnaires to assess the diet of a multiethnic population in The Netherlands. European journal of clinical nutrition 69, 579–584 (2015).

25. Rognes, T., Flouri, T., Nichols, B., Quince, C. & Mahé, F. VSEARCH: a versatile open source tool for metagenomics. PeerJ 4, e2584 (2016).

26. Callahan, B. J. et al. DADA2: High-resolution sample inference from Illumina amplicon data. Nat Methods 13, 581–583 (2016).

27. McMurdie, P. J. & Holmes, S. Phyloseq: An R Package for Reproducible Interactive Analysis and Graphics of Microbiome Census Data. PLoS ONE 10.1371/journal.pone.0061217 (2013) doi:10.1371/journal.pone.0061217.

28. Davis, N. M., Proctor, D. M., Holmes, S. P., Relman, D. A. & Callahan, B. J. Simple statistical identification and removal of contaminant sequences in marker-gene and metagenomics data. Microbiome 6, 226 (2018).

29. Di Tommaso, P. et al. Nextflow enables reproducible computational workflows. Nat Biotechnol 35, 316–319 (2017).

30. Chen, S., Zhou, Y., Chen, Y. & Gu, J. fastp: an ultra-fast all-in-one FASTQ preprocessor. Bioinformatics 34, i884–i890 (2018).

31. Langmead, B. & Salzberg, S. L. Fast gapped-read alignment with Bowtie 2. Nat Methods 9, 357–359 (2012).

32. Shen, W., Sipos, B. & Zhao, L. SeqKit2: A Swiss army knife for sequence and alignment processing. iMeta 3, e191 (2024).

33. Blanco-Míguez, A. et al. Extending and improving metagenomic taxonomic profiling with uncharacterized species using MetaPhlAn 4. Nat Biotechnol 41, 1633–1644 (2023).

34. Beghini, F. et al. Integrating taxonomic, functional, and strain-level profiling of diverse microbial communities with bioBakery 3. eLife 10, e65088.

35. Valles-Colomer, M. et al. The person-to-person transmission landscape of the gut and oral microbiomes. Nature 614, 125–135 (2023).

36. Ducarmon, Q. R., et al. Cayman enables large-scale analysis of gut microbiome carbohydrate-active enzyme repertoires. Nat Microbiol 1–15 (2026) doi:10.1038/s41564-026-02318-2.

37. Krakau, S., Straub, D., Gourlé, H., Gabernet, G. & Nahnsen, S. nf-core/mag: a best-practice pipeline for metagenome hybrid assembly and binning. NAR Genom Bioinform 4, lqac007 (2022).

38. Ewels, P. A. et al. The nf-core framework for community-curated bioinformatics pipelines. Nat Biotechnol 38, 276–278 (2020).

39. Li, D. et al. MEGAHIT v1.0: A fast and scalable metagenome assembler driven by advanced methodologies and community practices. Methods 102, 3–11 (2016).

40. Kang, D. D. et al. MetaBAT 2: an adaptive binning algorithm for robust and efficient genome reconstruction from metagenome assemblies. PeerJ 7, e7359 (2019).

41. Wu, Y.-W., Simmons, B. A. & Singer, S. W. MaxBin 2.0: an automated binning algorithm to recover genomes from multiple metagenomic datasets. Bioinformatics 32, 605–607 (2016).

42. Alneberg, J. et al. Binning metagenomic contigs by coverage and composition. Nat Methods 11, 1144–1146 (2014).

43. Sieber, C. M. K. et al. Recovery of genomes from metagenomes via a dereplication, aggregation and scoring strategy. Nat Microbiol 3, 836–843 (2018).

44. Chklovski, A., Parks, D. H., Woodcroft, B. J. & Tyson, G. W. CheckM2: a rapid, scalable and accurate tool for assessing microbial genome quality using machine learning. Nat Methods 20, 1203–1212 (2023).

45. Chaumeil, P.-A., Mussig, A. J., Hugenholtz, P. & Parks, D. H. GTDB-Tk: a toolkit to classify genomes with the Genome Taxonomy Database. Bioinformatics 36, 1925–1927 (2020).

46. Schwengers, O. et al. Bakta: rapid and standardized annotation of bacterial genomes via alignment-free sequence identification. Microbial Genomics 7, 000685 (2021).

47. Buchfink, B., Reuter, K. & Drost, H.-G. Sensitive protein alignments at tree-of-life scale using DIAMOND. Nat Methods 18, 366–368 (2021).

48. Liu, B., Zheng, D., Zhou, S., Chen, L. & Yang, J. VFDB 2022: a general classification scheme for bacterial virulence factors. Nucleic Acids Res 50, D912–D917 (2022).

49. Tonkin-Hill, G. et al. Producing polished prokaryotic pangenomes with the Panaroo pipeline. Genome Biol 21, 180 (2020).

50. Wong, T. K. F. et al. IQ-TREE 3: Phylogenomic Inference Software using Complex Evolutionary Models. https://ecoevorxiv.org/repository/view/8916/ (2025).

51. nf-core/funcscan: v3.0.0 -French Chocolatine -2025-10-04. https://zenodo.org/records/17267739.

52. Steinegger, M. & Söding, J. MMseqs2 enables sensitive protein sequence searching for the analysis of massive data sets. Nat Biotechnol 35, 1026–1028 (2017).

53. Larralde, M. Pyrodigal: Python bindings and interface to Prodigal, an efficient method for gene prediction in prokaryotes. Journal of Open Source Software 7, 4296 (2022).

54. Feldgarden, M. et al. AMRFinderPlus and the Reference Gene Catalog facilitate examination of the genomic links among antimicrobial resistance, stress response, and virulence. Sci Rep 11, 12728 (2021).

55. Ugarcina Perovic, S., et al. argNorm: normalization of antibiotic resistance gene annotations to the Antibiotic Resistance Ontology (ARO). Bioinformatics 41, btaf173 (2025).

56. Li, W. & Godzik, A. Cd-hit: a fast program for clustering and comparing large sets of protein or nucleotide sequences. Bioinformatics 22, 1658–1659 (2006).

57. Vasimuddin, Md., Misra, S., Li, H. & Aluru, S. Efficient Architecture-Aware Acceleration of BWA-MEM for Multicore Systems. in 2019 IEEE International Parallel and Distributed Processing Symposium (IPDPS) 314–324 (2019). doi:10.1109/IPDPS.2019.00041.

58. Willett, W., Howe, G. & Kushi, L. Adjustment for total energy intake in epidemiologic studies. The American Journal of Clinical Nutrition 65, 1220S–1228S (1997).

59. van der Vossen, E. W. J. et al. Gut microbiome transitions across generations in different ethnicities in an urban setting—the HELIUS study. Microbiome 11, 99 (2023).

60. Lozupone, C. A., Stombaugh, J. I., Gordon, J. I., Jansson, J. K. & Knight, R. Diversity, stability and resilience of the human gut microbiota. Nature 489, 220–230 (2012).

61. Forslund, K. et al. Disentangling type 2 diabetes and metformin treatment signatures in the human gut microbiota. Nature 10.1038/nature15766 (2015) doi:10.1038/nature15766.

62. Boulund, U. et al. Gut microbiome associations with host genotype vary across ethnicities and potentially influence cardiometabolic traits. Cell host & microbe 30, 1464–1480. e6 (2022).

63. He, G., Cao, Y., Wang, H. & Lv, X. Causal effects of gut microbiome on hypertension: a Mendelian randomization study. Front. Microbiol. 14, (2023).

64. Teng, D. et al. Causality of the gut microbiome and atherosclerosis-related lipids: a bidirectional Mendelian Randomization study. BMC Cardiovasc Disord 24, 138 (2024).

65. Jia, J. et al. Assessment of causal direction between gut microbiota-dependent metabolites and cardiometabolic health: A bidirectional mendelian randomization analysis. Diabetes 68, 1747–1755 (2019).

66. Madi, N., Chen, D., Wolff, R., Shapiro, B. J. & Garud, N. R. Community diversity is associated with intra-species genetic diversity and gene loss in the human gut microbiome. eLife 12, e78530 (2023).

67. van de Pol, A. C. et al. Impact of the COVID-19 Pandemic on Antibiotic Prescribing for Common Infections in The Netherlands: A Primary Care-Based Observational Cohort Study. Antibiotics 10, 196 (2021).

68. Schuts, E. C. et al. Knowledge and use of antibiotics in six ethnic groups: the HELIUS study. Antimicrob Resist Infect Control 8, 200 (2019).

69. Cassidy, M., Agyemang, C., Galenkamp, H., van Charante, E. M. & Chilunga, F. P. Healthcare utilisation among migrants in the Netherlands’ unique hybrid healthcare system: the HELIUS study. BMJ Glob Health 11, e020024 (2026).

70. Pärnänen, K. et al. Variation and prognostic potential of the gut antibiotic resistome in the FINRISK 2002 cohort. Nat Commun 16, 5963 (2025).

