## Supplementary Tables and Figures for "Ethnic differences in gut microbiota composition and functional capacity persist over six years in the HELIUS multi-ethnic longitudinal cohort"

### Supplementary Table 1: HELIUS population with 16S microbiome data

| Characteristic | Baseline | Follow-up | p-value | Missing (%) |
|---|---|---|---|---|
| **n** | 1,903 | 1,903 | | |
| **Age, years** | 52.47 ± 10.07 | 58.82 ± 10.29 | **<0.001** | 0.0 |
| **Sex, female** | 995 (52.3%) | — | — | 0.0 |
| **Ethnicity** | | | — | 0.0 |
| *Dutch* | 666 (35.0%) | — | — | |
| *South-Asian Surinamese* | 411 (21.6%) | — | — | |
| *African Surinamese* | 398 (20.9%) | — | — | |
| *Ghanaian* | 141 (7.4%) | — | — | |
| *Turkish* | 92 (4.8%) | — | — | |
| *Moroccan* | 169 (8.9%) | — | — | |
| *Other* | 26 (1.4%) | — | — | |
| **Follow-up time, years** | — | 6.31 ± 1.27 | — | 0.0 |
| **Residence duration, years** | 31.33 ± 10.04 | — | — | — |
| **BMI, kg/m²** | 26.72 ± 4.47 | 27.10 ± 4.59 | **0.010** | 0.0 |
| **Non-smoker** | 1,560 (82.5%) | 1,632 (85.8%) | **0.006** | 0.3 |
| **Alcohol consumption** | | | **<0.001** | 11.5 |
| *Low* | 1,296 (68.5%) | 1,169 (79.1%) | | |
| *Moderate* | 438 (23.2%) | 254 (17.2%) | | |
| *High* | 158 (8.4%) | 55 (3.7%) | | |
| **No alcohol use** | 673 (35.5%) | 635 (33.4%) | 0.175 | 0.3 |
| **Diabetes mellitus** | 207 (10.9%) | 304 (16.1%) | **<0.001** | 0.3 |
| **Hypertension** | 814 (42.8%) | 911 (47.9%) | **0.002** | 0.0 |
| **Metabolic syndrome** | 729 (38.4%) | 823 (43.4%) | **0.002** | 0.3 |
| **Dyslipidaemia** | 1,243 (65.3%) | 1,400 (73.6%) | **<0.001** | 0.0 |
| **Systolic BP, mmHg** | 130.18 ± 17.59 | 129.84 ± 17.99 | 0.554 | 0.0 |
| **Diastolic BP, mmHg** | 80.79 ± 10.22 | 79.02 ± 10.20 | **<0.001** | 0.1 |
| **Total cholesterol, mmol/L** | 5.13 ± 1.00 | 5.26 ± 1.09 | **<0.001** | 0.3 |
| **LDL cholesterol, mmol/L** | 3.16 ± 0.89 | 3.18 ± 0.96 | 0.669 | 0.9 |
| **Triglycerides, mmol/L** | 0.89 [0.61, 1.26] | 0.94 [0.67, 1.34] | **<0.001** | 0.3 |
| **HbA1c, mmol/mol** | 40.07 ± 8.40 | 40.48 ± 9.76 | 0.167 | 1.5 |
| **PPI use** | 213 (11.2%) | 311 (16.3%) | **<0.001** | 0.0 |
| **Metformin use** | 154 (8.1%) | 206 (10.8%) | **0.005** | 0.0 |
| **Statin use** | 242 (12.7%) | — | — | 50.0 |
| **Antihypertensive medication** | 420 (22.1%) | 591 (31.1%) | **<0.001** | 0.0 |
| **Glucose-lowering drugs** | 169 (8.9%) | 230 (12.1%) | **0.001** | 0.0 |
| **Psychotropic medication** | 115 (6.0%) | — | — | 50.0 |
| **Corticosteroids** | 96 (5.0%) | 119 (6.3%) | 0.122 | 0.0 |

*Continuous variables are reported as mean (SD) or median [IQR] for non-normally distributed variables (triglycerides); baseline and follow-up were compared using Student's t-test or Wilcoxon rank-sum test, respectively. Categorical variables are reported as n (%) and compared with chi-squared tests. Some variables were not available at follow-up, or redundant because they were static, these fields are empty.*

### Supplementary Table 2: HELIUS population with 16S microbiome data stratified for ethnicity

| | African Surinamese | | Dutch | | Ghanaian | |
|---|---|---|---|---|---|---|
| **Characteristic** | **Baseline** | **Follow-up** | **Baseline** | **Follow-up** | **Baseline** | **Follow-up** |
| **n** | 398 | 398 | 666 | 666 | 141 | 141 |
| **Age, years** | 53.68 ± 9.19 | 60.09 ± 9.51 | 53.53 ± 10.92 | 59.94 ± 11.11 | 49.11 ± 8.76 | 55.80 ± 8.92 |
| **Female sex** | 243 (61.1) | — | 321 (48.2) | — | 77 (54.6) | — |
| **Follow-up time, years** | 6.38 ± 1.39 | — | 6.37 ± 1.36 | — | 6.65 ± 0.89 | — |
| **Residence duration, years** | 33.15 ± 10.60 | — | — | — | 20.05 ± 7.58 | — |
| **BMI, kg/m²** | 28.27 ± 5.09 | 28.48 ± 5.01 | 25.04 ± 3.84 | 25.30 ± 4.00 | 27.56 ± 3.82 | 28.44 ± 3.90 |
| **Current non-smoker** | 306 (77.3) | 312 (78.4) | 552 (83.0) | 577 (86.6) | 135 (97.1) | 140 (99.3) |
| **Alcohol consumption** | | | | | | |
| Low | 327 (82.6) | 251 (88.1) | 235 (35.3) | 213 (48.7) | 121 (87.7) | 121 (91.7) |
| Moderate | 53 (13.4) | 29 (10.2) | 313 (47.1) | 182 (41.6) | 15 (10.9) | 9 (6.8) |
| High | 16 (4.0) | 5 (1.8) | 117 (17.6) | 42 (9.6) | 2 (1.4) | 2 (1.5) |
| **No alcohol use** | 139 (35.0) | 105 (26.4) | 50 (7.5) | 60 (9.0) | 71 (51.4) | 70 (49.6) |
| **Diabetes mellitus** | 40 (10.1) | 59 (14.9) | 28 (4.2) | 38 (5.7) | 15 (10.6) | 26 (18.4) |
| **Hypertension or BP medication** | 226 (56.9) | 238 (59.8) | 224 (33.6) | 256 (38.4) | 86 (61.0) | 91 (64.5) |
| **Metabolic syndrome** | 145 (36.5) | 182 (45.8) | 190 (28.6) | 213 (32.1) | 48 (34.0) | 59 (41.8) |
| **Dyslipidaemia** | 248 (62.3) | 279 (70.1) | 430 (64.6) | 488 (73.3) | 82 (58.2) | 95 (67.4) |
| **Systolic BP, mmHg** | 134.40 ± 17.89 | 132.92 ± 18.03 | 126.72 ± 16.80 | 126.59 ± 16.96 | 136.61 ± 17.90 | 137.53 ± 18.87 |
| **Diastolic BP, mmHg** | 83.16 ± 10.29 | 80.41 ± 10.33 | 78.58 ± 9.97 | 77.83 ± 9.94 | 84.32 ± 10.92 | 83.52 ± 10.71 |
| **PPI use** | 43 (10.8) | 61 (15.3) | 50 (7.5) | 79 (11.9) | 9 (6.4) | 12 (8.5) |
| **Metformin use** | 32 (8.0) | 37 (9.3) | 13 (2.0) | 17 (2.6) | 11 (7.8) | 13 (9.2) |
| **Statin use** | 36 (9.0) | — | 61 (9.2) | — | 15 (10.6) | — |
| **Antihypertensive use** | 130 (32.7) | 169 (42.5) | 92 (13.8) | 143 (21.5) | 44 (31.2) | 60 (42.6) |
| **Glucose-lowering drugs** | 36 (9.0) | 43 (10.8) | 15 (2.3) | 21 (3.2) | 12 (8.5) | 15 (10.6) |
| **Psychotropic medication** | 16 (4.0) | — | 37 (5.6) | — | 5 (3.5) | — |
| **Corticosteroid use** | 21 (5.3) | 28 (7.0) | 28 (4.2) | 34 (5.1) | 4 (2.8) | 4 (2.8) |
| **Total cholesterol, mmol/L** | 5.06 ± 0.96 | 5.20 ± 1.08 | 5.33 ± 0.99 | 5.52 ± 1.09 | 5.06 ± 1.01 | 5.24 ± 0.95 |
| **LDL cholesterol, mmol/L** | 3.12 ± 0.88 | 3.15 ± 0.97 | 3.24 ± 0.90 | 3.31 ± 0.95 | 3.12 ± 0.94 | 3.23 ± 0.88 |
| **Triglycerides, mmol/L** | 0.79 [0.56, 1.07] | 0.84 [0.58, 1.13] | 0.88 [0.59, 1.20] | 0.94 [0.65, 1.36] | 0.59 [0.45, 0.84] | 0.71 [0.53, 0.89] |
| **HbA1c, mmol/mol** | 40.94 ± 7.99 | 41.33 ± 10.74 | 36.85 ± 4.42 | 36.96 ± 5.23 | 40.39 ± 10.28 | 40.82 ± 10.68 |

| | Moroccan | | South-Asian Surinamese | | Turkish | |
|---|---|---|---|---|---|---|
| **Characteristic** | **Baseline** | **Follow-up** | **Baseline** | **Follow-up** | **Baseline** | **Follow-up** |
| **n** | 169 | 169 | 411 | 411 | 92 | 92 |
| **Age, years** | 49.49 ± 9.94 | 55.14 ± 10.11 | 52.92 ± 9.46 | 59.44 ± 9.53 | 47.50 ± 8.47 | 53.25 ± 8.51 |
| **Female sex** | 72 (42.6) | — | 222 (54.0) | — | 46 (50.0) | — |
| **Follow-up time, years** | 5.64 ± 0.82 | — | 6.48 ± 1.19 | — | 5.72 ± 0.96 | — |
| **Residence duration, years** | 31.94 ± 8.78 | — | 33.41 ± 8.41 | — | 30.14 ± 8.08 | — |
| **BMI, kg/m²** | 28.13 ± 4.02 | 28.62 ± 4.22 | 26.67 ± 4.23 | 27.06 ± 4.24 | 28.49 ± 4.34 | 29.45 ± 4.67 |
| **Current non-smoker** | 151 (89.9) | 156 (92.3) | 325 (79.7) | 349 (84.9) | 71 (78.0) | 77 (83.7) |
| **Alcohol consumption** | | | | | | |
| Low | 160 (95.2) | 162 (97.6) | 355 (86.8) | 313 (91.5) | 82 (90.1) | 89 (96.7) |
| Moderate | 7 (4.2) | 4 (2.4) | 37 (9.0) | 24 (7.0) | 7 (7.7) | 3 (3.3) |
| High | 1 (0.6) | 0 (0.0) | 17 (4.2) | 5 (1.5) | 2 (2.2) | 0 (0.0) |
| **No alcohol use** | 153 (91.1) | 156 (92.9) | 188 (45.9) | 177 (43.1) | 63 (69.2) | 59 (64.1) |
| **Diabetes mellitus** | 24 (14.2) | 33 (19.5) | 91 (22.2) | 129 (31.9) | 7 (7.6) | 13 (14.1) |
| **Hypertension or BP medication** | 55 (32.5) | 62 (36.7) | 188 (45.7) | 223 (54.3) | 19 (20.7) | 27 (29.3) |
| **Metabolic syndrome** | 71 (42.0) | 68 (40.2) | 234 (57.2) | 257 (63.1) | 31 (33.7) | 32 (34.8) |
| **Dyslipidaemia** | 104 (61.5) | 115 (68.0) | 302 (73.5) | 337 (82.0) | 59 (64.1) | 66 (71.7) |
| **Systolic BP, mmHg** | 128.96 ± 17.81 | 128.97 ± 19.27 | 131.22 ± 17.10 | 130.95 ± 18.03 | 123.77 ± 15.96 | 124.39 ± 14.77 |
| **Diastolic BP, mmHg** | 79.00 ± 9.44 | 77.22 ± 9.89 | 81.70 ± 9.61 | 79.01 ± 10.03 | 79.71 ± 10.37 | 77.62 ± 9.86 |
| **PPI use** | 31 (18.3) | 30 (17.8) | 60 (14.6) | 106 (25.8) | 16 (17.4) | 17 (18.5) |
| **Metformin use** | 16 (9.5) | 19 (11.2) | 75 (18.2) | 107 (26.0) | 6 (6.5) | 9 (9.8) |
| **Statin use** | 21 (12.4) | — | 99 (24.1) | — | 7 (7.6) | — |
| **Antihypertensive use** | 19 (11.2) | 36 (21.3) | 118 (28.7) | 157 (38.2) | 9 (9.8) | 16 (17.4) |
| **Glucose-lowering drugs** | 18 (10.7) | 21 (12.4) | 80 (19.5) | 115 (28.0) | 6 (6.5) | 10 (10.9) |
| **Psychotropic medication** | 12 (7.1) | — | 29 (7.1) | — | 11 (12.0) | — |
| **Corticosteroid use** | 12 (7.1) | 9 (5.3) | 22 (5.4) | 38 (9.2) | 8 (8.7) | 5 (5.4) |
| **Total cholesterol, mmol/L** | 4.79 ± 0.91 | 4.96 ± 0.96 | 5.05 ± 1.02 | 5.00 ± 1.12 | 5.00 ± 1.01 | 5.31 ± 1.11 |
| **LDL cholesterol, mmol/L** | 2.98 ± 0.80 | 3.01 ± 0.82 | 3.15 ± 0.91 | 3.00 ± 1.00 | 3.14 ± 0.90 | 3.30 ± 1.00 |
| **Triglycerides, mmol/L** | 0.88 [0.64, 1.21] | 1.02 [0.76, 1.33] | 1.06 [0.76, 1.52] | 1.09 [0.81, 1.52] | 1.09 [0.70, 1.48] | 1.15 [0.83, 1.47] |
| **HbA1c, mmol/mol** | 40.46 ± 8.24 | 41.21 ± 10.57 | 44.45 ± 10.89 | 45.42 ± 11.85 | 38.89 ± 7.20 | 38.63 ± 7.69 |

*Continuous variables are presented as mean ± SD or median [IQR]; categorical variables are presented as n (%). Some variables were not available at follow-up, or redundant because they were static, these fields are empty.*

### Supplementary Table 3: HELIUS population with shotgun metagenomics data

| Characteristic | Baseline | Follow-up |
|---|---|---|
| **n** | **476** | **476** |
| **Age, years** | 51.41 ± 10.54 | 57.89 ± 10.73 |
| **Female sex** | 276 (58.0) | |
| **Ethnicity** | | |
| *Dutch* | 238 (50.0) | |
| *South-Asian Surinamese* | 238 (50.0) | |
| **Follow-up time, years** | 6.46 ± 1.29 | |
| **BMI, kg/m$^2$** | 25.66 ± 3.94 | 26.03 ± 4.07 |
| **Smoking status** | | |
| *Current smoker* | 79 (16.7) | 65 (13.7) |
| *Never smoked* | 239 (50.4) | 213 (44.7) |
| *Former smoker* | 156 (32.9) | 198 (41.6) |
| **Alcohol consumption** | | |
| *Low* | 293 (61.8) | 251 (72.3) |
| *Moderate* | 121 (25.5) | 73 (21.0) |
| *High* | 60 (12.7) | 23 (6.6) |
| **Diabetes mellitus** | 25 (5.3) | 53 (11.2) |
| **Systolic BP, mmHg** | 127.36 ± 17.18 | 126.55 ± 17.98 |
| **Diastolic BP, mmHg** | 79.38 ± 9.88 | 77.69 ± 10.13 |
| **Hypertension** | 169 (35.5) | 194 (40.8) |
| **Metabolic syndrome** | 176 (37.1) | 205 (43.2) |
| **PPI use** | 44 (9.2) | 74 (15.5) |
| **Metformin use** | 18 (3.8) | 36 (7.6) |
| **Statin use** | 64 (13.4) | |
| **Total cholesterol, mmol/L** | 5.21 ± 1.00 | 5.32 ± 1.08 |
| **LDL cholesterol, mmol/L** | 3.21 ± 0.90 | 3.19 ± 0.94 |
| **Triglycerides, mmol/L** | 0.91 [0.64–1.31] | 1.00 [0.69–1.41] |
| **HbA1c, mmol/mol** | 38.91 ± 6.05 | 39.46 ± 7.39 |

*Continuous variables are presented as mean ± SD or median [IQR]; categorical variables are presented as n (%). Some variables were not available at follow-up, or redundant because they were static, these fields are empty. BP, blood pressure; PPI, proton pump inhibitor. 476 subjects are presented in this table since samples were matched using these subjects, while 475 subjects had paired shotgun data that could be included in the paired analyses.*

### Supplementary Table 4: HELIUS population with shotgun metagenomics data – stratified for ethnicity

| Characteristic | Baseline | | | Follow-up | | |
|---|---|---|---|---|---|---|
| | Dutch | South-Asian Surinamese | p | Dutch | South-Asian Surinamese | p |
| **n** | **238** | **238** | | **238** | **238** | |
| **Age, years** | 51.71 ± 11.24 | 51.12 ± 9.82 | 0.546 | 58.13 ± 11.47 | 57.66 ± 9.94 | 0.636 |
| **Female sex** | 138 (58.0) | 138 (58.0) | 1 | - | - | - |
| **Follow-up time, years** | - | - | - | 6.41 ± 1.35 | 6.50 ± 1.22 | 0.451 |
| **Residence duration, years** | - | 33.14 ± 7.84 | - | - | - | - |
| **BMI, kg/m²** | 25.46 ± 3.79 | 25.86 ± 4.10 | 0.271 | 25.79 ± 4.07 | 26.27 ± 4.07 | 0.201 |
| **Current non-smoker** | 203 (85.7) | 192 (81.0) | 0.218 | 212 (89.1) | 199 (83.6) | 0.109 |
| **Alcohol consumption** | | | <0.001 | | | <0.001 |
| *Low* | 87 (36.7) | 206 (86.9) | | 73 (47.7) | 178 (91.8) | |
| *Moderate* | 101 (42.6) | 20 (8.4) | | 58 (37.9) | 15 (7.7) | |
| *High* | 49 (20.7) | 11 (4.6) | | 22 (14.4) | 1 (0.5) | |
| **Diabetes mellitus** | 11 (4.6) | 14 (5.9) | 0.673 | 14 (5.9) | 39 (16.7) | <0.001 |
| **Hypertension** | 86 (36.1) | 83 (34.9) | 0.848 | 94 (39.5) | 100 (42.0) | 0.641 |
| **Metabolic syndrome** | 74 (31.1) | 102 (43.2) | 0.008 | 83 (34.9) | 122 (51.7) | <0.001 |
| **Dyslipidaemia** | 151 (63.4) | 162 (68.1) | 0.334 | 166 (69.7) | 185 (77.7) | 0.061 |
| **Systolic BP, mmHg** | 126.04 ± 17.42 | 128.68 ± 16.88 | 0.094 | 125.89 ± 18.17 | 127.21 ± 17.79 | 0.423 |
| **Diastolic BP, mmHg** | 77.96 ± 9.57 | 80.80 ± 10.00 | 0.002 | 77.16 ± 9.75 | 78.22 ± 10.49 | 0.251 |
| **PPI use** | 22 (9.2) | 22 (9.2) | 1 | 29 (12.2) | 45 (18.9) | 0.058 |
| **Metformin use** | 9 (3.8) | 9 (3.8) | 1 | 7 (2.9) | 29 (12.2) | <0.001 |
| **Statin use** | 32 (13.4) | 32 (13.4) | 1 | - | - | - |
| **Antihypertensive use** | 42 (17.6) | 42 (17.6) | 1 | 58 (24.4) | 71 (29.8) | 0.216 |
| **Glucose-lowering drugs** | 9 (3.8) | 9 (3.8) | 1 | 8 (3.4) | 32 (13.4) | <0.001 |
| **Total cholesterol, mmol/L** | 5.26 ± 1.04 | 5.17 ± 0.95 | 0.338 | 5.47 ± 1.06 | 5.16 ± 1.08 | 0.002 |
| **LDL cholesterol, mmol/L** | 3.17 ± 0.93 | 3.26 ± 0.87 | 0.286 | 3.25 ± 0.94 | 3.12 ± 0.94 | 0.165 |
| **Triglycerides, mmol/L** | 0.82 [0.55–1.19] | 0.98 [0.71–1.42] | <0.001 | 0.95 [0.62–1.41] | 1.02 [0.75–1.45] | 0.135 |
| **HbA1c, mmol/mol** | 36.78 ± 4.08 | 41.07 ± 6.91 | <0.001 | 37.02 ± 5.04 | 41.95 ± 8.50 | <0.001 |

*Continuous variables are presented as mean ± SD or median [IQR]; categorical variables are presented as n (%). Some variables were not available at follow-up, or redundant because they were static, these fields are empty. BP, blood pressure; PPI, proton pump inhibitor. 476 subjects are presented in this table since samples were matched using these subjects, while 475 subjects had paired shotgun data that could be included in the paired analyses.*

### Supplementary Table 5: STROBE checklist

| | Item No | Recommendation | Page number |
|---|---|---|---|
| **Title and abstract** | 1 | (*a*) Indicate the study's design with a commonly used term in the title or the abstract | 1 |
| | | (*b*) Provide in the abstract an informative and balanced summary of what was done and what was found | 2 |
| **Introduction** | | | |
| Background/rationale | 2 | Explain the scientific background and rationale for the investigation being reported | 4-5 |
| Objectives | 3 | State specific objectives, including any prespecified hypotheses | 5 |
| **Methods** | | | |
| Study design | 4 | Present key elements of study design early in the paper | 6 |
| Setting | 5 | Describe the setting, locations, and relevant dates, including periods of recruitment, exposure, follow-up, and data collection | 6-7 |
| Participants | 6 | (*a*) Give the eligibility criteria, and the sources and methods of selection of participants. Describe methods of follow-up | 6-7 |
| | | (*b*) For matched studies, give matching criteria and number of exposed and unexposed | 7-8 |
| Variables | 7 | Clearly define all outcomes, exposures, predictors, potential confounders, and effect modifiers. Give diagnostic criteria, if applicable | 21-23 |
| Data sources/ measurement | 8* | For each variable of interest, give sources of data and details of methods of assessment (measurement). Describe comparability of assessment methods if there is more than one group | 7-12 |
| Bias | 9 | Describe any efforts to address potential sources of bias | 6 |
| Study size | 10 | Explain how the study size was arrived at | 6, Fig S1 |
| Quantitative variables | 11 | Explain how quantitative variables were handled in the analyses. If applicable, describe which groupings were chosen and why | 6-12 |
| Statistical methods | 12 | (*a*) Describe all statistical methods, including those used to control for confounding | 12-17 |
| | | (*b*) Describe any methods used to examine subgroups and interactions | 12-17 |
| | | (*c*) Explain how missing data were addressed | 12-17 |
| | | (*d*) If applicable, explain how loss to follow-up was addressed | N/A |
| | | (*e*) Describe any sensitivity analyses | 12-17 |
| **Results** | | | |
| Participants | 13* | (a) Report numbers of individuals at each stage of study—eg numbers potentially eligible, examined for eligibility, confirmed eligible, included in the study, completing follow-up, and analysed | 6, Fig S1 |
| | | (b) Give reasons for non-participation at each stage | Fig S1 |
| | | (c) Consider use of a flow diagram | Fig S1 |
| Descriptive data | 14* | (a) Give characteristics of study participants (eg demographic, clinical, social) and information on exposures and potential confounders | Table S1-S4 |

| | | | |
|---|---|---|---|
| | | (b) Indicate number of participants with missing data for each variable of interest | Table S1-S4 |
| | | (c) Summarise follow-up time (eg, average and total amount) | 18, Fig 1, Fig S9 |
| Outcome data | 15* | Report numbers of outcome events or summary measures over time | N/A |
| Main results | 16 | (*a*) Give unadjusted estimates and, if applicable, confounder-adjusted estimates and their precision (eg, 95% confidence interval). Make clear which confounders were adjusted for and why they were included | Throughout ms |
| | | (*b*) Report category boundaries when continuous variables were categorized | N/A |
| | | (*c*) If relevant, consider translating estimates of relative risk into absolute risk for a meaningful time period | N/A |
| Other analyses | 17 | Report other analyses done—eg analyses of subgroups and interactions, and sensitivity analyses | 18-27 |
| **Discussion** | | | |
| Key results | 18 | Summarise key results with reference to study objectives | 28 |
| Limitations | 19 | Discuss limitations of the study, taking into account sources of potential bias or imprecision. Discuss both direction and magnitude of any potential bias | 28 |
| Interpretation | 20 | Give a cautious overall interpretation of results considering objectives, limitations, multiplicity of analyses, results from similar studies, and other relevant evidence | 28-30 |
| Generalisability | 21 | Discuss the generalisability (external validity) of the study results | 30 |
| **Other information** | | | |
| Funding | 22 | Give the source of funding and the role of the funders for the present study and, if applicable, for the original study on which the present article is based | 31 |

#### Supplementary Figure 1: Flowchart participant selection

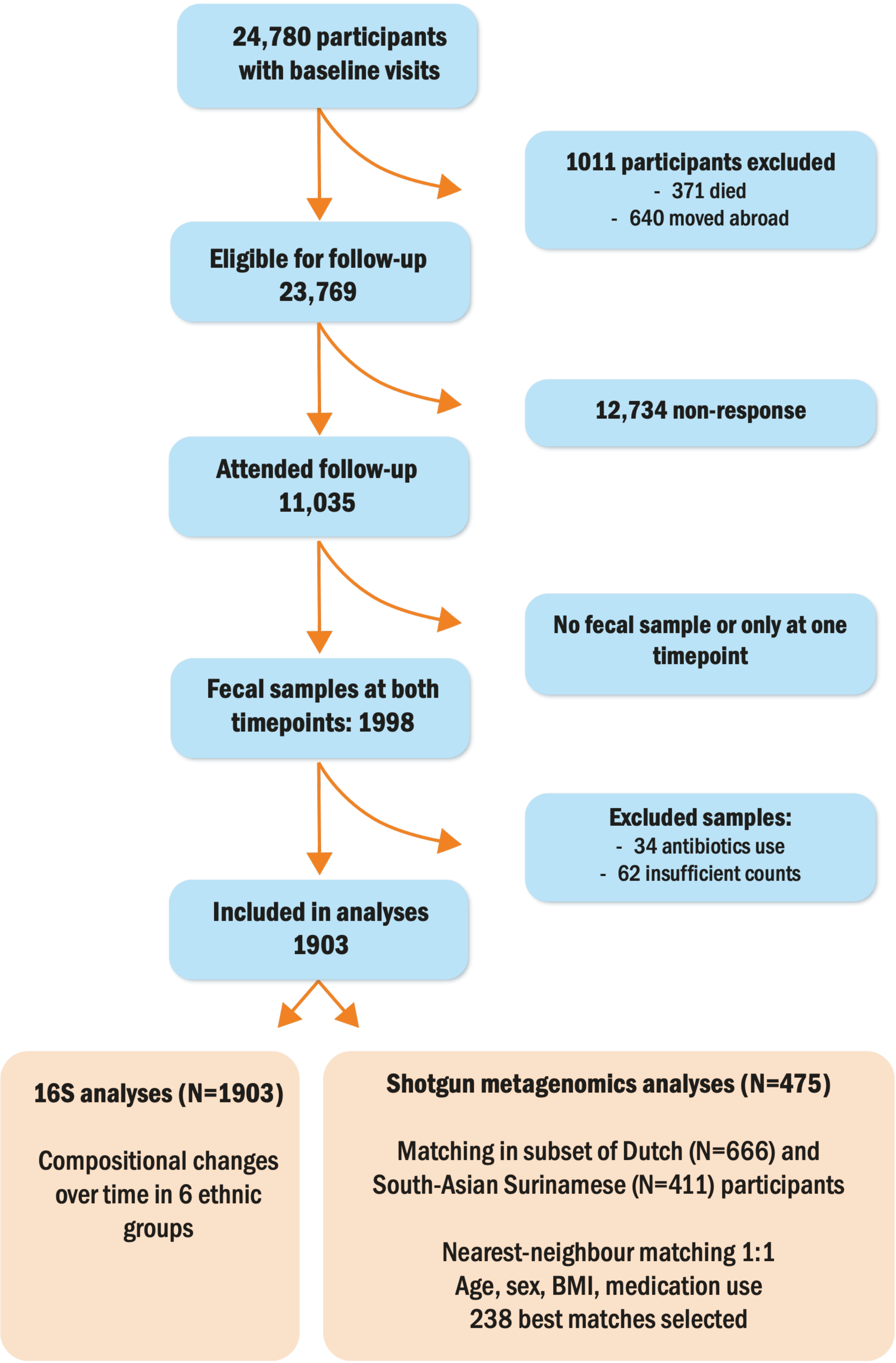

### Supplementary Figure 2: Ethnic differences in Bray-Curtis over time

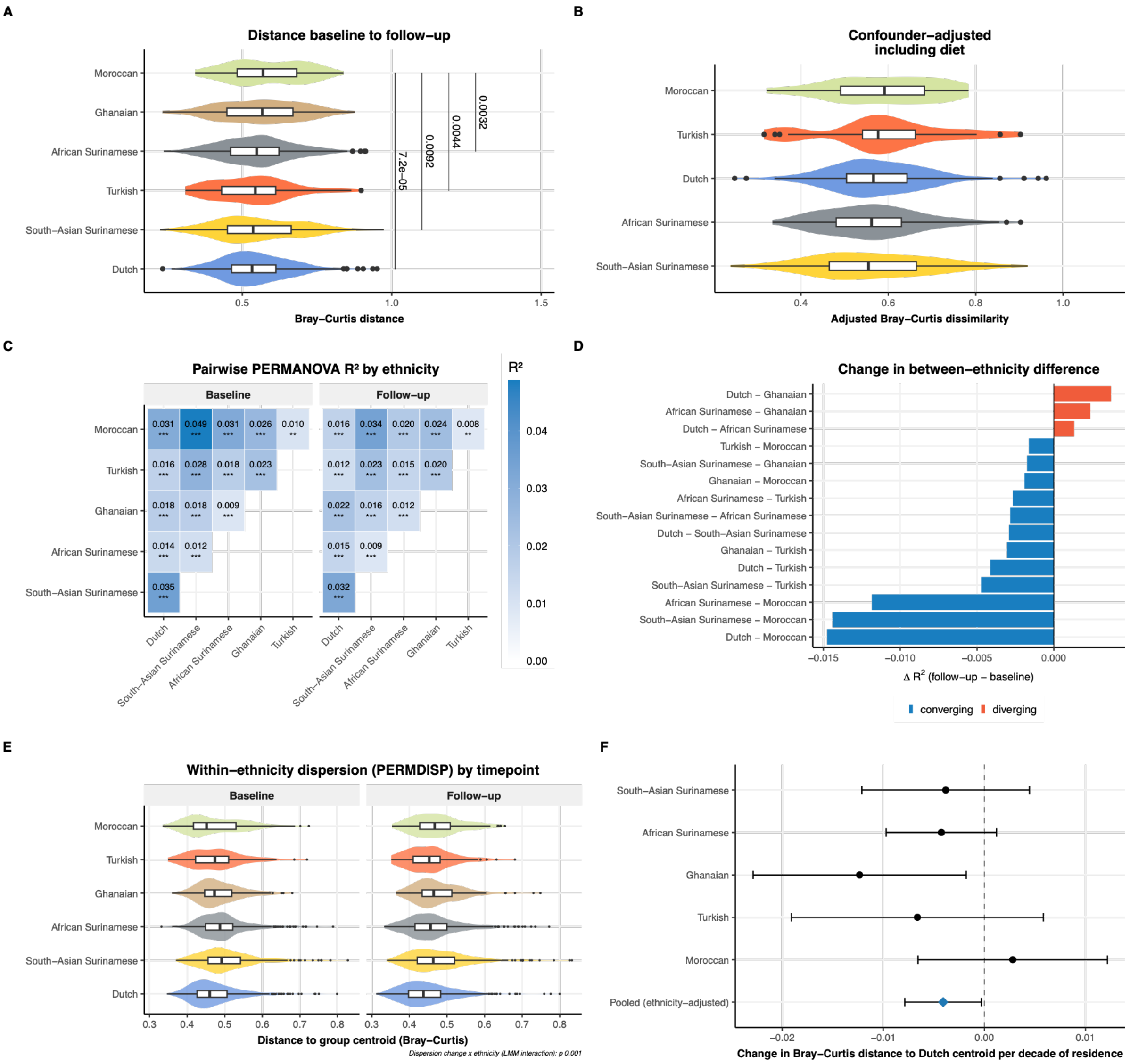


**Supplementary Figure 2: Between-ethnicity differences in gut microbiota compositional change over follow-up.** (A) Unadjusted Bray-Curtis dissimilarity between baseline and follow-up samples per participant, stratified by ethnicity. Distributions are shown as violin plots with embedded boxplots. Ethnic groups are ordered by median dissimilarity. P-values from pairwise Wilcoxon rank-sum tests are shown for selected comparisons. (B) Confounder- and diet-adjusted Bray-Curtis dissimilarity per ethnicity (N = 799). Residuals from a linear regression model including age, sex, BMI, diet principal components 1 and 2 (Supplementary Figure 4), metformin use, proton pump inhibitor use, and follow-up time were added to the grand mean to obtain adjusted dissimilarity values. Ethnic groups are ordered by median adjusted dissimilarity. (C) Pairwise PERMANOVA effect sizes at baseline and follow-up between all ethnic group pairs. (D) Change in pairwise PERMANOVA effect size ($\Delta R^2$) between baseline and follow-up for each pair of ethnic groups. $\Delta R^2$ was calculated as $R^2$(follow-up) − $R^2$(baseline). (E) Within-ethnicity dispersion in gut microbiota composition

(PERMDISP) at baseline and follow-up. The LMM ethnicity × timepoint interaction was significant (p-value 0.001). Differences between groups were tested with Wilcoxon rank-sum tests. (F) Estimates and 95% confidence intervals of the associations between residence duration and gut microbiota similarity to the Dutch reference population, tested with unadjusted linear models. The pooled estimate is adjusted for ethnicity.

### Supplementary Figure 3: Alpha diversity differences per timepoint

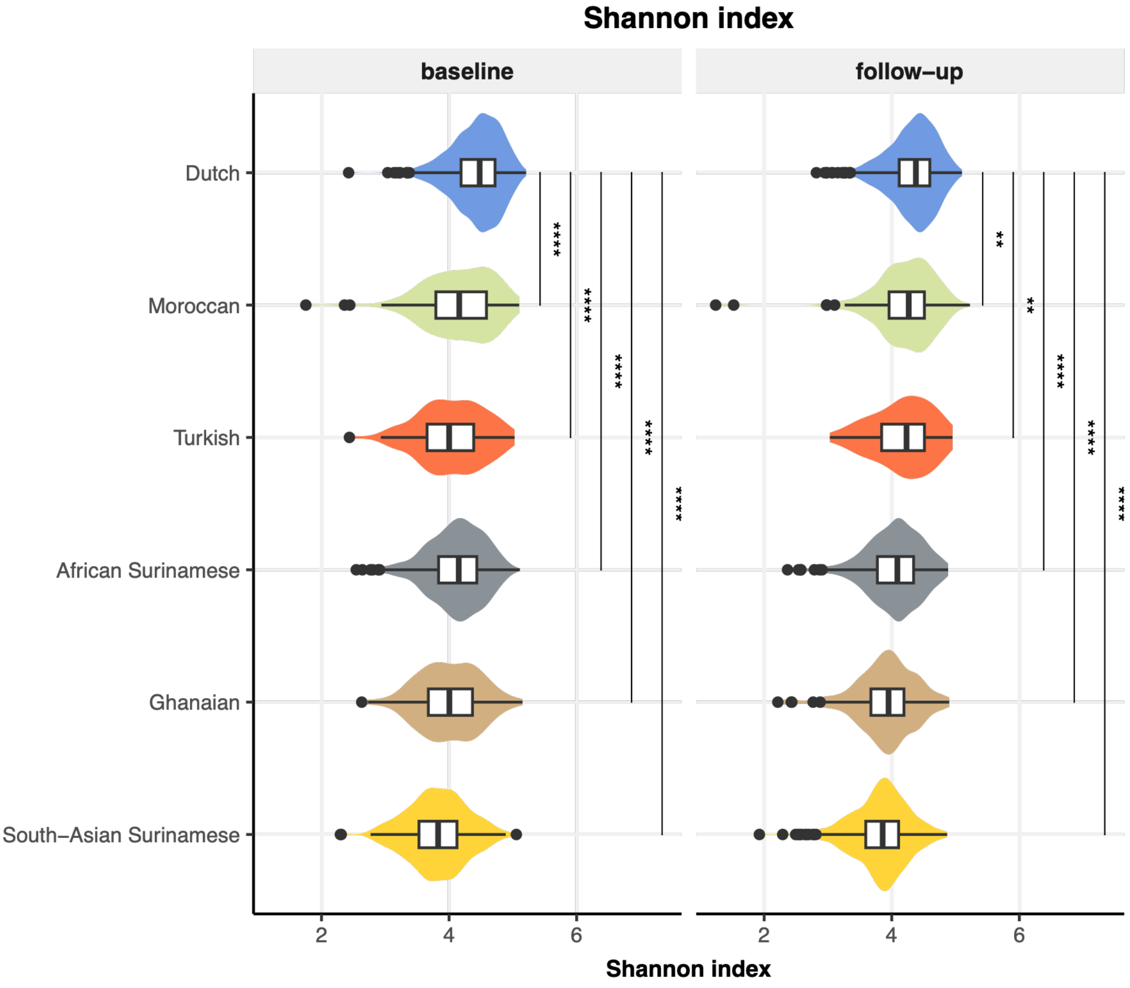


**Supplementary Figure 3: Shannon diversity by ethnic group at baseline and follow-up.** Shannon diversity index per participant, stratified by ethnic group and faceted by timepoint (baseline and follow-up). Distributions are shown as violin plots with embedded boxplots (median, IQR, 1.5× IQR). Ethnic groups are ordered by median Shannon index. Significant pairwise differences relative to Dutch participants were tested using Wilcoxon rank-sum tests; only significant comparisons are shown (ns hidden).

#### Supplementary Figure 4: PCA dietary variables

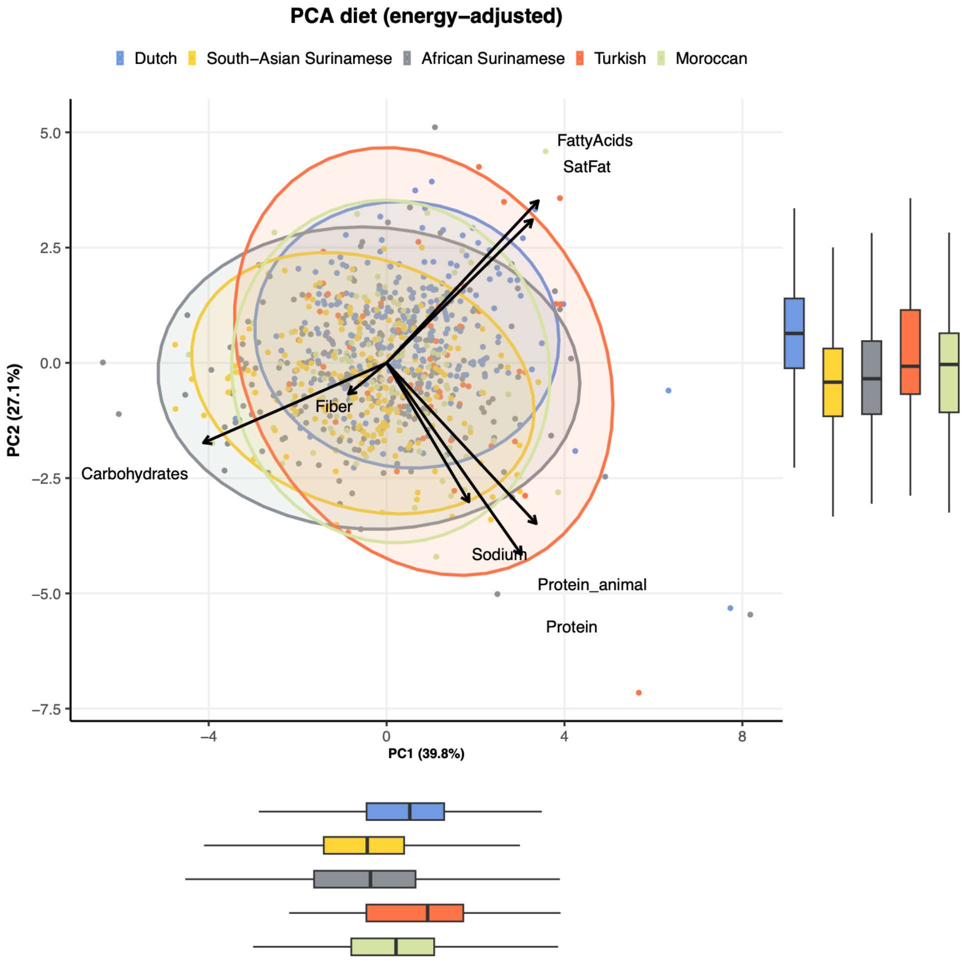


**Supplementary Figure 4. Principal component analysis of dietary macronutrient intake (energy-adjusted) by ethnicity.** Principal component analysis (PCA) was performed on nutrient variables (fibre, total protein, animal protein, fatty acids, saturated fatty acids, carbohydrates, and sodium). These variables were adjusted for total energy intake using Willett's residual method, for which nutrient groups were regressed on total energy intake using a linear regression model, followed by zero-mean-unit variance scaling. Each point represents one participant, coloured by ethnic group. Ellipses indicate 95% normal-distribution confidence regions per ethnic group. Loading vectors (arrows) show the direction and relative contribution of each macronutrient to the principal components. Boxplots show the distribution of PC1 and PC2 scores by ethnicity.

### Supplementary Figure 5: Ethnic differences in dietary intake

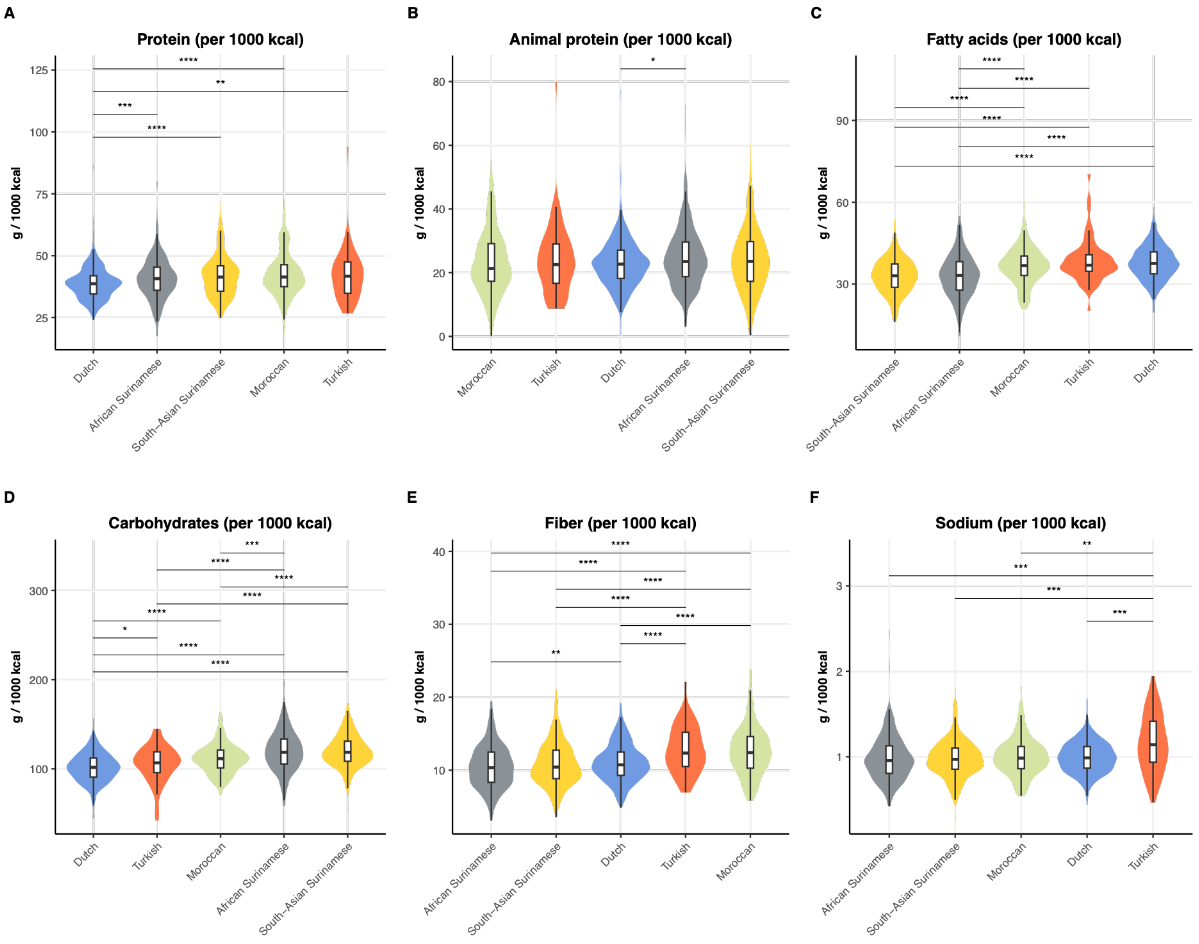


**Supplementary Figure 5: Energy-adjusted nutrient intake by ethnic group.** Distributions of energy-adjusted (per 1000 kcal total energy intake) nutrient intake stratified by ethnic group at baseline, shown as violin plots with boxplots. Quantities expressed per 1000 kcal to account for differences in total energy intake between ethnicities. Ethnic groups are ordered by median intake per nutrient. Only statistically significant pairwise comparisons are shown (Mann Whitney U tests, $p < 0.05$; ns hidden). Dietary data were not available for the Ghanaian participants. (A) Total protein (gram / 1000 kcal). (B) Animal protein (gram / 1000 kcal). (C) Fatty acids (gram / 1000 kcal). (D) Carbohydrates (gram / 1000 kcal). (E) Dietary fibre (gram / 1000 kcal). (F) Sodium (gram / 1000 kcal).

#### Supplementary Figure 6: Sensitivity analysis: baseline predictors of beta diversity

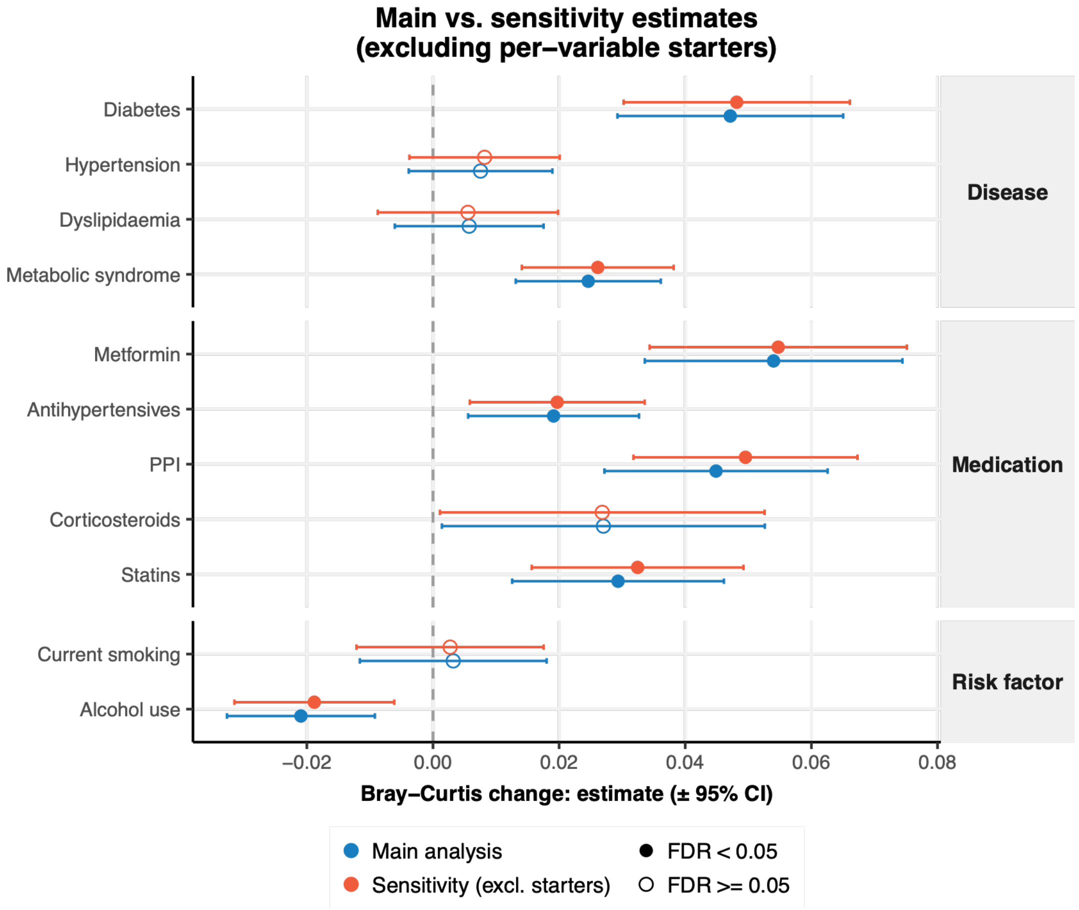


**Supplementary Figure 6: Sensitivity analysis: baseline predictors of beta diversity.** Forest plot of the association between baseline disease, medication and lifestyle status and Bray-Curtis change in gut microbiota composition between baseline and follow-up (16S). Points show linear regression estimates (Yes versus No at baseline) with 95% confidence intervals, from models adjusted for follow-up time (distance ~ predictor + follow-up time), as in Figure 2B. Blue, main analysis including all participants with available data (as in Figure 2B). Red, sensitivity analysis excluding participants who did have the exposure at follow-up for that specific variable ("starters" during follow-up). Filled points indicate FDR < 0.05 and open points FDR ≥ 0.05 (Benjamini-Hochberg, applied across the 11 variables within each analysis).

### Supplementary Figure 7: Baseline predictors of alpha diversity change

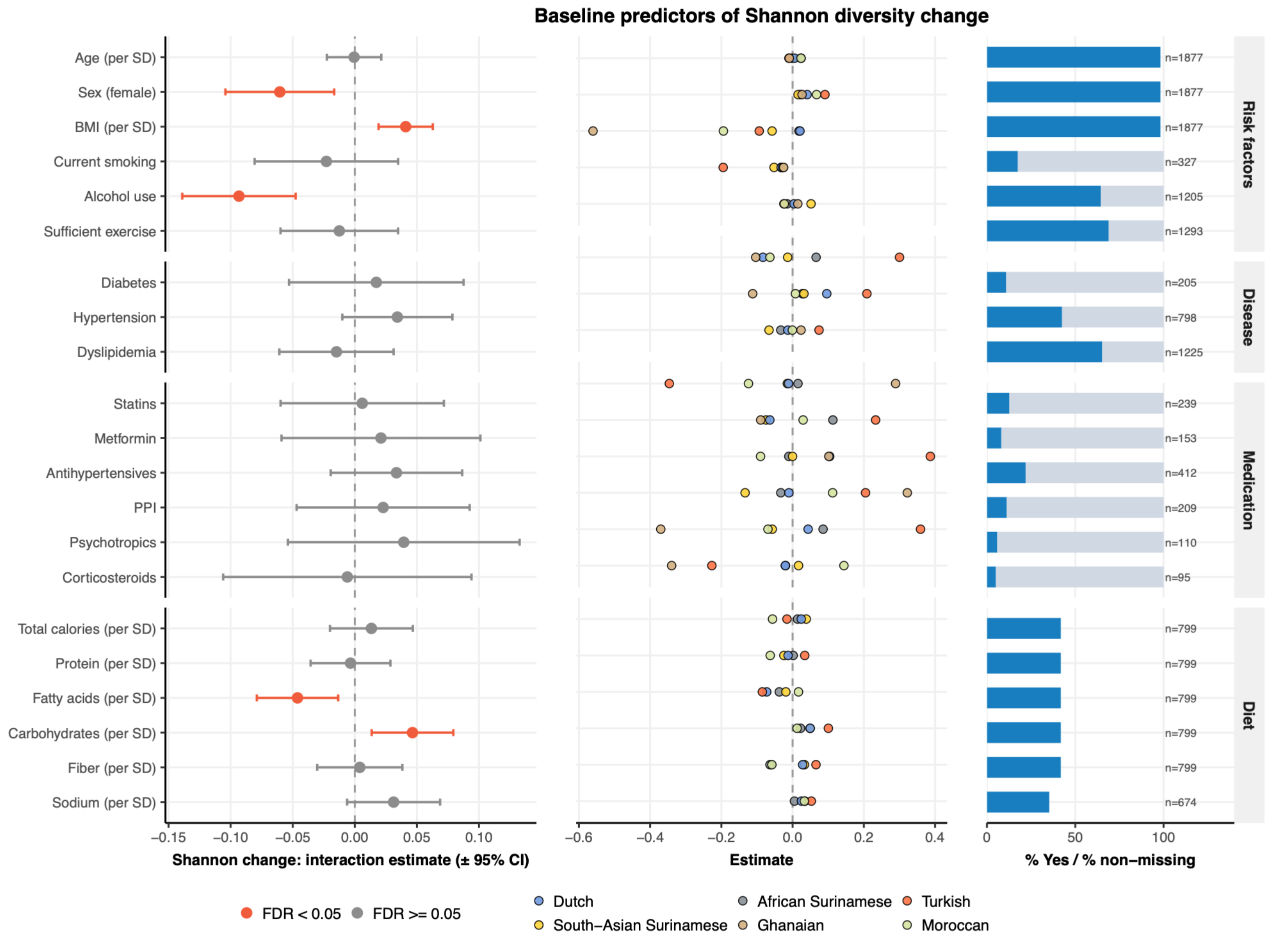


**Supplementary Figure 7: Baseline predictors of Shannon diversity change from baseline to follow-up.** For each predictor, a linear mixed model was fitted with Shannon diversity as the outcome, including predictor × timepoint interaction and follow-up time as covariate, and a random intercept for participant. The interaction estimate reflects whether the predictor is associated with a differential Shannon diversity change. Continuous predictors were standardised to unit variance (per SD). Binary predictors (disease, medication, lifestyle) were coded Yes vs. No (reference). P-values were corrected for multiple testing using the Benjamini-Hochberg procedure; estimates with FDR < 0.05 are highlighted in orange. **(Left)** Interaction estimates (±95% CI) for the full cohort, stratified by predictor domain (risk factors, cardiometabolic disease, medication, diet). **(Middle)** Ethnicity-stratified interaction estimates for the same predictors. Each point represents a separate model fitted within one ethnic group; colour indicates ethnicity. **(Right)** Prevalence of each predictor at baseline. For binary variables, the bar shows the percentage of participants with "Yes"; for continuous variables, the percentage with non-missing data. Sample sizes are annotated.

### Supplementary Figure 8: Differences in dietary intake within shotgun subset

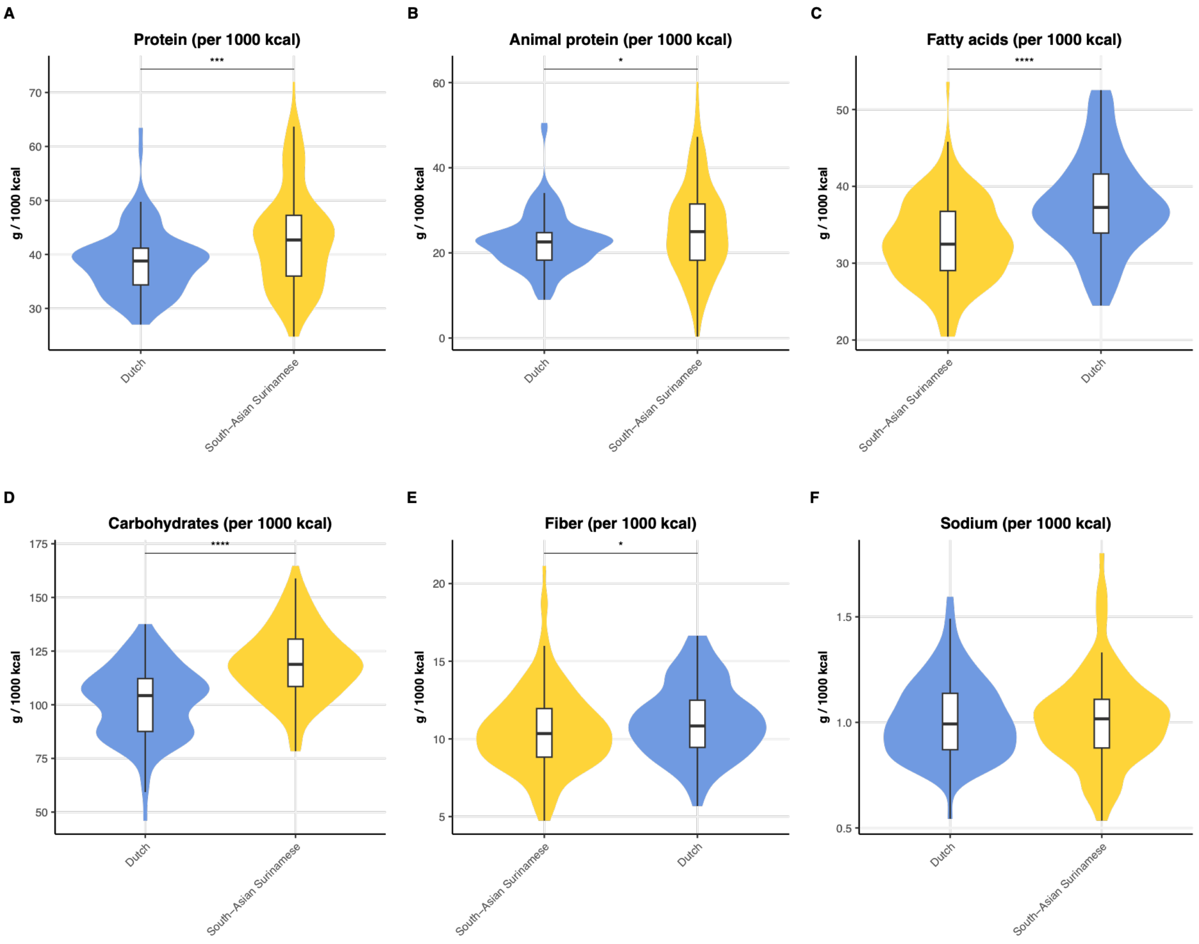


**Supplementary Figure 8: Energy-adjusted nutrient intake by ethnic group (N=233, 108 Dutch and 125 South-Asian Surinamese).** Distributions of energy-adjusted dietary nutrient intake stratified by ethnic group, shown as violin plots with embedded boxplots. Quantities expressed per 1000 kcal to account for differences in total energy intake between ethnicities. Only statistically significant pairwise comparisons are shown (Mann-Whitney U test, $p < 0.05$). (A) Total protein (gram/1000 kcal). (B) Animal protein (gram / 1000 kcal). (C) Fatty acids (gram / 1000 kcal). (D) Carbohydrates (gram / 1000 kcal). (E) Dietary fibre (gram / 1000 kcal). (F) Sodium (gram / 1000 kcal).

### Supplementary Figure 9: Alpha and beta diversity differences in shotgun data

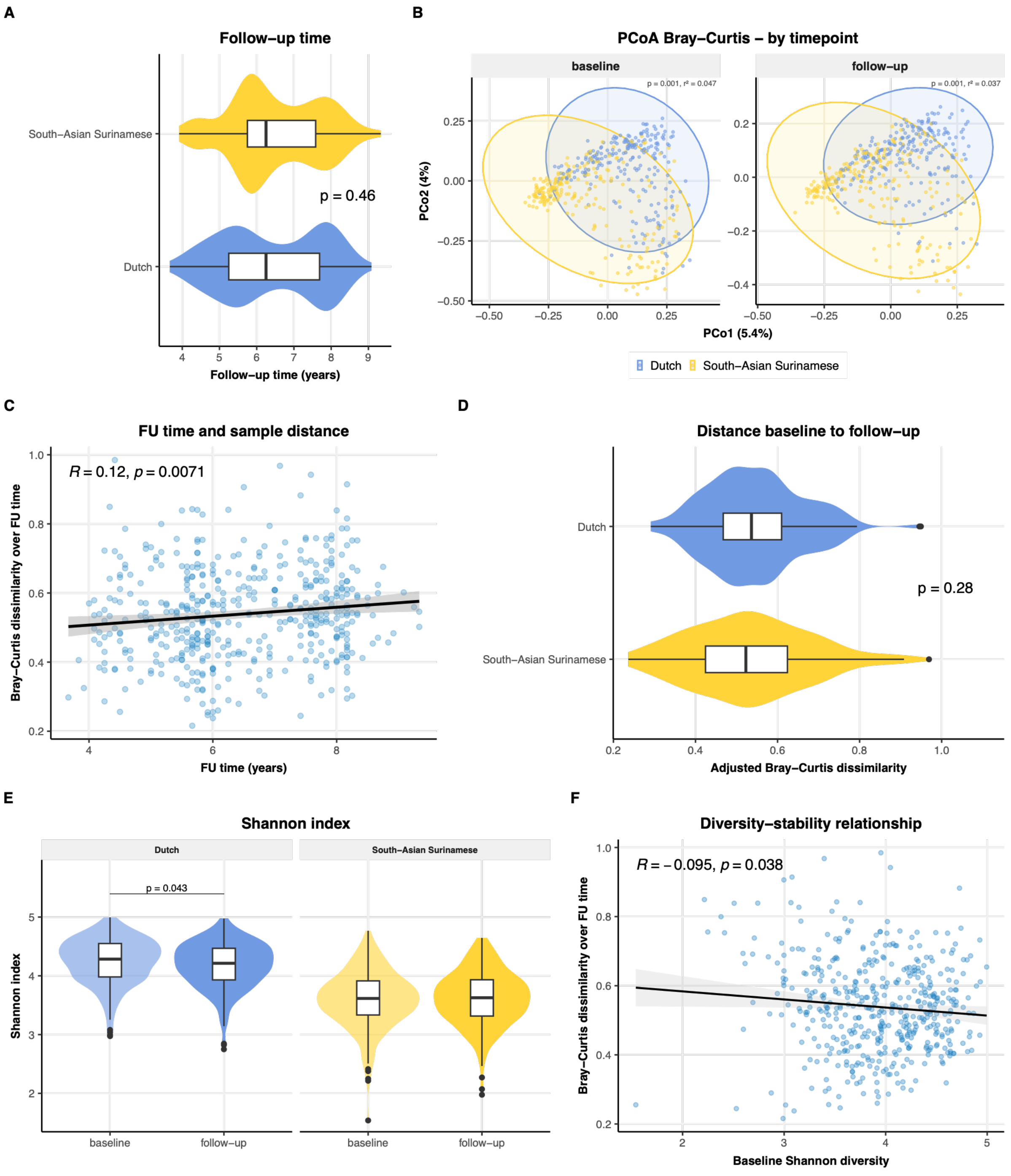


**Supplementary Figure 9. Longitudinal gut microbiome composition in the shotgun metagenomics subcohort.** (A) Follow-up time (years) between baseline and follow-up visit, stratified by ethnic group, differences tested with a Mann-Whitney U test. (B) Principal coordinates analysis (PCoA) of Bray-Curtis dissimilarity, stratified by ethnic group. Each point represents one sample; ellipses denote the 95% confidence region for each timepoint. PERMANOVA $R^2$ and p-value are shown per panel. (C) Association between follow-up time (years) and intra-individual Bray-Curtis dissimilarity between paired samples. Line shows linear regression with 95% confidence interval; Pearson correlation coefficient and p-value are annotated. (D) Intra-individual Bray-Curtis dissimilarity between paired baseline and follow-up samples, stratified by ethnic group, after adjustment for age, sex, BMI, Metformin use, PPI use, and follow-up time. Residuals were mean-centred. P-value from Wilcoxon rank-sum test. (E) Shannon diversity index at baseline and follow-up, stratified by ethnic group. P-values from Wilcoxon signed-rank test; only significant comparisons ($p < 0.05$) are shown. (F) Association between baseline Shannon diversity and intra-individual Bray-Curtis dissimilarity over the follow-up period. Line shows linear regression with 95% confidence interval; Pearson correlation coefficient and p-value are annotated.

### Supplementary Figure 10: Cross-sectional differences

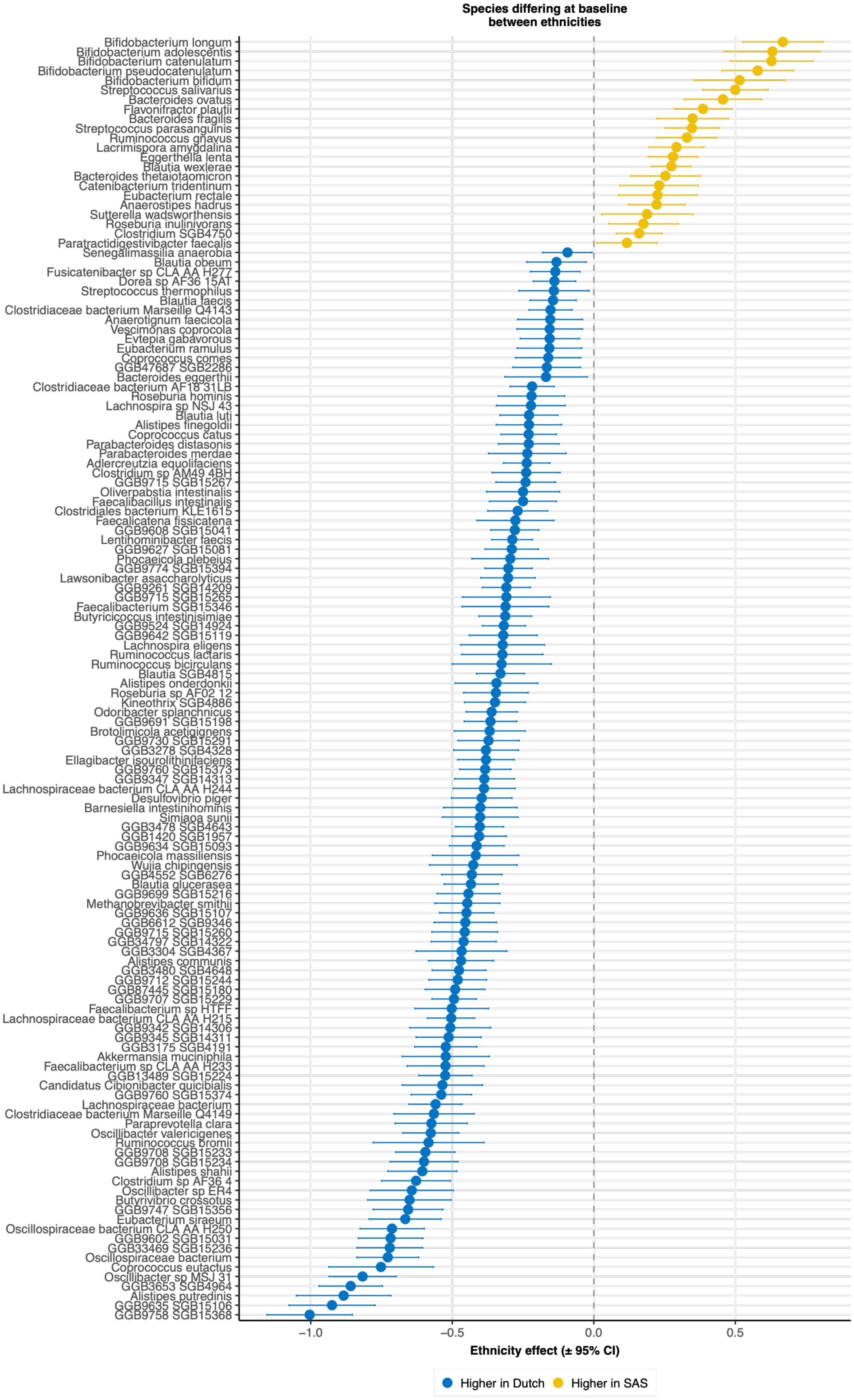

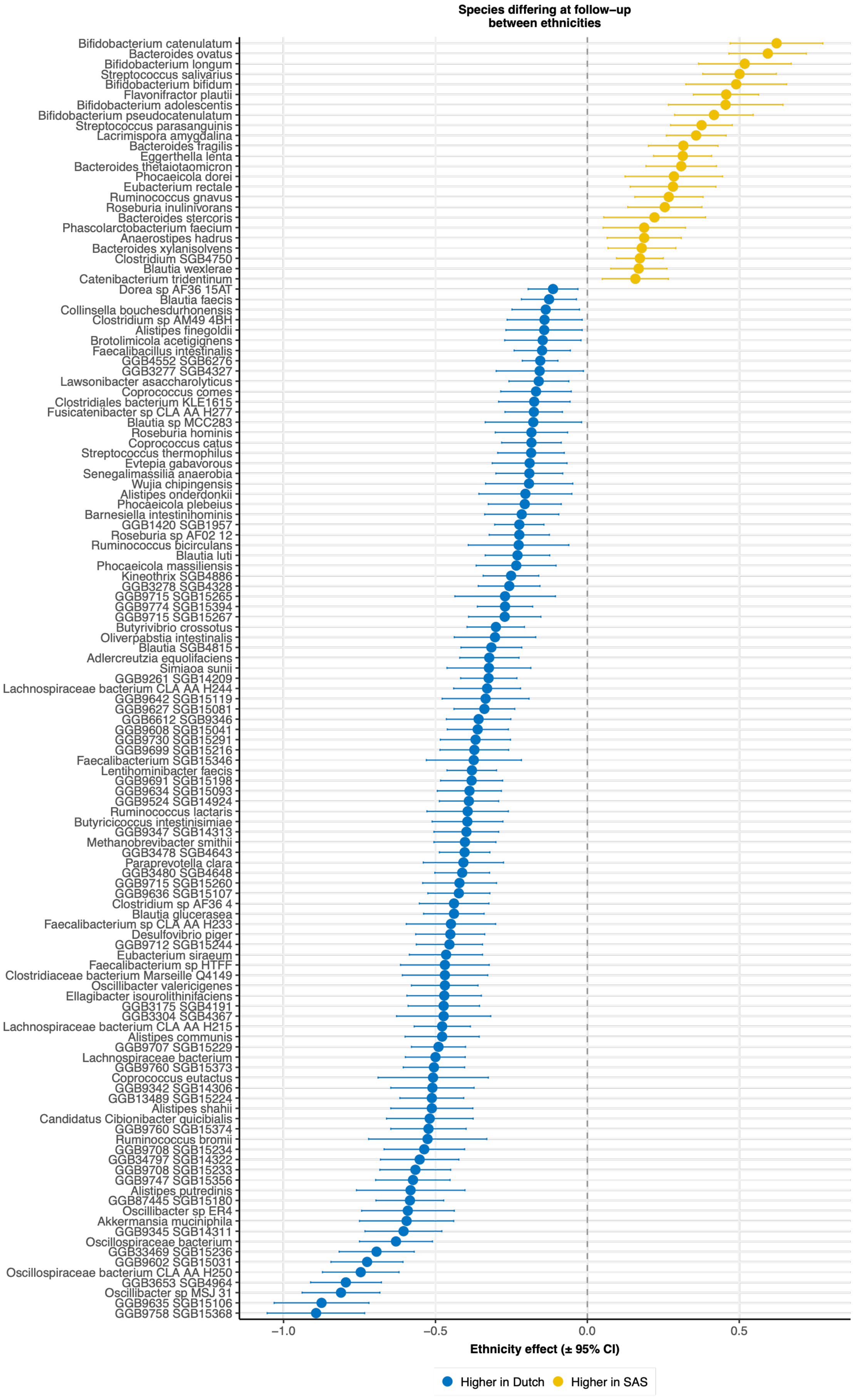
Species differing at follow−up
between ethnicities
Bifidobacterium catenulatum
Bacteroides ovatus
Bifidobacterium longum
Streptococcus salivarius
Bifidobacterium bifidum
Flavonifractor plautii
Bifidobacterium adolescentis
Bifidobacterium pseudocatenulatum
Streptococcus parasanguinis
Lacrimispora amygdalina
Bacteroides fragilis
Eggerthella lenta
Bacteroides thetaiotaomicron
Phocaeicola dorei
Eubacterium rectale
Ruminococcus gnavus
Roseburia inulinivorans
Bacteroides stercoris
Phascolarctobacterium faecium
Anaerostipes hadrus
Bacteroides xylanisolvens
Clostridium SGB4750
Blautia wexlerae
Catenibacterium tridentinum
Dorea sp AF36 15AT
Blautia faecis
Collinsella bouchesdurhonensis
Clostridium sp AM49 4BH
Alistipes finegoldii
Brotolimicola acetigignens
Faecalibacillus intestinalis
GGB4552 SGB6276
GGB3277 SGB4327
Lawsonibacter asaccharolyticus
Coprococcus comes
Clostridiales bacterium KLE1615
Fusicatenibacter sp CLA AA H277
Blautia sp MCC283
Roseburia hominis
Coprococcus catus
Streptococcus thermophilus
Evtepia gabavorous
Senegalimassilia anaerobia
Wujia chipingensis
Alistipes onderdonkii
Phocaeicola plebeius
Barnesiella intestinihominis
GGB1420 SGB1957
Roseburia sp AF02 12
Ruminococcus bicirculans
Blautia luti
Phocaeicola massiliensis
Kineothrix SGB4886
GGB3278 SGB4328
GGB9715 SGB15265
GGB9774 SGB15394
GGB9715 SGB15267
Butyrivibrio crossotus
Oliverpabstia intestinalis
Blautia SGB4815
Adlercreutzia equolifaciens
Simiaoa sunii
GGB9261 SGB14209
Lachnospiraceae bacterium CLA AA H244
GGB9642 SGB15119
GGB9627 SGB15081
GGB6612 SGB9346
GGB9608 SGB15041
GGB9730 SGB15291
GGB9699 SGB15216
Faecalibacterium SGB15346
Lentihominibacter faecis
GGB9691 SGB15198
GGB9634 SGB15093
GGB9524 SGB14924
Ruminococcus lactaris
Butyricicoccus intestinisimiae
GGB9347 SGB14313
Methanobrevibacter smithii
GGB3478 SGB4643
Paraprevotella clara
GGB3480 SGB4648
GGB9715 SGB15260
GGB9636 SGB15107
Clostridium sp AF36 4
Blautia glucerasea
Faecalibacterium sp CLA AA H233
Desulfovibrio piger
GGB9712 SGB15244
Eubacterium siraeum
Faecalibacterium sp HTFF
Clostridiaceae bacterium Marseille Q4149
Oscillibacter valericigenes
Ellagibacter isourolithinifaciens
GGB3175 SGB4191
GGB3304 SGB4367
Lachnospiraceae bacterium CLA AA H215
Alistipes communis
GGB9707 SGB15229
Lachnospiraceae bacterium
GGB9760 SGB15373
Coprococcus eutactus
GGB9342 SGB14306
GGB13489 SGB15224
Alistipes shahii
Candidatus Cibionibacter quicibialis
GGB9760 SGB15374
Ruminococcus bromii
GGB9708 SGB15234
GGB34797 SGB14322
GGB9708 SGB15233
GGB9747 SGB15356
Alistipes putredinis
GGB87445 SGB15180
Oscillibacter sp ER4
Akkermansia muciniphila
GGB9345 SGB14311
Oscillospiraceae bacterium
GGB33469 SGB15236
GGB9602 SGB15031
Oscillospiraceae bacterium CLA AA H250
GGB3653 SGB4964
Oscillibacter sp MSJ 31
GGB9635 SGB15106
GGB9758 SGB15368
−1.0
−0.5
0.0
0.5
Ethnicity effect (± 95% CI)
Higher in Dutch
Higher in SAS

**Supplementary Figure 10. Gut microbiota species differing between ethnicities at baseline and follow-up.** Forest plots showing gut microbiota species with significantly different abundance between Dutch and South Asian Surinamese (SAS) participants at baseline (A) and follow-up (B). Species were included if present at >0.1% relative abundance in >20% of participants at either timepoint. Abundances were log10-transformed prior to analysis. Linear regression models were fitted for each species with ethnicity as the primary predictor. Only species with a false discovery rate (FDR)-corrected q-value <0.05 are shown. Points represent the regression coefficient for ethnicity (SAS vs. Dutch); horizontal lines indicate 95% confidence intervals. Colour indicates the direction of the effect (blue: higher in Dutch; orange: higher in SAS). Species are ordered by effect size.

### Supplementary Figure 11: Differential change in species over time

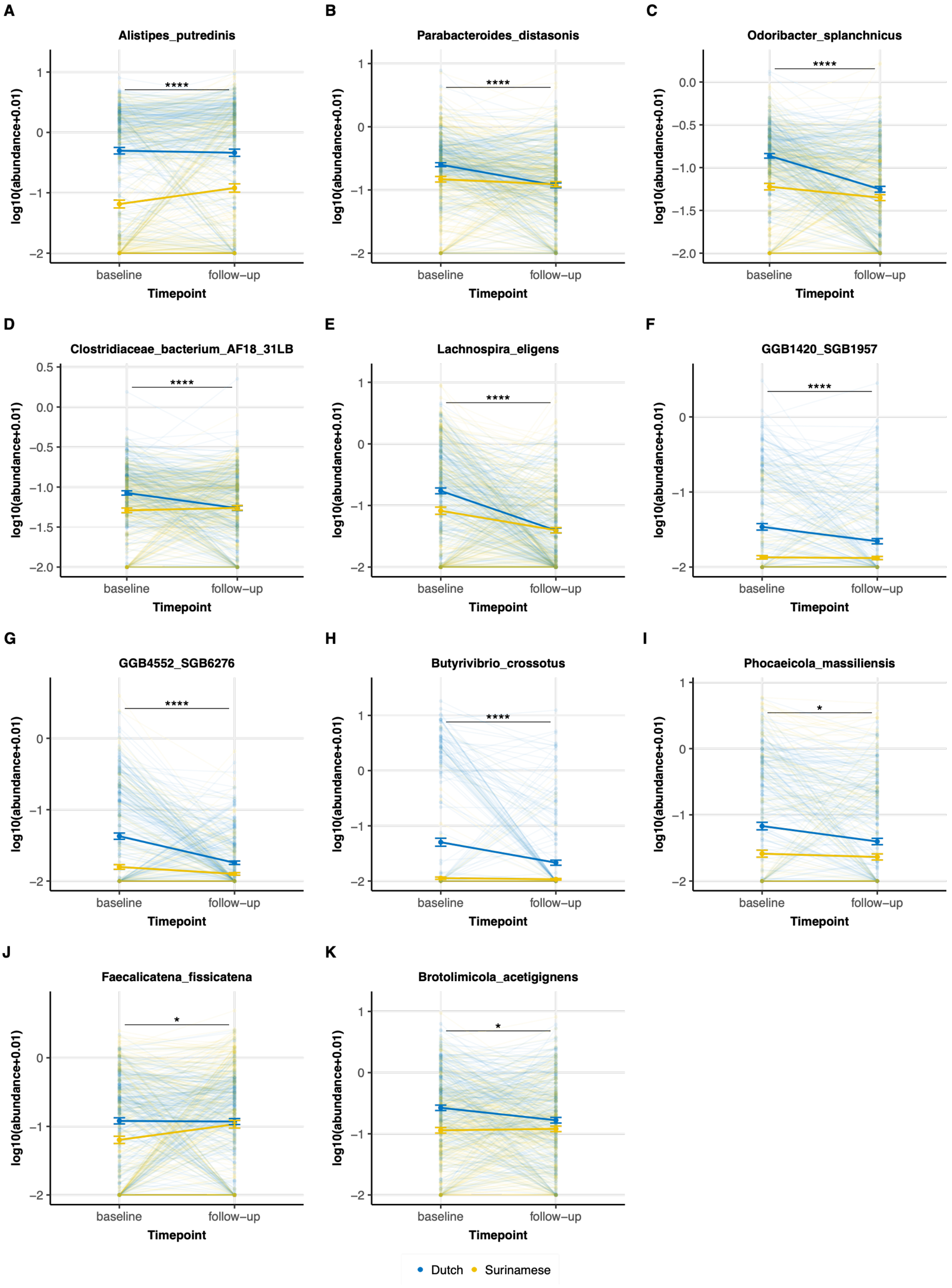

**Supplementary Figure 11. Gut microbiota species with a significant ethnicity × time interaction.** Abundance trajectories from baseline to follow-up for gut microbiota species showing a significant ethnicity × timepoint interaction. Species were included if present at >0.1% relative abundance in >20% of participants in either the Dutch or South-Asian Surinamese group. Abundances were log10-transformed. Linear mixed models (LMM) were fitted for each species with ethnicity, timepoint, and their interaction as fixed effects, with participant included as a random intercept. Panels show each species passing an FDR-corrected q-value threshold of <0.05 for the interaction term. Thin lines represent individual participant trajectories; bold lines and points represent group means ± standard error of the mean. Colours indicate ethnicity. Significance brackets reflect FDR-corrected q-values (* q<0.05, ** q<0.01, *** q<0.001, **** q<0.0001). GGB4552 SGB6276 (MetaPhlAn taxonomy) was labelled as Clostridia TANB77 CAG-245 and GGB1420 SGB1957 as Bacilli UBA660 CAG-302 in GTDB.

#### Supplementary Figure 12: Strain divergence and microdiversity in *A. putredinis*

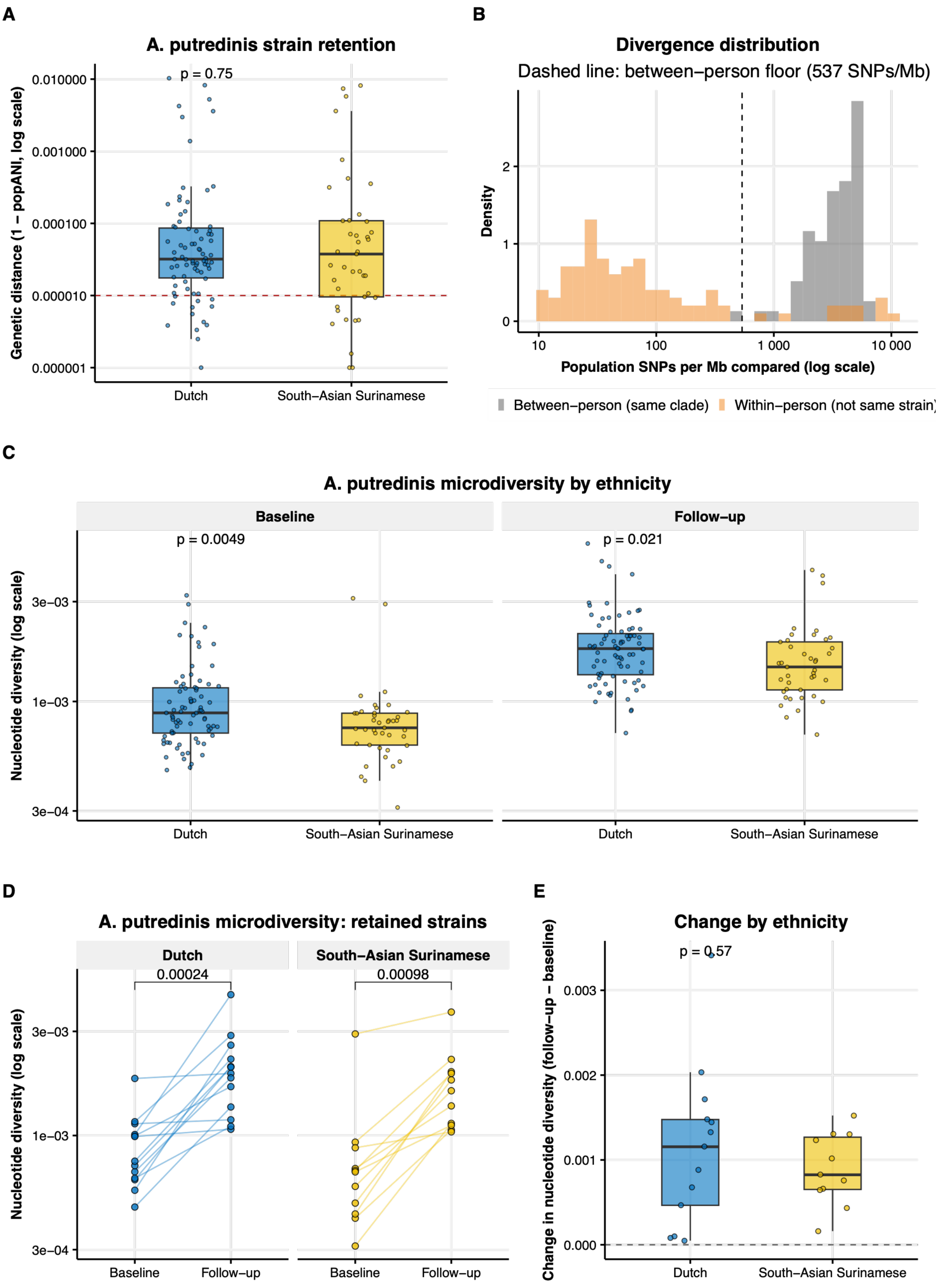

**Supplementary Figure 12: *Alistipes putredinis* within-strain microdiversity**. (A) Genetic distance (1 − popANI) between each participant's baseline and follow-up *A. putredinis* population, by ethnicity (n = 120: 79 Dutch, 41 South-Asian Surinamese). The red dashed line marks the conventional same-strain threshold (popANI ≥ 0.99999), differences between ethnicities tested with a Mann-Whitney U test. (B) Distribution of population-level SNP density (SNPs per Mb compared) for within-person comparisons (n=120) versus the between-person background from unrelated individuals carrying the same *A. putredinis* clade (n = 75, grey). The dashed line is the between-person background floor (537 SNPs/Mb), used as the cutoff distinguishing divergence consistent with a persisting, related population from divergence consistent with replacement by an unrelated genome.
Nucleotide diversity (π, inStrain profile) is shown for the subset of participants whose *A. putredinis* strain was retained between timepoints (popANI≥99.999%; 13 Dutch, 11 South-Asian Surinamese). (C) Nucleotide diversity (log scale) per ethnicity, faceted by timepoint, differences tested with Mann-Whitney U tests. (D) Nucleotide diversity (log scale) at baseline and follow-up, faceted by ethnicity, differences tested with a paired two-sided Wilcoxon signed-rank tests within each ethnic group. (E) Change in nucleotide diversity (follow-up minus baseline) per participant, compared between ethnic groups; the dashed line marks no change. Difference tested with a Mann-Whitney U test.

### Supplementary Figure 13: Baseline predictors of strain stability

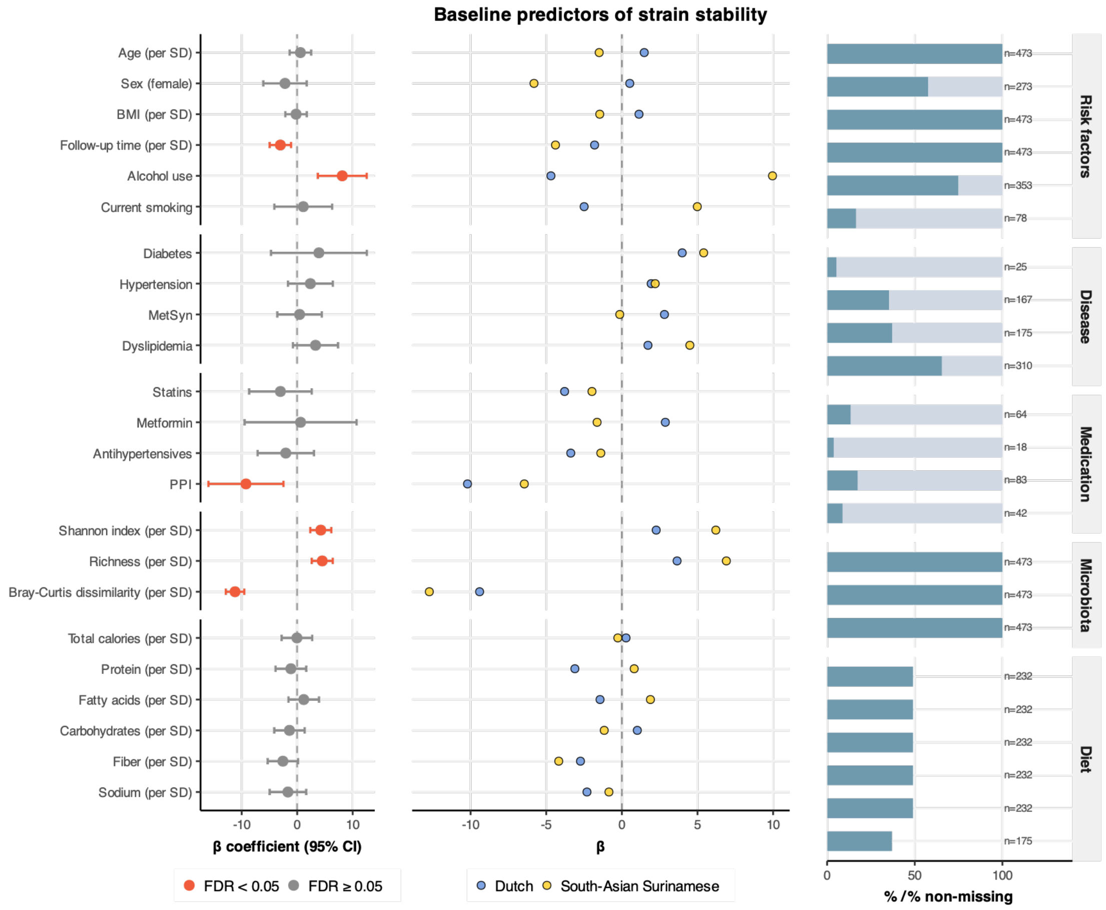


**Supplementary figure 13: Baseline predictors of longitudinal strain stability.** Forest plots showing the association between clinical covariates and overall strain retention between baseline and follow-up. Strain stability is shown in %, a proportion of the in total 211 SGBs. The outcome is the percentage of species genome bins (SGBs) for which the same strain was detected at both timepoints (strain sharing percentage). *Left panel:* β coefficients with 95% confidence intervals from linear regression models, faceted by covariate block (risk factors, disease, medication, microbiota, and diet). Continuous predictors were z-scored prior to analysis to allow comparison of effect sizes across variables. All models were adjusted for follow-up time. P-values were corrected for multiple testing using the Benjamini–Hochberg procedure; covariates with FDR < 0.05 are highlighted in orange. *Middle panel:* Stratified β coefficients estimated separately in Dutch and South-Asian Surinamese participants (unadjusted). *Right panel:* For each covariate, the proportion of participants with the exposed category (Yes / Female) or, for continuous variables, the proportion with non-missing data; absolute numbers are shown to the right of each bar.

### Supplementary Figure 14: Strain stability associations

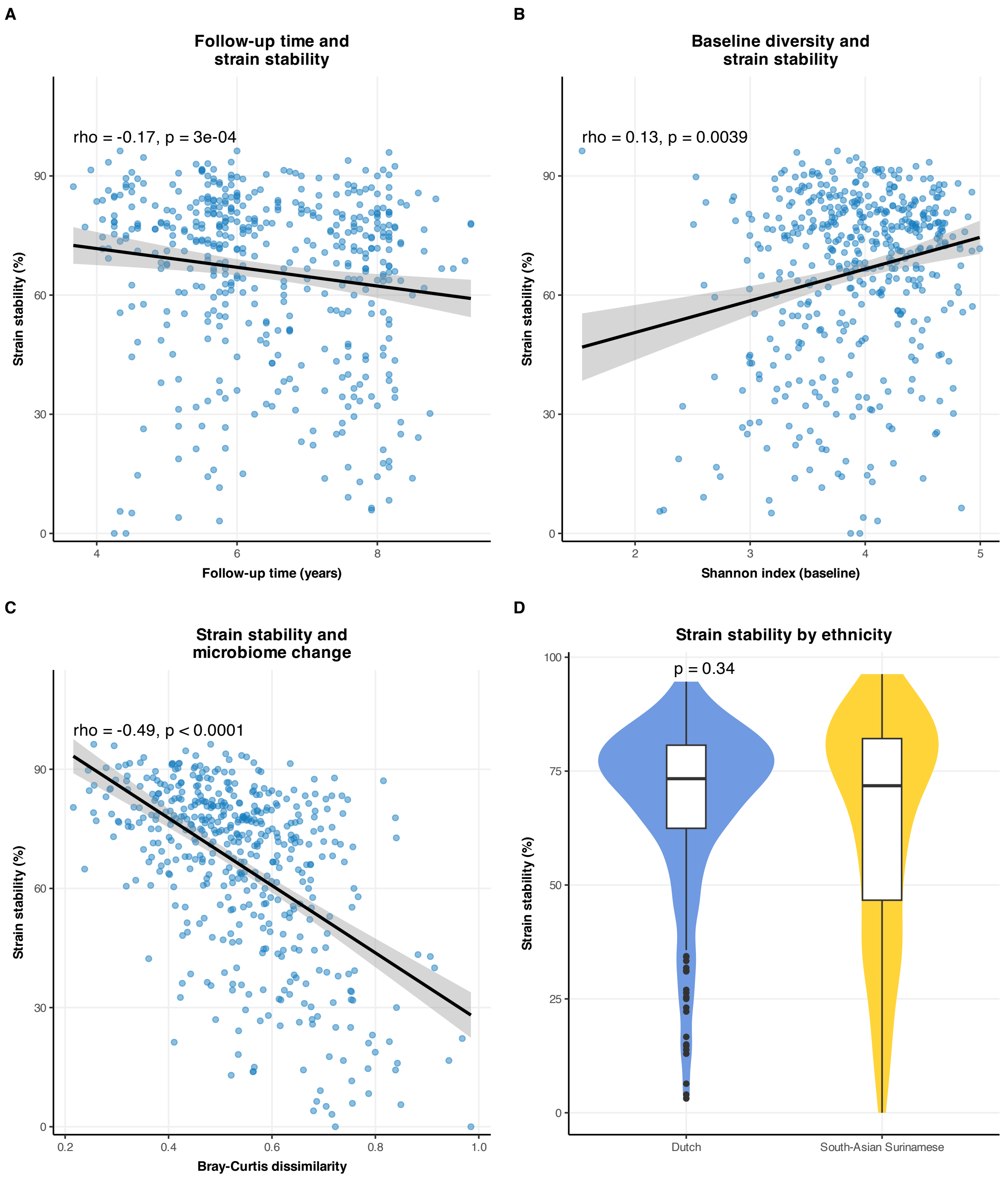


**Supplementary Figure 14: Strain stability associations.** (A-C) Strain stability (in %; proportion of 211 SGBs) in relation to follow-up time with Spearman's correlation coefficients (A), baseline Shannon diversity index (B), and Bray-Curtis dissimilarity between baseline and follow-up (C); lines show linear regression with 95% confidence intervals (β and p-value from linear regression). (D) Violin plot comparing the percentage of shared strains between baseline and follow-up timepoints in Dutch versus SAS participants.

### Supplementary Figure 15: Ethnicity and strain stability over time – logistic regression analysis

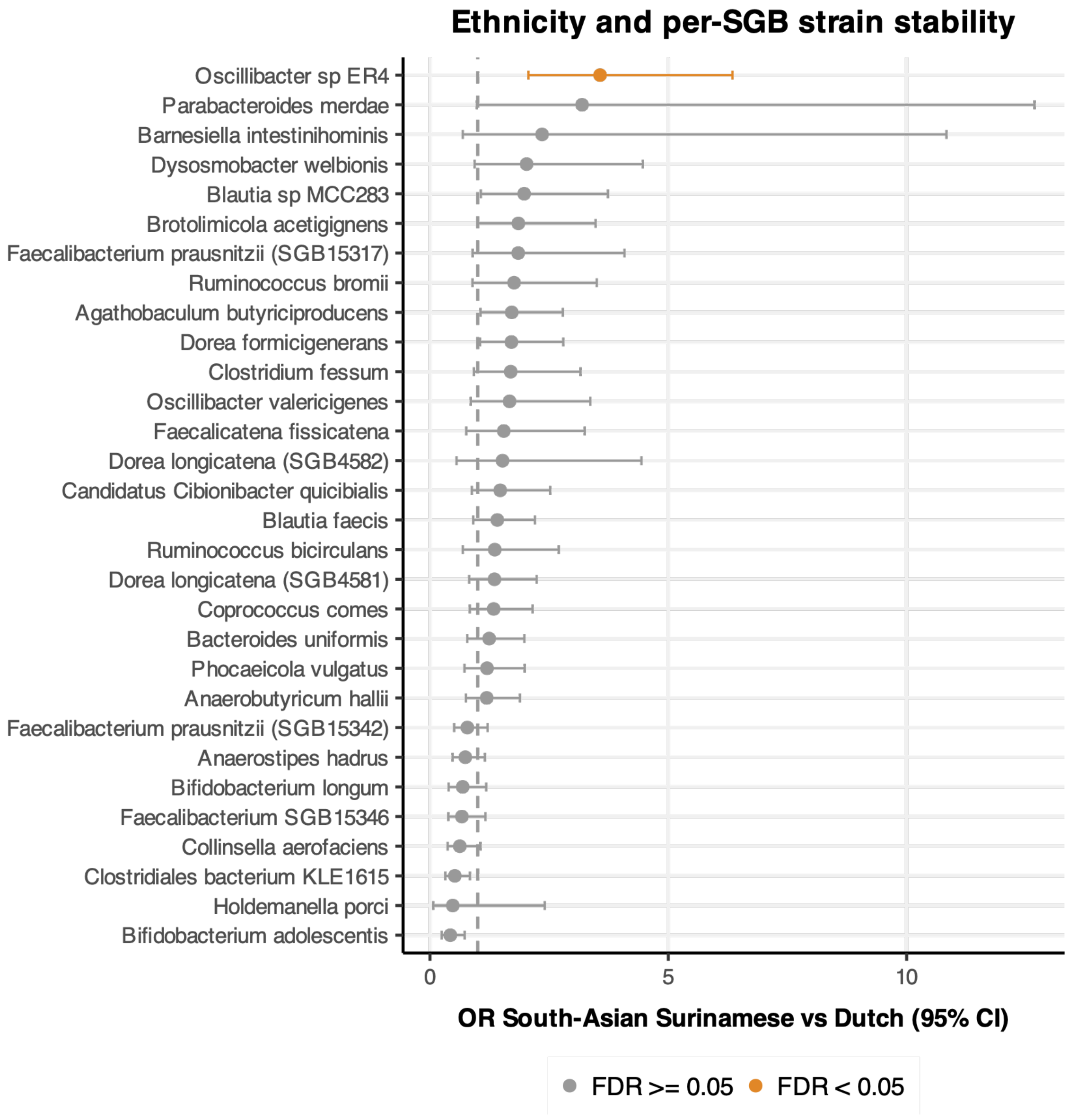


**Supplementary Figure 15: Ethnicity-associated differences in longitudinal strain retention, per species genome bin (SGB).** Forest plot showing odds ratios (OR) with 95% confidence intervals for strain retention in South-Asian Surinamese participants compared to Dutch participants, estimated separately for each SGB. Only 58 SGBs with at minimum N=50 per ethnic group were included in these analyses; this plot shows the top 30. For each SGB, a logistic regression model was fitted with ethnicity as the primary predictor, adjusted for age and follow-up time. P-values were corrected for multiple testing using the Benjamini–Hochberg procedure.

### Supplementary Figure 16: HUMAnN pathway abundance changes over time between ethnic groups

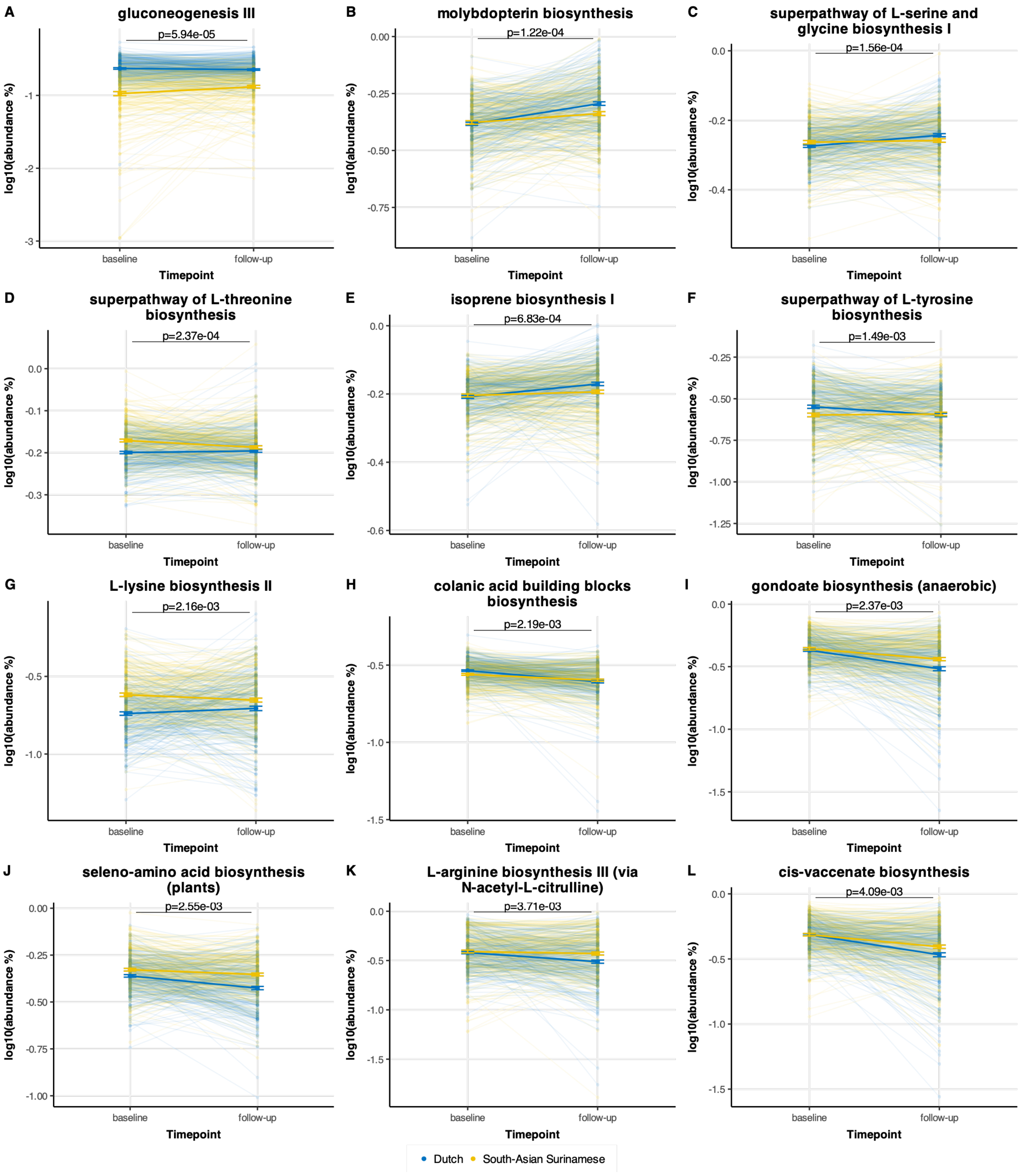

**Supplementary Figure 16: Longitudinal changes in HUMAnN pathway abundance differ between ethnic groups.** Each panel shows $\log_{10}$-transformed relative abundance (%) of pathways at baseline and follow-up for Dutch participants (blue) and South-Asian Surinamese (SAS) participants (yellow). Thin lines connect repeated measurements from the same individual. Thick lines and filled circles represent group means; error bars indicate standard error of the mean. Panels are ordered by strength of the ethnicity × timepoint interaction effect and include only pathways with a significant interaction term in a linear mixed model (LMM; FDR < 0.05) adjusted for follow-up time, with random intercepts for individual participants. Interaction p-values are displayed above each panel.

### Supplementary Figure 17: Ethnic differences in specific pathways and CAZyme families

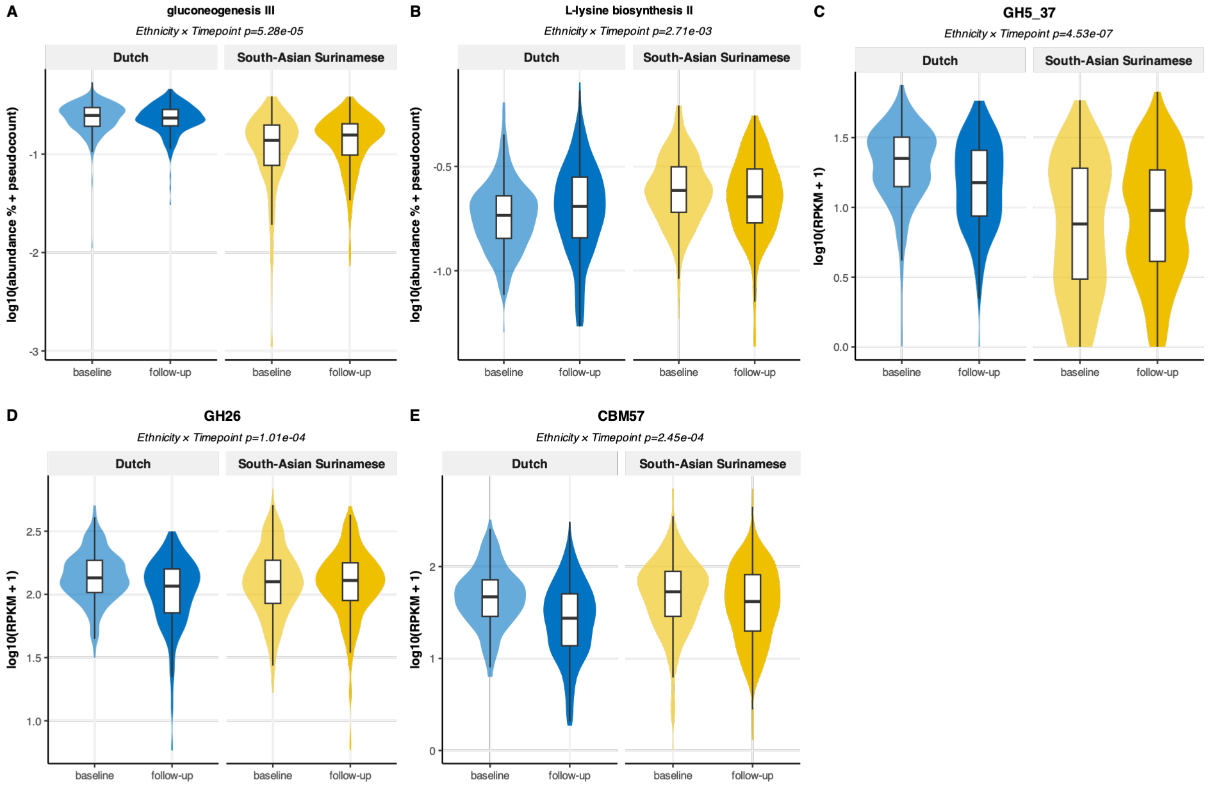


**Supplementary Figure 17: Ethnic differences in specific HUMAnN pathways and CAZyme families.** (A-B) Violin–boxplot distributions of Gluconeogenesis III (A) and L-lysine biosynthesis II (B) pathway abundance (log10-transformed relative abundance) at baseline and follow-up, stratified by ethnicity. (C-D) Violin–boxplot distributions of GH5_37 (C), GH26 (D) and CBM57 log10-transformed abundance (RPKM) at baseline and follow-up, stratified by ethnicity. GH5_37 is a cellulase subfamily of glycoside hydrolase family 5; GH26 is an endo-beta-1,4-mannanase family; CBM57 is a malectin-like carbohydrate-binding module. These families showed the highest significance in the linear mixed model (group*time).

### Supplementary Figure 18: Differentially changing CAZyme families

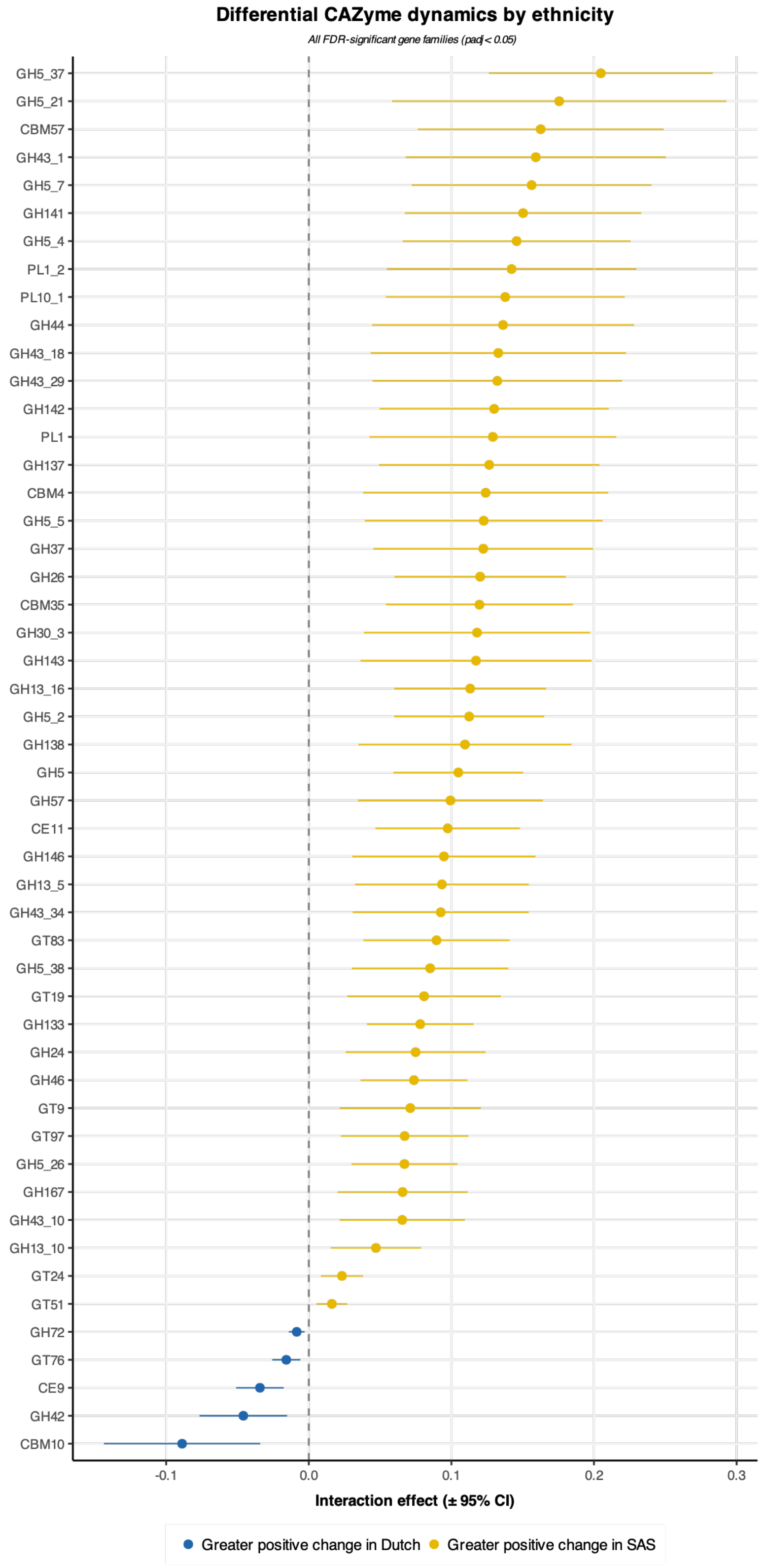

**Supplementary Figure 18: Differentially changing CAZyme families**. Forest plot of CAZyme gene families with a significant ethnicity × timepoint interaction (LMM, FDR < 0.05).

### Supplementary Figure 19: Correlations between baseline CAZyme ratios and clinical variables

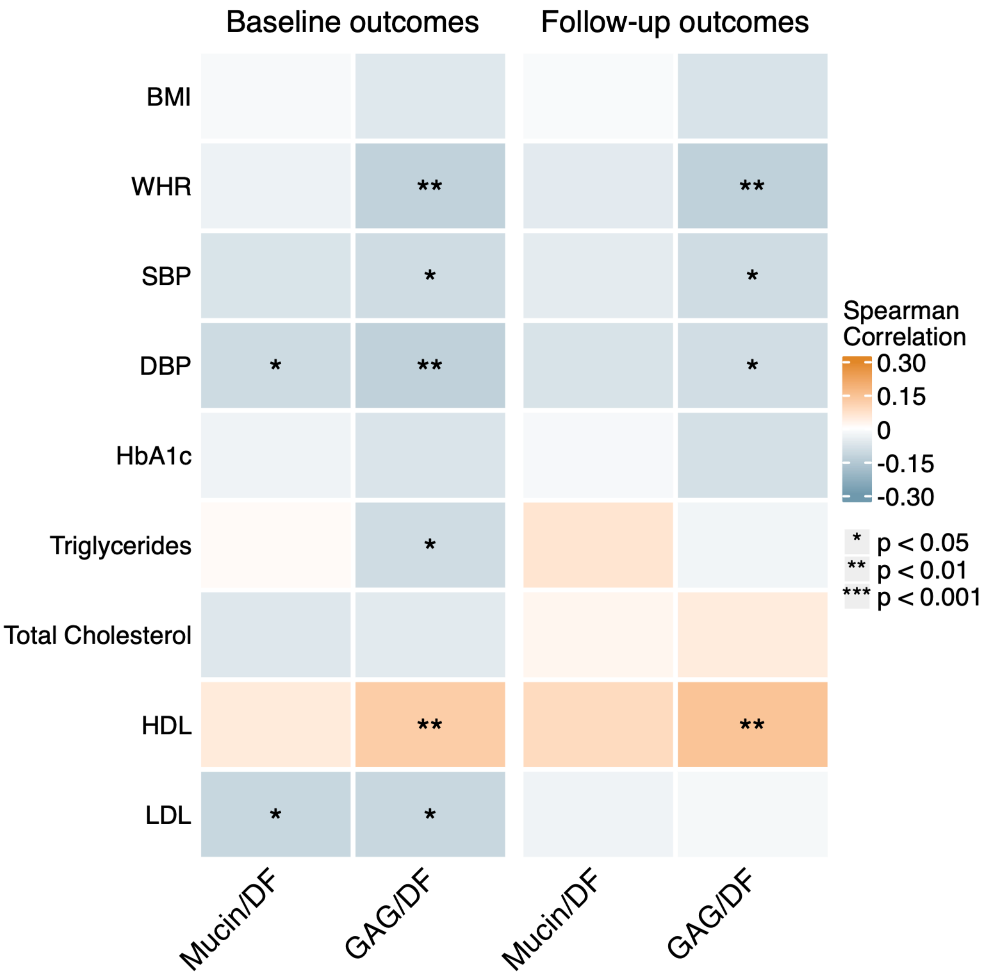


**Supplementary Figure 19: Spearman correlations CAZyme ratios and cardiometabolic parameters at baseline and follow-up.** Heatmaps show Spearman correlation coefficients (ρ) between the log10-transformed Mucin/DF and GAG/DF ratios and cardiometabolic outcomes at baseline (left) and follow-up (right). These are nominal p-values; none of these correlations was significant after multiple testing correction. BMI, body mass index; WHR, waist-to-hip ratio; SBP/DBP, systolic/diastolic blood pressure; HbA1c, glycated haemoglobin; Trig, triglycerides; TC, total cholesterol; HDL/LDL, high-/low-density lipoprotein cholesterol; DF, dietary fibre degraders; GAG, glycosaminoglycan degraders.

### Supplementary Figure 20: Differential change in ARGs

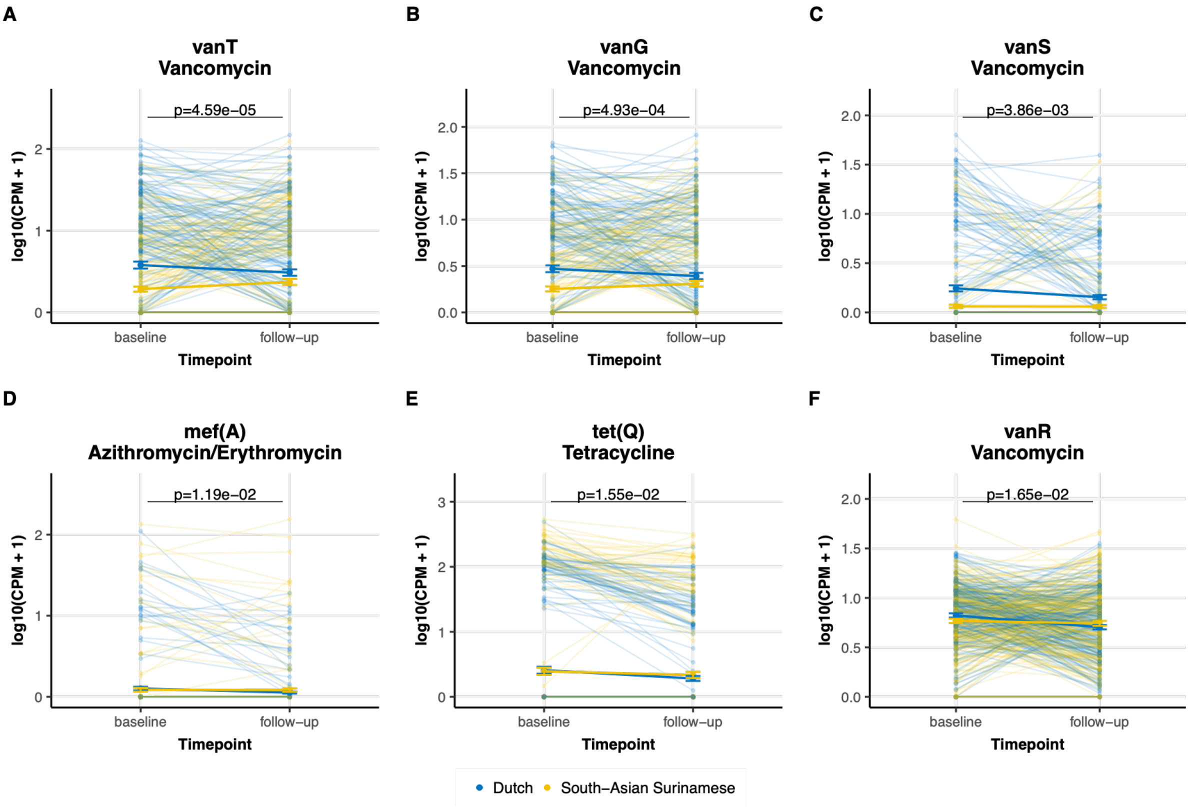


**Supplementary Figure 20: Longitudinal change in antibiotic resistance gene (ARG) abundance by ethnicity.** Spaghetti plots showing individual-level and group-mean trajectories from baseline to follow-up for individual ARG genes with a significant ethnicity × timepoint interaction (A-F). Thin semi-transparent lines represent individual participant trajectories; thick lines connect group means ± standard error at each timepoint. ARG abundance is expressed as log10(CPM + 1) for individual genes. Only genes with a prevalence >5% across all samples were included. Interaction p-values were derived from linear mixed models (LMM) with ethnicity, timepoint, and their interaction as fixed effects, and participant ID as a random intercept. Only *vanT* and *vanG* were significant after multiple testing adjustment.
